# Baby makes three: maternal, paternal, and zygotic genetic effects shape larval phenotypic evolution

**DOI:** 10.1101/2020.12.10.419838

**Authors:** Christina Zakas, Matthew V. Rockman

## Abstract

The evolutionary potential of a population is shaped by the genetic architecture of its life-history traits. Early-life phenotypes are influenced by both maternal and offspring genotype, and efforts to understand life-history evolution therefore require consideration of the interactions between these separate but correlated genomes. We used a four-generation experimental pedigree to estimate the genetic architecture of early-life phenotypes in a species with dramatic variation in larval size and morphology. In the polychaete annelid *Streblospio benedicti*, females make either many small eggs that develop into complex larvae that feed in the plankton or few large eggs that develop into benthic juveniles without having to feed as larvae. By isolating the contributions of maternal, paternal, and zygotic genotype to larval traits, we determined that larval anatomical structures are governed by the offspring genotype at a small number of large-effect loci. Larval size is not shaped by the larva’s own genotype but instead depends on loci that act in the mother, and at two genomic locations, by loci that act in the father. The overall phenotype of each larva thus depends on three separate genomes, and a population’s response to selection on larval traits will reflect the interactions among them.

## INTRODUCTION

Trade-offs between the number and quality of offspring sit at the heart of life-history theory, dictated by finite maternal resources that can be allocated in different ways. Mothers can produce small numbers of large, well-provisioned offspring, or large numbers of small, poorly-provisioned offspring. Ecological factors govern the optimal position of a population along this trade-off front (Vance 1973; Smith and Fretwell 1974; Stearns 1992; Moran and Emlet 2001; Roff 2002).

Efforts to connect theory to the details of real organisms have identified two key areas that theory tends to neglect. First, comparative biology has shown that quantitative differences in size are often associated with dramatic, qualitative differences in form (Thorson 1950; Morgan 1995). Large offspring often develop directly into a miniature version of the adult, while small offspring often develop into complex larval forms, adapted to different conditions than those experienced by their parents. A famous example is the *Heliocidaris* sea urchins (Raff 1996). *H. tuberculata* makes many small eggs that develop into bilaterally-symmetrical planktonic larvae, feeding in the plankton until undergoing a radical metamorphosis to produce the pentaradial adult. Females of the closely related species *H. erythrogramma*, nearly indistinguishable as adults, produce smaller numbers of large eggs, each developing directly into pentaradial juveniles, bypassing the complex anatomy of the feeding larva. Models of evolution along the size-number front should therefore address the concomitant changes in the organismal form of the offspring (Sinervo and McEdward 1988; Wray 1996; Gibson and Gibson 2004; Smith et al. 2007).

Second, the ability of a population to evolve larval size and form, or to remain locally adapted in the face of gene flow, depends on the genetic architecture of variation in the underlying traits (McEdward 2000). Classical models of marine larval life-history suggest that only the extreme strategies — offspring large enough for direct development or else as small as possible — can be stable optima, in which case highly polygenic architectures will trap populations on their local fitness peak (Vance 1973). In such cases a switch-like architecture should be favored, and the underlying genes should congeal into a single-locus supergene (Yeaman 2013). If other fitness landscapes apply, then polygenic architectures may be favored, allowing populations to fine-tune their positions along the size-number and larval form axes.

Critically, the genetic architecture of larval life-history phenotypes has a very particular idiosyncrasy: it involves genomes of both the mother and the offspring, and consequently the many weirdnesses of maternal-zygotic genetics enter the picture (Wade 1998; Wolf 2000). When fitness depends on specific combinations of maternal and zygotic alleles, as it does during embryogenesis and morphogenesis of complex larval traits, frequency-dependence plays an important evolutionary role, as maternal genetic background represents the environment in which zygotic genes act their roles. Linkage and mating system further shape the interaction of maternal and zygotic genetic effects (Drown and Wade 2014). Paternal genetic effects, though rarely considered in evolutionary genetics, could play similar roles. Models of life-history evolution must consider the number and effect-size of the causal genes, and their patterns of linkage and modes of action.

To understand how genetic architecture shapes the evolution of larval life-history traits, in the context of qualitative transitions in larval form, we have developed an experimental model in the estuarine polychaete *Streblospio benedicti*. These animals, which are native to the east coast of North America and live in intertidal and subtidal mud as adults, have two different ways of developing (Levin 1984). Some *S. benedicti* females produce small (~100 μm diameter) eggs that develop into characteristic planktonic larvae, which feed on unicellular algae for a period of days or weeks before undergoing a metamorphosis and settling into the benthic mud. Other females produce a much smaller number of much larger eggs (~200 μm diameter, 8-fold larger in volume). These eggs develop into large larvae that do not need to feed and can immediately metamorphose and settle. In both modes of development, embryogenesis and larval morphogenesis occur in brood pouches on the female’s back, and offspring are released into the water as fully formed planktonic larvae that have to feed (planktotrophs) or metamorphosis-competent larvae that do not require planktonic food, provisioned by yolk (lecithotrophs).

The two classes of *S. benedicti* larvae differ substantially in form (Gibson et al. 2010; Pernet and McHugh 2010), with the small planktotrophic larvae bearing two kinds of larva-specific structures. These are the long, barbed larval chaetae, which they flare when perturbed, putatively as a defense (Pennington and Chia 1984), and a set of four tail-tip appendages called anal cirri, each occupied by a large and morphologically distinctive cell of unknown function, known as a bacillary cell (Gibson et al. 2010). Both the larval chaetae and the anal cirri are lost at metamorphosis. The large lecithotrophic larvae, in contrast, never develop the larval chaetae or anal cirri. Instead, they accelerate development of juvenile morphology, including additional segments and juvenile structures. Development of larval chaetae has been lost or suppressed in many independent lineages that have evolved lecithotrophy, suggesting that these structures are disadvantageous for larvae with short planktonic periods (Lewis 1998; Gibson and Gibson 2004).

A key feature of *S. benedicti* is that larval mode is insensitive to environmental cues or manipulations (Levin and Creed 1986; Levin and Bridges 1994) and instead exhibits a highly heritable basis (Levin et al. 1991; Zakas and Rockman 2014). Previously, we identified quantitative trait loci (QTL) that affect larval traits by measuring genotypes and phenotypes in an F_2_-like experimental panel derived from a cross of a lecithotroph and a planktotroph (Zakas et al. 2018). Because the animals shared F1 mothers, heterozygous at every locus that differs between lecithotrophs and planktotrophs, the experimental panel was blind to maternal-effect genetic variation. Here, we study the next generation of the experiment, where both larval genotype and maternal genotype are segregating. The design of the cross allows us to explicitly test for loci that act in the mother, the father, or the larva.

## METHODS

To determine the empirical genetic architecture of larval traits, we used a four-generation pedigree of *S. benedicti,* as described previously (Zakas et al. 2018). Briefly, we crossed a planktotrophic female from Bayonne, NJ, and a lecithotrophic male from Long Beach, CA. The Long Beach population is the result of a recent introduction from the East Coast (Schulze et al. 2000). We then crossed a single F_1_ male with each of four F_1_ females to generate four F_2_ families, each related to the others as half-sibs. We refer to this generation as a whole as G_2_ rather than F_2_ to mark the fact that animals in this generation are not all full siblings. We then intercrossed G_2_ individuals to generate G_3_ families. To test for maternal effects on larval morphology, we here add phenotype data for 3,954 larvae from the G_3_ generation. These represent 115 families (mean and median 34 full-sib larvae per family), made from crosses among 115 G_2_ females and 53 G_2_ males. Individual G_2_ males sired an average of 2 G_3_ families (number of families per male: 1 (n=20), 2 (n=12), 3 (n=13), 4 (n=8)). The G_2_s used as parents are a subset of the 183 females and 58 males described in our earlier study (Zakas et al. 2018).

Previously, we genotyped the G_2_ animals at 702 nuclear markers (Zakas et al. 2018). We use those genotypes here, filtered down to 275 with unique segregation patterns. In addition, we imputed missing data at several markers on the X chromosome (4% of 6,266 X-linked genotype calls), using the sim.geno function in rqtl (Broman et al. 2003).

To phenotype G_3_ larvae, we fixed newly released broods in 4% formaldehyde overnight, washed them with PBS, and mounted them on microscope slides in stable mountant. We imaged each larva with a Zeiss Axio Imager M_2_ with 20x objective and DIC optics and we used ImageJ to score each larva for size (area), number of defensive chaetae, length of the longest chaeta, and presence or absence of anal cirri.

To map quantitative-trait loci (QTL) underlying trait variation in the G_3_s to genotypes from the G_2_s, we employed mixed-effect models that account for family structure and accommodate parental genetic effects. We performed a series of genome-wide linkage scans to detect QTL. We subsequently tested the significant QTL for their transmission mode (e.g., zygotic vs. maternal effect), dominance, and epistasis.

To search the genome for QTL, we step through each genetic marker position, modeling the G_3_ phenotype vector *y* as

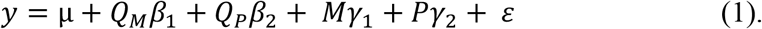

Here μ is the global mean vector, the *Q*_M_ vector holds the count of lecithotroph alleles at the marker in the G_2_ mother (0 to 2), the Q_P_vector holds the count of lecithotroph alleles at the marker in the G_2_ father, and betas are the additive effects of maternal and paternal alleles at the marker. We model the family structure with random effects for the mother and father. Incidence matrices *M* and *P* connect each larva with the appropriate entry in the γs, vectors of random effects associated with each parent. Here the random effect of mother accounts for diverse, sources of similarity among full sibs; these include background maternal and zygotic genetic effects, maternal environmental effects, and effects due to early development in a common environment. The paternal effect accounts for similarity among half sibs, which should only be present if there are zygotic or paternal genetic effects. ε is the residual error.

We compare the fit of this model to a no-QTL model,

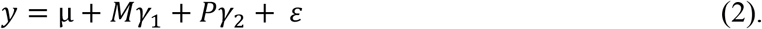

This model is nested within the previous model and has two fewer parameters. We use twice the difference between the models in log likelihood as a test statistic. This is the canonical likelihood ratio test statistic, asymptotically distributed as a chi-square under the null hypothesis. For genome scans, we use permutations, described below, to generate genome-wide null statistics.

For purely zygotic-effect inheritance, we can achieve higher power by using a single beta, as the effects of maternal and parental alleles should be identical. Additive zygotic effects can therefore be modeled as the effect of the sum of alleles across the two parents of each G_3_ larva,

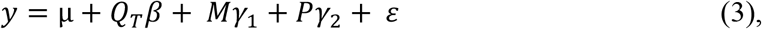

where the *T* subscript indicates total allele count (0 to 4). We again test this model against (2), the no-QTL null model.

To directly test for parental effects at QTL detected by the tests above, we compare models 1 and 3 to one another, testing whether the model fit is significantly improved by allowing allelic effects to differ according to the parental sex. Rejection of model 3 is evidence of maternal effects, paternal effects, both kinds of effects, or a combination of parental and zygotic effects. To test a purely maternal-effect model at such loci, we compare model 1 to a reduced model with no effect of paternal genotype at the locus:

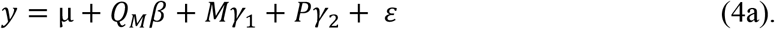

Rejection of model 4 indicates that the paternal genotype at the marker matters, rejecting a purely maternal-effect model. Similarly, a paternal-genotype-only model tests whether the maternal genotype matters:

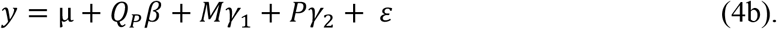

To define thresholds for genome-wide significance, we used structured permutations (Churchill and Doerge 2008). We shuffled G_2_ male and G_2_ female genotypes independently, preserving the underlying full- and half-sib relatedness among G_3_s. The permutations are also stratified by G_2_ family (i.e., genotypes are shuffled only among G_2_s that share a single F_1_ mother), as in (Zakas et al. 2018). Any effects due to structure within the G_2_ generation are preserved in the permutations and thereby controlled.

When the initial genome scan identified a significant locus (i.e., rejection of model 2), we then incorporated the genotype at that locus into a second genome scan, using the genetic model for the locus identified by comparisons among models 1, 3, and 4. This forward search continued until no additional loci exceeded a genome-wide permutation-based threshold of p = 0.05. For each round of search, the permutation models incorporated the loci accepted in the previous round as a fixed-effect covariate (Doerge and Churchill 1996). For example, the no-QTL null (model 2) becomes the single-QTL null *y* = μ + *Q*_*T*1_*β*_*QT*1_ + *Mγ*_1_ + *Pγ*_2_ + *ε* when an additive-effect QTL is incorporated into the model with genotype vector *Q*_*T*1_ and effect *β*_*QT*1_ and genome-wide linkage scans (models 1 and 3) then test for additional *Qβ* terms. Note that we use permutations to define genome-wide significance for linkage scans (step 1 in the testing framework), and for subsequent tests involving the loci detected in step 1 use uncorrected p = 0.05 threshold from likelihood ratio tests.

Testing framework:
| Models | Null | Rejection implies |
| --- | --- | --- |
| Step 1. Genome-wide linkage scans |  |  |
| (1) vs (2): | No QTL | Presence of a QTL, parental or zygotic or both |
| (3) vs (2) | No QTL | Presence of an additive-effect QTL |
| Genome-wide significance determined by analysis of permuted datasets. |  |  |
| Step 2. Tests for parental effects at significant QTL |  |  |
| (1) vs (3) | Additive-effect QTL | Parental-effect QTL |
| Significance determined by likelihood ratio test with chi-square null distribution. |  |  |
| Step 3. Tests for multiple parental/zygotic effects at significant parental-effect loci |  |  |
| (1) vs (4a) | Maternal-effect QTL | More than just maternal-effect QTL |
| (1) vs (4b) | Paternal-effect QTL | More than just paternal-effect QTL |
| Significance determined by likelihood ratio test with chi-square null distribution. |  |  |
| Step 4. Add significant QTL to the models and repeat steps 1 to 3 until no additional loci are detected. |  |  |

The analyses described above were applied to autosomal markers. The models are not precisely suited to the transmission genetics of the X chromosome, because of G_2_ male hemizygosity, unknown sexes among the G_3_ larvae, and unbalanced G_2_ genotypes due to the direction of the pedigree-founding cross. As the sole F1 male in the pedigree carried the X chromosome from its planktotroph mother, all G_2_s females carry at least one planktotroph X chromosome. Denoting the lecithotroph allele L and planktotroph allele P, G_2_ females can never be LL homozygotes at X-linked loci. We can nevertheless test for linkage to X-chromosome markers by comparing the full model to the no-QTL model (1 vs 2), and we can test for parental effects by comparing the full model to the one-parent-only model (1 vs 4).

In addition, we consider a modified model 3 (zygotic additive model) with X-linked allelic affects weighted according to the sex and cross. For X-linked markers in our G_2_ population, there are four types of crosses (Supplementary Figure 1). Under the assumption that males hemizygous for an allele have the same genotypic value as females homozygous for that allele, we can, calculate the expected offspring genotype for each cross in terms of expected lecithotrophic allele count. As shown in Supplementary figure 1, these are 5/8, 3/8, 1/4, and 0 for the four possible classes of G_3_ broods.

For all analysis of X-linked markers, we defined X-specific significance thresholds via permutation. Note that our previous analysis of G_2_ phenotypes found no evidence for sex-linkage or for differences between sexes in any larval phenotypes (Zakas et al. 2018), suggesting an absence of X-linked zygotic-effect loci. The present analysis allows for tests of X-linked parental-effect loci and, weakly, for zygotic effects that manifest in LL homozygous females, which are present among the G_3_s as 1/4 of the offspring of X^L^X^P^ × X^L^Y crosses. Note that all males in the pedigree derive their Y chromosome from the single lecithotrophic male founder, and so Y-linked effects cannot contribute to phenotypic variation in the cross.

Traits are larval size (area occupied by the larva on a microscope slide), number of larval chaetae (on one side of the larva), length of the longest larval chaeta, and presence or absence of anal cirri. Number of chaetae is a mixture of a point mass at zero (24.8% of larvae) and distribution with a mode at 4 chaetae and a long right tail (i.e., overdispersed relative to a Poisson distribution). We therefore modeled number of chaetae as two separate traits: presence/absence of chaetae as a binary trait, fit with a generalized linear mixed model approach (i.e., mixed-effect logistic regression), and Box-Cox-transformed chaetae number conditional on their presence, using the standard approach. These two approaches to the data yielded extremely similar results, and so we proceeded by combining the two models (i.e., summing the log likelihoods). The result is a mixed-model analog of the two-part model of (Broman 2003). The larvae with no chaetae are missing data for the chaetal length phenotype, which is normally distributed conditional on their exclusion. Finally, presence of anal cirri is a binary trait, modeled by mixed-effect logistic regression.

To test for directional polygenic effects, under the assumption that loci that differ between lecithotrophs and plankotrophs will tend to affect larval traits in a predictable direction, we used expected autosomal ancestry (fraction lecithotroph alleles) as a predictor in a mixed effect model that also includes the genome-wide significant loci. We consider both biparental ancestry (i.e., zygotic model) and uniparental ancestry (maternal-effect or paternal-effect model). We estimate ancestry as the fraction of lecithotroph alleles at autosomal markers, excluding the chromosomes that carry genome-wide significant loci for the trait (parent-specifically for parental-effect loci).

For loci inferred to act via parental genetic effect, we tested for dominance effects by incorporating a term for parental heterozygosity. For loci inferred to act via zygotic effects, we cannot use this direct approach because we lack individual-level genotypes for the G_3_ larvae. Instead, we tested for dominance by asking whether the fit of the additive zygotic model was significantly improved by addition of a coefficient for the expected heterozygosity of the larva (e.g., 1 for offspring of a PPxLL cross, 0.5 for PLxPL or PPxPL, 0 for PPxPP, etc).

To improve mapping resolution beyond that achieved with G_3_ phenotypes, we integrated G_2_ and G_3_ phenotype data into a single genome scan for zygotic effects by summing the G_2_ and G_3_ log likelihoods at each marker under an additive zygotic model and testing against the summed log likelihoods of the relevant no-QTL models. We also performed this genome scan after incorporating each of the established QTLs into the model, with dominance where appropriate, as covariates. Likelihoods for the G_2_s were estimated under simple fixed effect models. Note that this approach allows for the allelic effects of a marker to differ between the generations. Significance thresholds were estimated by structured permutation of the male and female G_2_ genotypes, preserving relatedness between each G_2_ animal and its progeny.

We estimated narrow-sense heritability (*h*^2^) using mixed-effect parent-offspring regressions for traits with continuous distributions (i.e., excluding presence of cirri). For additive zygotic-effect inheritance, the slope of the regression of offspring phenotypes on the average phenotype of the parents, accounting for family structure with random effects, is an estimator of *h*^2^. We tested whether the heritabilities were different from zero by comparing the regressions to a null model with no effect of parental phenotype, and we tested for differences between maternal and paternal estimates by including the two parental phenotypes as separate fixed effects. Note that our heritability estimates apply only to transmission of traits from G_2_ to G_3_ in this specific cross. To estimate whether the detected QTLs account for the estimated heritability, we incorporated the QTLs as fixed effects in the regressions and compared these to no-QTL regressions. We tested whether heritabilities were different from zero using likelihood ratio tests.

Analyses were performed in R (R Core Team 2015), version 4.0.2, using functions from *rqtl* (Broman et al. 2003) and *lme4* (Bates et al. 2015). Phenotypes, genotypes, and mapping functions are provided in Supplementary File 1.

## RESULTS

We studied the great-grandchildren from a cross of a planktotrophic female and a lecithotrophic male. The grandchildren (G_2_ animals) are genetically heterogeneous and the phenotypes of their G_3_ offspring therefore permit us to detect loci acting both parentally in the G_2_s and zygotically in the G_3_s.

The G_3_ larvae exhibited a broad range of phenotypes, including many forms not encountered in natural populations, such as large larvae with chaetae and small larvae without (Figure 2).

**Figure 1:**
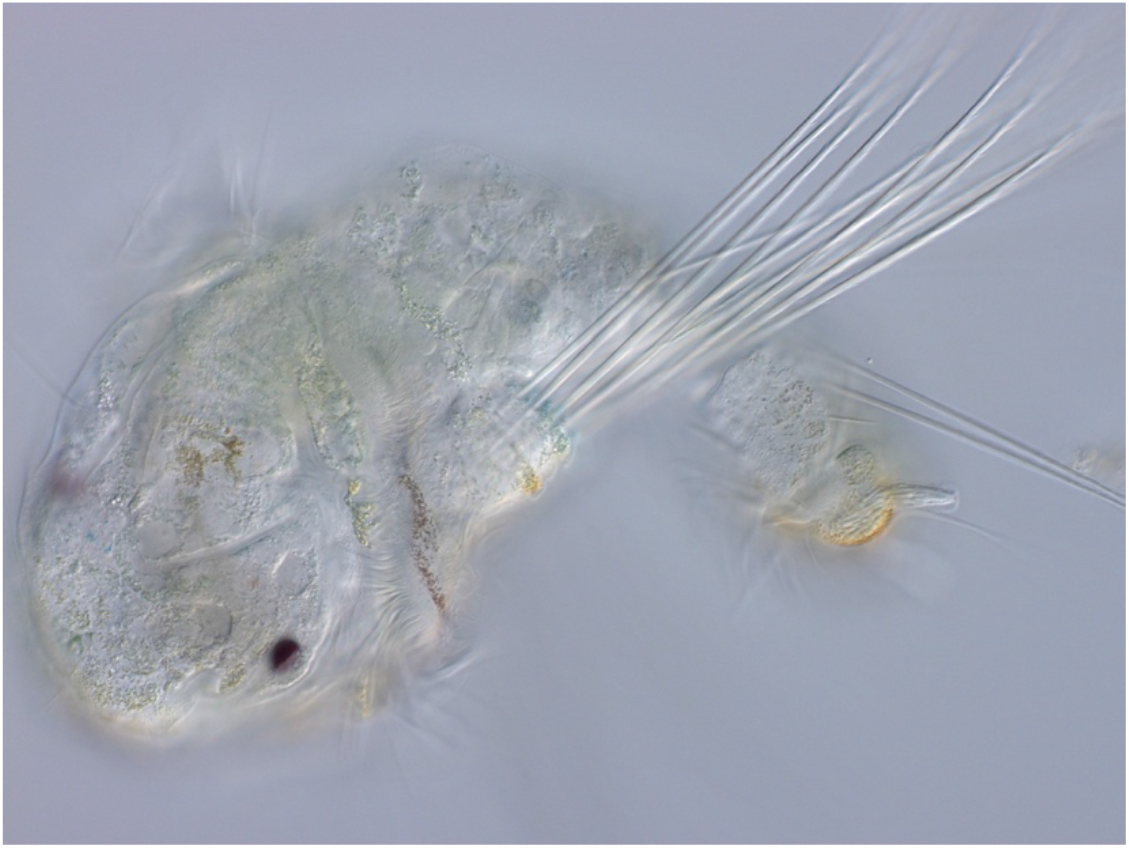
A freshly released planktotrophic larva, anterior to the left. The larva-specific chaetae are those that emerge from the lateral chaetal pouches; the left pouch and its 10 chaetae are visible here. Other chaetae visible in the image are segmental chaetae, which are present in both planktotrophs and lecithotrophs and are retained in juveniles and adults. At the posterior end of the animal (right), one anal cirrus is in focus. The cirrus is occupied by a large bacillary cell, so-called because of the rod-shaped (bacillus-like) inclusions that fill it.

**Figure 2.**
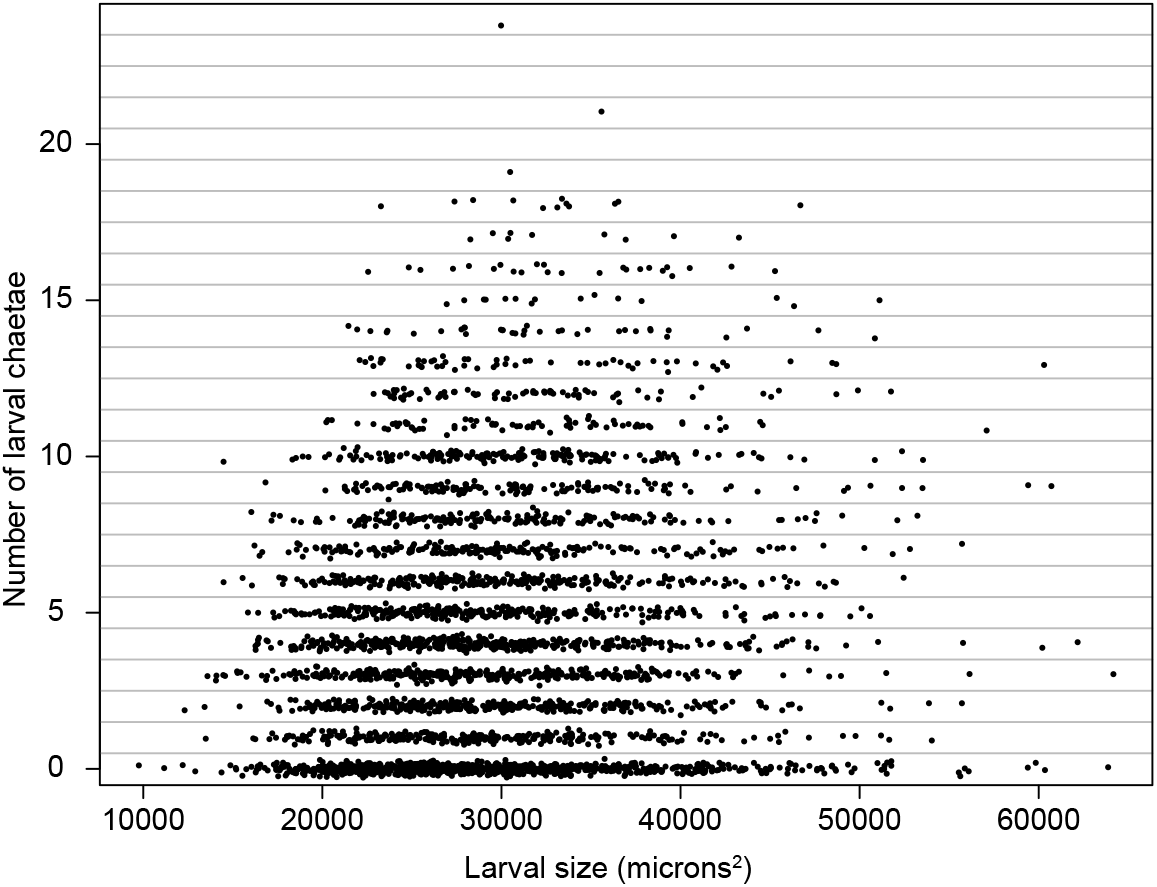
The joint distribution of G_3_ larval chaetae number and larval size. To make the points visible we have added variation in their y-axis positions within each chaetae-number bin. The G_3_s include forms not seen in nature, such as large larvae with chaetae and small larvae without.

Formally, lecithotrophy and planktotrophy refer to larval diets, and as we fixed larvae at release from their mothers’ brood pouches, we did not assay diet and cannot assign diet-based labels. Instead, we measured larval size at release and three larval traits: the number of larva-specific serrated chaetae, the length of the chaetae, and the presence or absence of anal cirri.

The number of larval chaetae has a bimodal distribution: a quarter of the G_3_ larvae have zero chaetae, and the remainder come from a distribution with a mode of four chaetae (Figure 3). A simple interpretation of the distribution is that there is a single normally-distributed underlying genetic liability centered on four chaetae, and because negative counts are impossible, the section of the liability distribution that falls below zero accumulates at zero. Consistent with this view, our separate analyses of the presence or absence of chaetae and of the number of chaetae among larvae with chaetae pointed to nearly identical genetic architectures. We therefore mapped loci affecting the presence/absence and number of chaetae as a single trait, summing the log-likelihoods from a logistic regression of the former and a linear regression of the latter (Box-Cox transformed).

**Figure 3.**
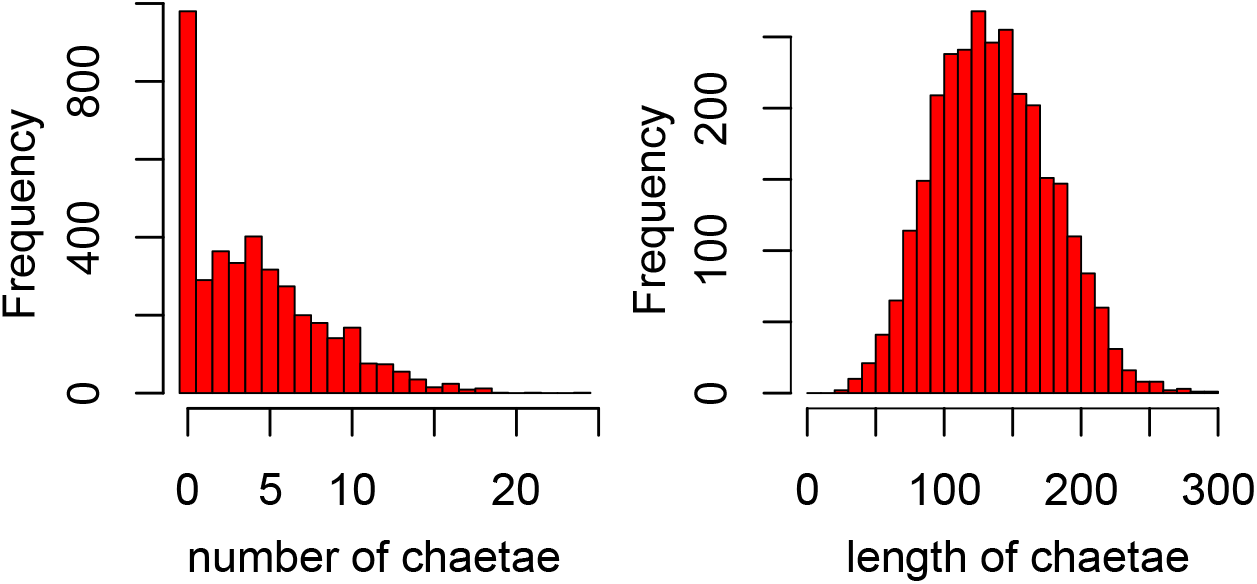
Distributions of the number of larval chaetae (left) and their length, among the larvae that have such chaetae (right).

A linkage scan with a forward-search strategy revealed major-effect QTL for each trait (Supplementary Table 1). We identified two loci that affect chaetae number, two that affect chaetae length, and one that affects presence or absence of the anal cirri (Table 1). At none of these loci did we detect any evidence for maternal effects. One locus, on chromosome 3, affects both length and number of the chaetae. The chromosome 3 locus exhibits significant dominance in its effect on the number of chaetae (the number-reducing allele is dominant), precisely recapitulating the zygotic effects detected previously in the G_2_ generation (Zakas 2018). We detected dominance at both loci affecting chaetae number, but this dominance only manifests in the part of the model describing the presence or absence of chaetae and not their number when present. We interpret this as a reflection of the threshold nature of the absence of chaetae; the loci appear to act additively on the number of chaetae, but this translates into a nonlinear relationship with chaetae presence or absence, because the genetic value of the homozygous lecithotroph genotype is fewer than zero chaetae. We found nominal evidence for epistasis between the two loci affecting chaetae number, but only for the part of the continuous part of the model. Families with lecithotroph alleles at both QTL have more chaetae than would be expected from the additive effects of the two loci. We found no evidence for X-linkage or directional polygenic effects on any of these traits.

**TABLE 1.**
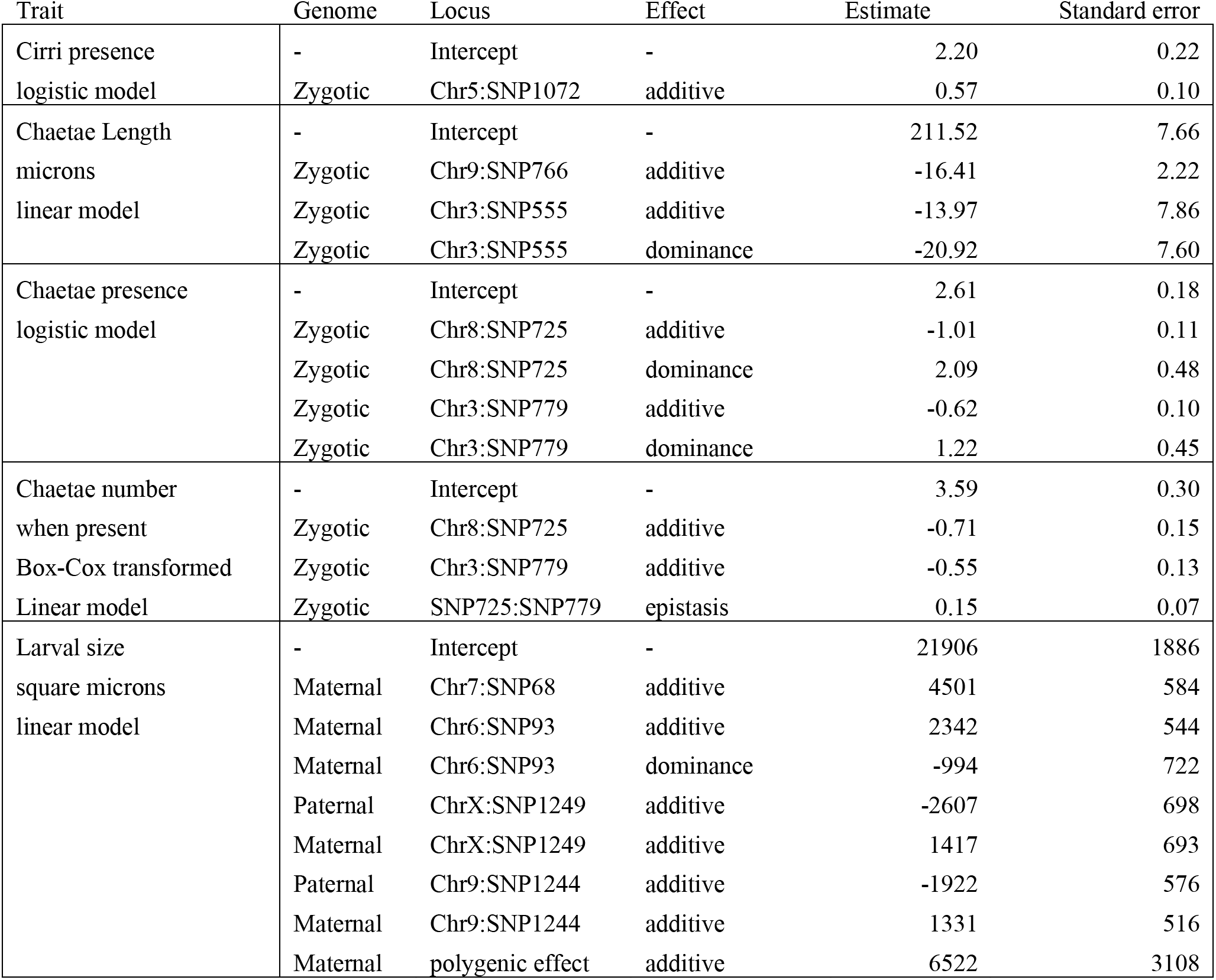
QTL effects. QTL identified from forward search of G_3_ data. Additive effects of SNPs represent the effects of each additional allele from the lecithotrophic founder genome.

| Trait | Genome | Locus | Effect | Estimate | Standard error |
| --- | --- | --- | --- | --- | --- |
| Cirri presence<br>logistic model | - | Intercept | - | 2.20 | 0.22 |
|  | Zygotic | Chr5:SNP1072 | additive | 0.57 | 0.10 |
| Chaetae Length<br>microns<br>linear model | - | Intercept | - | 211.52 | 7.66 |
|  | Zygotic | Chr9:SNP766 | additive | -16.41 | 2.22 |
|  | Zygotic | Chr3:SNP555 | additive | -13.97 | 7.86 |
|  | Zygotic | Chr3:SNP555 | dominance | -20.92 | 7.60 |
| Chaetae presence<br>logistic model | - | Intercept | - | 2.61 | 0.18 |
|  | Zygotic | Chr8:SNP725 | additive | -1.01 | 0.11 |
|  | Zygotic | Chr8:SNP725 | dominance | 2.09 | 0.48 |
|  | Zygotic | Chr3:SNP779 | additive | -0.62 | 0.10 |
|  | Zygotic | Chr3:SNP779 | dominance | 1.22 | 0.45 |
| Chaetae number<br>when present<br>Box-Cox transformed<br>Linear model | - | Intercept | - | 3.59 | 0.30 |
|  | Zygotic | Chr8:SNP725 | additive | -0.71 | 0.15 |
|  | Zygotic | Chr3:SNP779 | additive | -0.55 | 0.13 |
|  | Zygotic | SNP725:SNP779 | epistasis | 0.15 | 0.07 |
| Larval size<br>square microns<br>linear model | - | Intercept | - | 21906 | 1886 |
|  | Maternal | Chr7:SNP68 | additive | 4501 | 584 |
|  | Maternal | Chr6:SNP93 | additive | 2342 | 544 |
|  | Maternal | Chr6:SNP93 | dominance | -994 | 722 |
|  | Paternal | ChrX:SNP1249 | additive | -2607 | 698 |
|  | Maternal | ChrX:SNP1249 | additive | 1417 | 693 |
|  | Paternal | Chr9:SNP1244 | additive | -1922 | 576 |
|  | Maternal | Chr9:SNP1244 | additive | 1331 | 516 |
|  | Maternal | polygenic effect | additive | 6522 | 3108 |

We used mixed-model parent-offspring regression to estimate the heritability of chaetae length and number (Table S2). We estimated *h*^2^ = 0.43 ± 0.10 for chaetae length and *h*^2^ = 0.43 ± 0.09 for chaetae number. After incorporating the genotypes at the mapped QTLs, the residual heritability was not significantly different from zero (0.03 ± 0.09 and 0.02 ± 0.09). These results imply that we have detected the loci responsible for the bulk of the additive genetic basis for these traits.

By combining mapping data from the G_2_ and G_3_ generations, we can achieve higher power to detect and localize zygotic-effect quantitative trait loci. We found that the genetic signals added and background noise largely cancelled, leaving the few significant QTL defined more clearly (Figure 4, Table S3). In addition, the joint analysis improved the localization of some loci; for example, the chromosome 5 locus affecting the presence of anal cirri shifted several centimorgans from its position in the G_3_ linkage analysis. The loci on chromosome 3 affecting chaetae length and number colocalize to a single peak position in the joint analysis. A joint G_2_-G_3_ scan incorporating all the known QTL as covariates did not identify additional loci at the genome-wide significance threshold.

**Figure 4.**
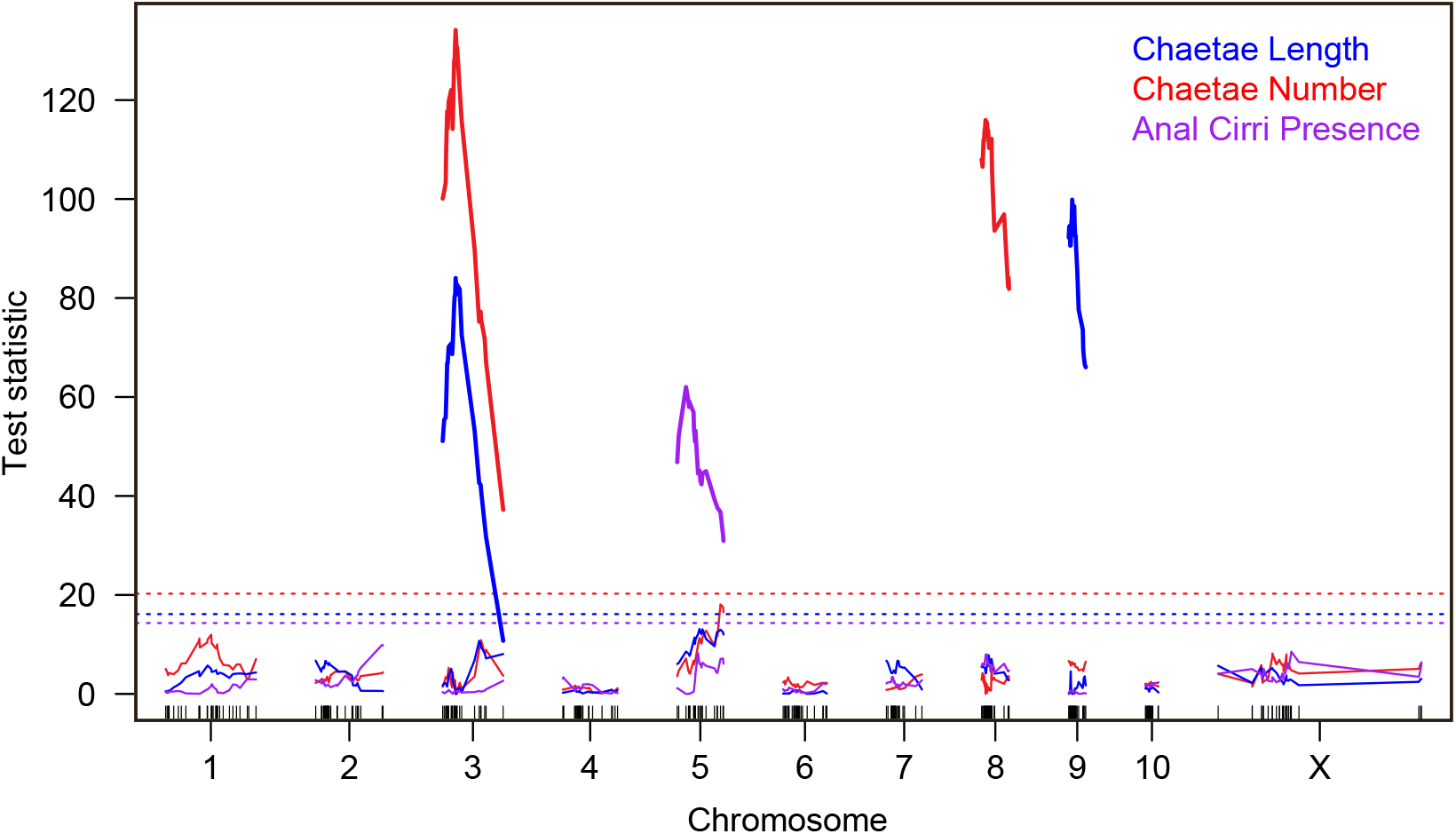
Joint analysis of G_2_ and G_3_ phenotype data identified four loci affecting three larval anatomy traits. Thick lines show the test statistics for the chromosomes with significant QTL identified in successive rounds of forward search. Test statistics for the first round with no significant QTLs are shown as thin lines, below the dashed lines marking the trait-specific p = 0.05 thresholds.

For larval size, we previously reported a maternal-effect QTL analysis based on the brood-means of 183 G_2_ females (Zakas et al. 2018). The 115 broods analyzed here are the subset for which we also have genotype data for the father of the brood. We applied our mixed-model approach to formally test alternative genetic models and detected four significant loci, each with pronounced parental effects (Figures 5 and 6, Table 1, Table S1). The top two loci show exclusively maternal effects. These loci, on chromosomes 6 and 7, coincide with those we identified previously (Zakas et al. 2018), and each lecithotroph allele is associated with an increased larval size. A uniparental maternal-effect model is not significantly different from the full model with different effects from each parent, and a uniparental paternal-effect model is strongly rejected.

**Figure 5.**
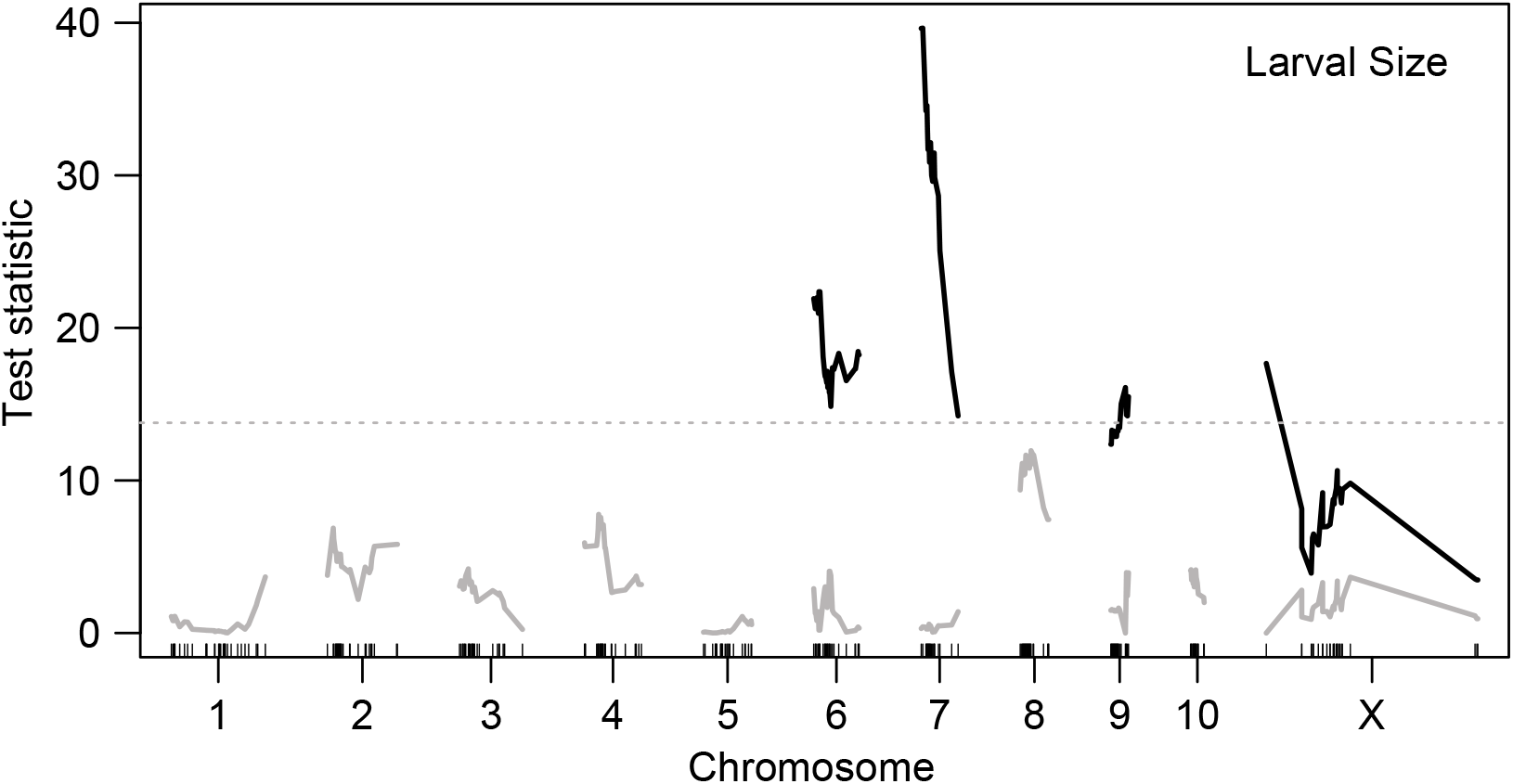
A forward-search strategy identified four genome-wide significant parental-effect loci affecting larval size. Shown in black are the test statistics for the chromosomes with significant QTL identified in four rounds of search with the full model. Test statistics for the fifth round of search are shown in gray, below the dashed line marking the genome-wide p = 0.05 threshold.

**Figure 6.**
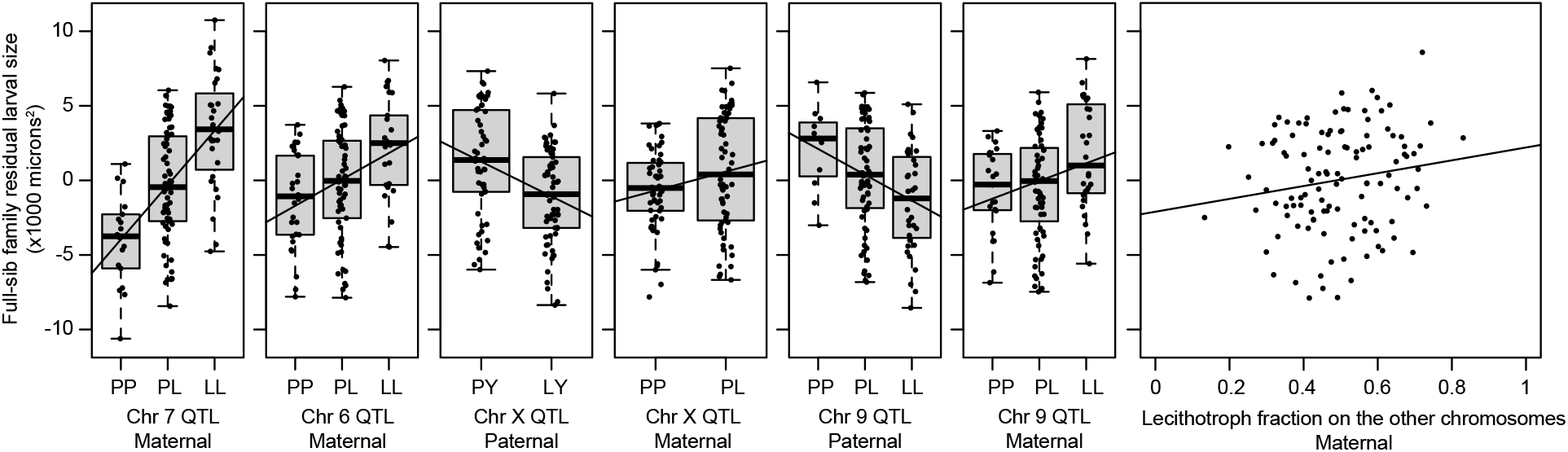
These seven plots show the effects of each parental QTL and of the polygenic background on the chromosomes without significant QTL. Each point represents the Best Linear Unbiased Predictor (BLUP) for one of the 115 full-sib families, accounting for all the other genetic effects as fixed effects in a mixed-effect model. For example, the variation among points in the leftmost panel represent residual variation among full-sib families from a model that includes the six other genetic effects. To aid visualization, we show linear regressions of these BLUPs on lecithotroph gene dosage. The plotted slopes differ slightly from the effects estimated in the full model (Table 1), which appropriately weights each family.

At the other two loci, on chromosomes 9 and X, the data strongly reject an additive zygotic model in favor of the full model with separate allelic effects for each parent, and they strongly reject the maternal-only uniparental model (p = 0.0004 and 0.0010 for the loci on X and 9 respectively), consistent with paternal effects. Both loci also nominally reject the uniparental paternal-effect model (p-values 0.0460 and 0.0196), suggesting maternal effects. The direction of the inferred effects depends on the parent; lecithotroph alleles carried by mothers are associated with larger larvae, and the same alleles carried by fathers are associated with smaller progeny (Figure 6). The latter effects are larger.

Note that for X-linked loci, the test of additive vs. parental effects is complicated by sex-dependent parental genotypes (XX vs XY). However, this complication should not result in a parent-dependent change in the sign of an allelic effect, such as we observe on the X-linked locus affecting larval size.

Although the data are consistent with antagonistic paternal and maternal effects at the loci on chromosomes 9 and X, other explanations are possible in principle. For example, Hager et al. (2008) show that a locus with a maternal effect and a zygotic effect in the opposite direction can give the appearance of a paternal effect. Our G_3_ data do not allow us to simultaneously estimate maternal, paternal, and zygotic effects, but we can test for zygotic effects in a setting that controls for maternal and paternal effects. Specifically, the G_2_ generation varies zygotically but all individuals share heterozygous F1 parents and so parental-effect variation is absent. We found no evidence for direct effects of the loci on X (p = 0.88) or chromosome 9 (p = 0.21) in an analysis of 240 G_2_ larvae (data from Zakas et al. 2018).

The maternal-effect QTL on chromosome 6 exhibited partial dominance with respect to larval size, with heterozygotes smaller than the additive expectation. We also detected a significant directional maternal polygenic effect: setting aside the chromosomes that carry significant maternal-effect QTL (6, 7, 9, and X), the fraction of remaining maternal genome derived from the lecithotrophic founder explains additional variation in larval size (Figure 6). We detected no analogous directional polygenic effect of paternal or zygotic genome. We also found no evidence for statistical interactions among the genome-wide significant loci, either within or between parental genomes. Thus maternal-effect loci with small effect sizes are distributed across some or all of the seven autosomes that lack major-effect QTL.

As expected, mixed-effect parent-offspring regression found no evidence for heritability of larval size between the G_2_ and G_3_ generations (p = 0.19; Table S2). Given purely parental-effect inheritance, phenotypic variation in larval size in the G_2_ generation reflects only environmental effects, as all G_2_s shared F1 parents, each equally heterozygous at all loci that differ between the lecithotroph and planktotroph founders of the pedigree. By contrast, variation in the G_3_ generation reflects both environmental effects and genetic variation among the G_2_s.

## DISCUSSION

Our analysis of almost four thousand G_3_ larvae provides strong evidence that the genetic basis of size and form are separable and independent in *S. benedicti*. By mapping the G_3_ phenotypes to the genotypes of their G_2_ parents, we were able for the first time to separately interrogate the genome for parental- and zygotic-effect loci. We identified four unlinked loci that act, zygotically, in the developing larva, to shape the number and length of defensive chaetae and the presence of anal cirri. The major loci affecting larval size are on two other chromosomes and act maternally. In addition, we detect polygenic maternal effects for size and two loci that act via simultaneous maternal and paternal effect. Our data indicate that larval morphology depends only on the zygotic genome, and size only on the separate genomes of the two parents. Larval size and form depend on different loci, acting in different individuals.

Our findings are congruent with our previous findings from linkage mapping in the G_2_ generation (Zakas et al. 2018), showing that the additive effects estimated from direct measurements of G_2_ are manifest in their transmission of trait variation to the next generation. The present study adds the finding that neither these loci, nor any others in the genome, have detectable maternal or paternal effects on larval anatomy traits. Our combined analysis of G_2_ and G_3_ data allowed us to increase the linkage signal at the zygotically-acting loci, and the resulting narrowed linkage peaks will facilitate the discovery of the causal molecular genes when we anchor the map in the physical genome assembly.

Notwithstanding the success of the QTL-mapping strategy, our data point to some remaining questions about the genetic architectures of larval anatomy. For example, the presence of anal cirri is influenced only by one major-effect zygotic locus, with no evidence for additional polygenic effects. Nevertheless, broods that are entirely homozygous for the alternative alleles at the major-effect locus include individuals with and without cirri (albeit at very different frequencies). Wild *S. benedicti* planktotrophs always have cirri and wild lecithotrophs always lack them (Gibson et al. 2010), as do their lab-reared descendants, implying that we are missing a component of the genetic basis for this trait.

This result may in part be down to numbers. Although our study includes thousands of individuals, they come from only 115 full-sib families. Particular homozygous genotype combinations are rare in this collection. For example, only 3 of the families involve crosses of a male and a female that are both homozygous LL at the chromosome 5 locus affecting cirri. For traits affected by multiple loci, particular parental genotype combinations may be altogether absent. This reinforces an important feature of the genetic architecture of larval life-history in *S. benedicti*: its multilocus basis guarantees that parental types will be exceedingly rare among segregating cross progeny, and most advanced-generation hybrids will be phenotypically intermediate. The genetic architecture therefore implies that the absence of intermediates in nature reflects strong selection against advanced-generation hybrids.

In addition, our findings are partly inconsistent with previous work that studied inheritance of larval traits using classical line-cross methods, comparing phenotype distributions in reciprocal crosses and backcrosses rather than tracking individual genotypes (Zakas and Rockman 2014). That work found, as we did here, that length of larval chaetae is purely zygotic, but it also found that the number of larval chaetae is influenced by both zygotic and maternal effect; our analysis here found no evidence for any maternal effect. That earlier study found that the distribution of chaetae number differed substantially between reciprocal F1s, whose autosomal genomes should be equivalent: F1s with lecithotroph mothers often lacked chaetae entirely, while those with planktotroph mothers almost always had chaetae. The findings from reciprocal F1s suggest that we may be missing maternal effect loci.

We recognize at least two non-exclusive explanations for the missing maternal effects. First, there are two kinds of loci that are not fully interrogated by our experimental cross. The mitochondrial genome in all G_2_s and G_3_s in the current analysis derives from the planktotrophic P0 female founder of the pedigree. Effects of the lecithotroph mitochondrial genome, including interactions between that genome and the rest of the genome, are missing from our analysis but would be present in F1s with lecithotroph mothers. Similarly, our analysis will miss maternal effects that are due to recessive lecithotroph alleles on the X chromosome. Because of the direction of the pedigree-founding cross, the G_2_ females have PP or LP genotypes but not LL genotypes. Although our design gives us access to recessive LL zygotic effects in the G_3_ generation, we would not see recessive LL maternal effects, which would have an effect on F1s with lecithotroph mothers.

The second class of explanation for missing maternal effects invokes epistasis, specifically genes that only show their effects when a large fraction of the maternal genome comes from a single background. In reciprocal F1s, the maternal-effect genotype is 100% lecithotroph or 0%. In the present study, the fraction of lecithotrophic maternal genome among the G_2_s ranges from 19% to 74%. We failed to find a linear dependence of any larval anatomy trait on percent-lecithotrophic-alleles in the mother (controlling for the direct effects of the zygotic-effect alleles that the mother transmits). To test the hypothesis of a polygenic epistatic effect that manifests only at the extremes, we tested the G_3_ data for quadratic and cubic effects of maternal genetic background and found no significant effects. Thus, if present, maternal-effect epistasis is likely to involve higher order interactions that are absent or rare in the G_3_ population. In nature, however, most hybrids will be F1s, with purely lecithotroph or planktotroph maternal backgrounds, and these animals will most often backcross to the locally common form rather than intercross with other hybrids. We therefore anticipate that higher-order epistasis, manifest only in “pure” genetic backgrounds, could play an important role in inter-population gene flow in wild *S. benedicti*.

One surprising result of our analysis is the discovery of paternal-effect loci acting on larval size. This finding is consistent with our earlier line-cross study of larval size (Zakas and Rockman 2014), which found statistical evidence for more-than-maternal genetics, including negative effects of lecithotrophic paternity on larval size, at least in some maternal backgrounds. Paternal genetic effects in species without parental care are rare but not unprecedented (Crean and Bonduriansky 2014; Tigreros et al. 2019). The biology of *Streblospio benedicti* permits several possible mechanisms of paternal effect. Because our larval size trait is measured after embryogenesis, it may reflect both prezygotic variation, such as egg size, and postzygotic variation, such as differential allocation of yolk resources to growth and metabolism.

Although egg-size is superficially a purely maternal trait, females may alter oogenesis in response to genetically variable signals from males. *S. benedicti* males and females do not have to interact to mate; males release spermatophores onto the benthic mud and females retrieve them and store sperm for subsequent fertilizations. Plausible mechanisms for paternal effects in this case include effects of spermatophore components on oogenesis. During laboratory matings, males and females are paired in small arenas for days or weeks, during which time water-borne chemical signaling between male and female is also possible. A suggestion of signaling between adults comes from an account of experimental crosses between genetically divergent *Streblospio*, populations and species: incubated with distantly related females, *S. benedicti* males stopped producing spermatophores, even though they readily produce spermatophores when housed by themselves (Schulze et al. 2000).

Alternatively, paternal genetic variation may influence larval size through postzygotic effects on growth and metabolism. Although such paternal effects could be mediated directly by sperm-borne cytoplasmic components (Seidel et al. 2011), a parsimonious hypothesis for antagonistic maternal and paternal effects in *Streblospio* is genetic variation in imprinting, or parent-of-origin-dependent gene expression. Imprinting is fundamentally different from ordinary parental effects; the former involves gene expression in the offspring and the latter gene expression in the parents. Nevertheless, genotype-dependent imprinting generates a genotype-by-parent-of-origin interaction whose phenotypic effects can mimic antagonistic maternal and paternal effects. Alleles at the two paternal-effect loci may themselves be differentially imprinted, or they may act in trans to differentially regulate imprinting of other loci. To test imprinting models with transmission genetics, we would require additional data on the genotypes of individual G_3_s and their parental origins (Hager et al. 2008). Imprinting often involves DNA methylation, and so we may also predict differences in sperm DNA methylation as a function of genotype at the paternal-effect loci.

The larvae of marine invertebrates provide a rich testing ground for models of life-history evolution, with major transitions between planktotrophy and lecithotrophy a frequent motif (Thorson 1950; Jablonski and Lutz 1983) and powerful example of offspring number/quality tradeoffs (Vance 1973; Smith and Fretwell 1974). Our results show that the genetic basis for these transitions has the striking characteristic that the genotypes of three different individuals influence the phenotype of each offspring. Larval chaetae and cirri are likely to be beneficial in small planktotrophic larvae and disadvantageous in large lecithotrophic larvae, yet size and larval anatomy are governed by unlinked loci active in different generations within a family. Indirect genetic effects, including maternal and paternal effects, are increasingly recognized as a ubiquitous feature of inheritance (Young et al. 2019). They have the potential to alter the pace and trajectory of a population’s response to selection (Wade 1998; Wolf and Brodie 1998; Wolf 2000; Drown and Wade 2014), and future models of larval life-history evolution will need to take them into account.

---

**Supplementary Figure 1.**
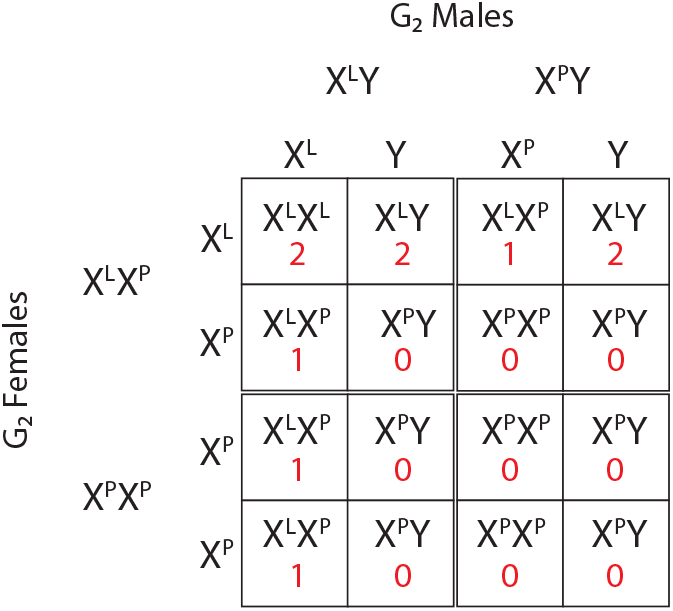
Punnett squares showing the genotypes among the four possible crosses among G_2_s. Assuming hemizygotes are phenotypically equivalent to homozygotes, the red numbers record the effective number of L alleles for each G_3_ genotype. The effective lecithotroph allele count for each cross (*Q*_T_ in model 3) is then the proportion of L alleles for each cross: 5/8, 3/8, 2/8, and 0.

## ACKNOWLEDGMENTS

We thank D Tandon, I Ukegbu, R Freih, C Fayyazi, and L Jessell for laboratory assistance with maintenance and crossing of the animals, L Noble for help with image analysis, and members of the Rockman and Zakas labs for valuable feedback on drafts. This research is funded by the Zegar Family Foundation, NSF grant IOS-1350926 to MVR and NIH grant GM108396 to CZ.

## Supplementary Material

**Table S1. G_*3*_ linkage scan results**

Each box shows the log likelihoods for three models: a null model, a model that includes the single additional additive zygotic QTL (one beta) that provides the greatest increase in log likelihood, and a model that includes the single additional parental-effect QTL (two betas) that provides the greatest increase in log likelihood. To test whether the addition of the QTL is significant, each of the QTL models is compared to the null by a likelihood ratio test whose test statistic (Chisq) is shown. Genome-wide significance thresholds (p = 0.05) for the test statistics were determined by permutation. When the QTL is significant (i.e., “QTL test Chisq” exceeds “Threshold Chisq (p=0.05)”), the two-beta and one-beta models are compared and a likelihood ratio test statistic calculated, as reported in the “Chisq 1 vs 2 betas” column. If this value exceeds, 3.84 (the p=0.05 threshold for significance for a chi-square-distributed random variable with one degree of freedom), we reject the one-beta model and proceed to test uniparental models, comparing each to the two-beta model again by likelihood ratio test with one degree of freedom. At each successive round of genome scan, significant QTL from the previous round were incorporated into the relevant null model. “Script” names the R script in supplementary file 1 that reproduces the results shown.

**Table S2. Heritability tests, including heritability estimates where significant.**

**Table S3. Joint G*_2_*-G*_3_* additive zygotic linkage scan results**

Each box shows the log likelihoods for two models, a null model and a second model that includes the single additional QTL that provides the greatest increase in log likelihood. These models are compared by a likelihood ratio test whose test statistic is shown. Genome-wide significance thresholds (p = 0.05) for the test statistics were determined by permutation. QTL were incorporated into successive rounds of genome scans if their test statistic exceeded the permutation-based threshold.

**Supplementary File 1: Data and Code**

Supplementary File 1 contains genotype data, phenotype data, and R scripts to replicate the analyses reported in the text. The zipped file contains two directories:

### DataHarmonization

This directory includes the G_2_ genotypes (previously reported in Zakas et al. 2018), the G_3_ phenotypes, and the R script G_2_andG_3_DataHarmonization.R. This script combines the genotype data for G_2_ larvae with the new phenotype data for G_3_ larvae. It trims the genotype data to non-redundant markers, imputes missing data on the X chromosome, and generates the G_3_AnalysisObjects.R file that is used in all downstream analyses.

### DataAnalysis

This directory contains G_3_AnalysisObjects.R, the dataset on which all analyses are performed. It also contains commented scripts that perform all of the analyses reported in the paper. These R script files also report the complete results of all analyses, including permutations. Script DataFigures.R contains code to reproduce the figures from the paper.

