## Supplementary File 1. Data and Code for "Baby makes three: maternal, paternal, and zygotic genetic effects shape larval phenotypic evolution": AreaScanAndTestsForDominanceEtc.R

```
# Area trait scans
# This script includes forward searches for QTL affecting G3 larval size and permutations for each round of search.
# Then there are tests for dominance, polygenic effects, and epistasis among the significant QTL
# Then there are tests for direct (zygotic) effects on G2 larval size

library(qtl)
library(lme4)

load(file = "G3AnalysisObjects.R")

# Create an empty scanone object to hold the results
scan.holder <- scanone(MatGeno, pheno = "nChaetae")
# This will give a "calc.genoprob" warning; this can be ignored because we are just making an empty scanone object to fill in later.
scan.holder$lod <- 0
scan.holder[,4:7] <- 0

AreaMZscan1 <- function(geno.mat, geno.pat, scan.object) {
	area.pheno <- geno.mat$pheno$area
	dadID <- geno.mat$pheno$Dad_G2_Name
	momID <-  geno.mat$pheno$Mom_G2_Name
	null.LL <- logLik(lmer(area.pheno  ~ 1 + (1|momID), REML = F))[1]
		for(j in 1: 10){	# This function is only for the autosomes
		chr <- names(nmar(geno.mat))[j]
		area.both <- NULL
		area.summed <- NULL
		area.mat <- NULL
		area.pat <- NULL

			for(i in 1:(nmar(geno.mat)[chr])){ 							
				qtlMom <- geno.mat$geno[[chr]]$data[,i] -1 # Count of lecithotroph alleles: 0, 1, or 2.
				qtlDad <- geno.pat$geno[[chr]]$data[,i] -1 # Count of lecithotroph alleles: 0, 1, or 2.
				qtlAll <- qtlMom + qtlDad				
		# Fit mixed models		
				area.both[i] <- logLik(lmer(area.pheno  ~ qtlMom + qtlDad + (1|momID), REML = F))[1]
				area.summed[i] <- logLik(lmer(area.pheno ~ qtlAll + (1|momID), REML = F))[1]
				area.mat[i] <- logLik(lmer(area.pheno  ~ qtlMom + (1|momID), REML = F))[1]
				area.pat[i] <- logLik(lmer(area.pheno  ~ qtlDad + (1|momID), REML = F))[1]			
		}				
	scan.object[which(scan.object$chr == chr),3] <- area.both
	scan.object[which(scan.object$chr == chr),4] <- area.summed
	scan.object[which(scan.object$chr == chr),5] <- area.mat
	scan.object[which(scan.object$chr == chr),6] <- area.pat
	scan.object[which(scan.object$chr == chr),7] <- null.LL
	}

			for(j in 11){	# X only
		chr <- names(nmar(geno.mat))[j]
		area.both <- NULL
		area.summed <- NULL
		area.mat <- NULL
		area.pat <- NULL

			for(i in 1:(nmar(geno.mat)[chr])){ 	
						
#				print(paste(chr, i))	
						
				qtlMom <- geno.mat$geno[[chr]]$data[,i] -1 # Count of lecithotroph alleles: 0, or 1.
				qtlDad <- geno.pat$geno[[chr]]$data[,i] -1 # Count of lecithotroph alleles: 0, or 1.
				qtlAll <- (1/4)*qtlDad + (3/8)*qtlMom # X-specific G3 additive effect parameterization
		# Fit mixed models		
				area.both[i] <- logLik(lmer(area.pheno  ~ qtlMom + qtlDad + (1|momID), REML = F))[1]
				area.summed[i] <- logLik(lmer(area.pheno ~ qtlAll + (1|momID), REML = F))[1]
				area.mat[i] <- logLik(lmer(area.pheno  ~ qtlMom + (1|momID), REML = F))[1]
				area.pat[i] <- logLik(lmer(area.pheno  ~ qtlDad + (1|momID), REML = F))[1]
		}				
	scan.object[which(scan.object$chr == chr),3] <- area.both
	scan.object[which(scan.object$chr == chr),4] <- area.summed
	scan.object[which(scan.object$chr == chr),5] <- area.mat
	scan.object[which(scan.object$chr == chr),6] <- area.pat
	scan.object[which(scan.object$chr == chr),7] <- null.LL

	}
	names(scan.object)[3:7] <- c("AreaBoth","AreaSummed","AreaMat","AreaPat", "AreaNull")
		
	return(scan.object)
}


date()
scan.out <- AreaMZscan1(MatGeno, PatGeno, scan.holder)
date()

scan.out[,8:12] <- NA

scan.out[,8:12] <- cbind(2*(scan.out$AreaBoth - scan.out$AreaNull), 2*(scan.out$AreaSummed - scan.out$AreaNull), 2*(scan.out$AreaBoth - scan.out$AreaSummed), 2*(scan.out$AreaBoth - scan.out$AreaMat), 2*(scan.out$AreaBoth - scan.out$AreaPat))
names(scan.out)[8:12] <- c("BothChisq", "SummedChisq", "BetasChisq","MatVBothChisq", "PatVBothChisq")

AreaScan1.results <- scan.out
rm(scan.out)

#### End of first forward search.

# Summary of results for each column
apply(AreaScan1.results[,c(3:6)], 2, function(x){AreaScan1.results[which(x == max(x)),]})

#$AreaBoth
#      chr       pos  AreaBoth AreaSummed   AreaMat   AreaPat  AreaNull BothChisq SummedChisq BetasChisq MatVBothChisq PatVBothChisq
#SNP68   7 0.6208756 -39311.74  -39318.85 -39312.29 -39331.27 -39331.56  39.64944    25.42273    14.2267       1.10906      39.07556
#$AreaSummed
#       chr pos  AreaBoth AreaSummed   AreaMat  AreaPat  AreaNull BothChisq SummedChisq BetasChisq MatVBothChisq PatVBothChisq
#SNP443   7   0 -39311.74   -39318.7 -39312.29 -39331.3 -39331.56  39.63228    25.72968   13.90259        1.0919      39.10891
#$AreaMat
#       chr       pos  AreaBoth AreaSummed   AreaMat   AreaPat  AreaNull BothChisq SummedChisq BetasChisq MatVBothChisq PatVBothChisq
#SNP443   7 0.0000000 -39311.74  -39318.70 -39312.29 -39331.30 -39331.56  39.63228    25.72968   13.90259       1.09190      39.10891
#SNP68    7 0.6208756 -39311.74  -39318.85 -39312.29 -39331.27 -39331.56  39.64944    25.42273   14.22670       1.10906      39.07556
#$AreaPat
#        chr pos  AreaBoth AreaSummed   AreaMat   AreaPat  AreaNull BothChisq SummedChisq BetasChisq MatVBothChisq PatVBothChisq
#SNP1249   X   0 -39326.06  -39331.27 -39330.81 -39326.46 -39331.56  10.99843   0.5853389   10.41309      9.504917     0.8039249


# permutations for autosomes and X, areascan1

perm.combo <- c(MatGeno,PatGeno)
combo.dups <- findDupMarkers(perm.combo, adjacent.only = F)
MGnd <- drop.markers(MatGeno, unlist(combo.dups))
PGnd <- drop.markers(PatGeno, unlist(combo.dups))

# How many columns of results?
n.results <- 5
# How many permutations?
n.perms <- 200

#Prepare data for permutation by parent
cross.rows <- row.names(unique(data.frame(MatGeno$pheno$Male_ID, MatGeno$pheno$Female_ID)))
#Make mini cross objects that include each cross only once. 
#mini.G3s.nodups <- subset(G3s.nodups, ind = as.numeric(cross.rows))
mini.MGnd <- subset(MGnd, ind = as.numeric(cross.rows))
#make a mini.Gpat cross object:
mini.PGnd <- subset(MGnd, ind = as.numeric(cross.rows))

perm.scan.holder <- scanone(mini.MGnd, pheno = "nChaetae")
perm.scan.holder$lod <- 0
perm.scan.holder[,3:(2+n.results)] <- 0
perm.results <- matrix(nrow = n.perms, ncol = 2*n.results) # matrix(nrow = n.perms, ncol = n.results)

date()
for(m in 1:n.perms){
	
	male.perm <- data.frame(levels(MatGeno$pheno$Male_ID), levels(MatGeno$pheno$Male_ID))
	# Permutations are stratified by the F1 mother of the F2 fathers, i.e., 1A, 3C, 6F, 8H.
	male.perm[1:10,2] <- sample(male.perm[1:10,1])	
	male.perm[11:24,2] <- sample(male.perm[11:24,1])	
	male.perm[25:32,2] <- sample(male.perm[25:32,1])	
	male.perm[33:53,2] <- sample(male.perm[33:53,1])	

	female.perm <- data.frame(levels(MatGeno$pheno$Female_ID), levels(MatGeno$pheno$Female_ID)) 
	# Permutations are stratified by the F1 mother of the F2 mothers, i.e., 1A, 3C, 6F, 8H.
	female.perm[1:23,2] <- sample(female.perm[1:23,1])	
	female.perm[24:60,2] <- sample(female.perm[24:60,1])		
	female.perm[61:81,2] <- sample(female.perm[61:81,1])	
	female.perm[82:115,2] <- sample(female.perm[82:115,1])	
	
#	perm.G3s <- G3s.nodups
	perm.Gmat <- MGnd
	perm.Gpat <- PGnd
	
	for(i in 1:dim(MatGeno$pheno)[1]){
		permdad <- male.perm[which(male.perm[,1] == MatGeno$pheno$Male_ID[i]),2]
		permmom <- female.perm[which(female.perm[,1] == MatGeno$pheno$Female_ID[i]),2]
		for(j in 1:11){
			perm.Gpat$geno[[j]]$data[i,] <- mini.PGnd$geno[[j]]$data[which(mini.MGnd$pheno$Male_ID == permdad)[1],]
			perm.Gmat$geno[[j]]$data[i,] <- mini.MGnd$geno[[j]]$data[which(mini.MGnd$pheno$Female_ID == permmom)[1],]
		}
	}
	
permscan <- AreaMZscan1(perm.Gmat, perm.Gpat, perm.scan.holder)

perm.results[m,1:n.results] <- apply(as.matrix(permscan[permscan$chr != "X",3:(2+n.results)]), 2, max)
perm.results[m, (n.results+1):(2*n.results)] <- apply(as.matrix(permscan[permscan$chr == "X",3:(2+n.results)]), 2, max)
perm.out <- data.frame(perm.results)
names(perm.out)  <- colnames(permscan)[3:(2+n.results)]
print(m); print(date());
}

rm(perm.results)
perm.out[,11:20] <- NA

perm.out[,11:20] <- cbind(2*(perm.out$AreaBoth - perm.out$AreaNull), 2*(perm.out$AreaSummed - perm.out$AreaNull), 2*(perm.out$AreaBoth - perm.out$AreaSummed), 2*(perm.out$AreaBoth - perm.out$AreaMat), 2*(perm.out$AreaBoth - perm.out$AreaPat), 2*(perm.out[,6] - perm.out$AreaNull), 2*(perm.out[,7] - perm.out$AreaNull), 2*(perm.out[,6] - perm.out[,7]), 2*(perm.out[,6]- perm.out[,8]), 2*(perm.out[,6] - perm.out[,9]))
names(perm.out)[11:20] <- c("BothChisq", "SummedChisq", "BetasChisq","MatVBothChisq", "PatVBothChisq", "XBothChisq", "XSummedChisq", "XBetasChisq","XMatVBothChisq", "XPatVBothChisq")

apply(perm.out[,11:20], 2, function(x){quantile(x,0.95)})
#     BothChisq    SummedChisq     BetasChisq  MatVBothChisq  PatVBothChisq     XBothChisq   XSummedChisq    XBetasChisq 
#     12.420760       9.746515       6.153530       4.899335       7.535075       8.884158       5.538124       5.258517 
#XMatVBothChisq XPatVBothChisq 
#      4.428420       6.814911 

perm.out -> AreaMZScan1Perms
save(AreaMZScan1Perms, file = "AreaScan1.200perms.R")
rm(perm.out)

# At peak marker 7:SNP68 the 2-beta model is prefered to the 1-beta (chisq = 14.227). The paternal-only model is strongly rejected and the maternal-only model is not rejected.
# SNP68 is accepted as a maternal-effect QTL.

#################

area.qtl.7.68 <- MatGeno$geno[[7]]$data[,"SNP68"] - 1
last.column <- dim(MatGeno$pheno)[2]
MatGeno$pheno[,last.column+1] <- area.qtl.7.68
names(MatGeno$pheno)[last.column + 1] <- "area.qtl.7.68"

AreaMZscan2 <- function(geno.mat, geno.pat, scan.object) {
	area.pheno <- geno.mat$pheno$area
	dadID <- geno.mat$pheno$Dad_G2_Name
	momID <-  geno.mat$pheno$Mom_G2_Name
	null.LL <- logLik(lmer(area.pheno  ~ 1 + geno.mat$pheno$area.qtl.7.68 + (1|momID), REML = F))[1]
		for(j in 1: 10){	# This function is only for the autosomes
		chr <- names(nmar(geno.mat))[j]
		area.both <- NULL
		area.summed <- NULL
		area.mat <- NULL
		area.pat <- NULL

			for(i in 1:(nmar(geno.mat)[chr])){ 							
				qtlMom <- geno.mat$geno[[chr]]$data[,i] -1 # Count of lecithotroph alleles: 0, 1, or 2.
				qtlDad <- geno.pat$geno[[chr]]$data[,i] -1 # Count of lecithotroph alleles: 0, 1, or 2.
				qtlAll <- qtlMom + qtlDad				
		# Fit mixed models		
				area.both[i] <- logLik(lmer(area.pheno  ~ qtlMom + qtlDad +  geno.mat$pheno$area.qtl.7.68 + (1|momID), REML = F))[1]
				area.summed[i] <- logLik(lmer(area.pheno ~ qtlAll +  geno.mat$pheno$area.qtl.7.68 + (1|momID), REML = F))[1]
				area.mat[i] <- logLik(lmer(area.pheno  ~ qtlMom +  geno.mat$pheno$area.qtl.7.68 + (1|momID), REML = F))[1]
				area.pat[i] <- logLik(lmer(area.pheno  ~ qtlDad +  geno.mat$pheno$area.qtl.7.68 + (1|momID), REML = F))[1]			
		}				
	scan.object[which(scan.object$chr == chr),3] <- area.both
	scan.object[which(scan.object$chr == chr),4] <- area.summed
	scan.object[which(scan.object$chr == chr),5] <- area.mat
	scan.object[which(scan.object$chr == chr),6] <- area.pat
	scan.object[which(scan.object$chr == chr),7] <- null.LL
	}

			for(j in 11){	# X only
		chr <- names(nmar(geno.mat))[j]
		area.both <- NULL
		area.summed <- NULL
		area.mat <- NULL
		area.pat <- NULL

			for(i in 1:(nmar(geno.mat)[chr])){ 	
						
#				print(paste(chr, i))	
						
				qtlMom <- geno.mat$geno[[chr]]$data[,i] -1 # Count of lecithotroph alleles: 0, or 1.
				qtlDad <- geno.pat$geno[[chr]]$data[,i] -1 # Count of lecithotroph alleles: 0, or 1.
				qtlAll <- (1/4)*qtlDad + (3/8)*qtlMom # X-specific G3 additive effect parameterization
		# Fit mixed models		
				area.both[i] <- logLik(lmer(area.pheno  ~ qtlMom + qtlDad + geno.mat$pheno$area.qtl.7.68 + (1|momID), REML = F))[1]
				area.summed[i] <- logLik(lmer(area.pheno ~ qtlAll + geno.mat$pheno$area.qtl.7.68 + (1|momID), REML = F))[1]
				area.mat[i] <- logLik(lmer(area.pheno  ~ qtlMom + geno.mat$pheno$area.qtl.7.68 + (1|momID), REML = F))[1]
				area.pat[i] <- logLik(lmer(area.pheno  ~ qtlDad + geno.mat$pheno$area.qtl.7.68 + (1|momID), REML = F))[1]
		}				
	scan.object[which(scan.object$chr == chr),3] <- area.both
	scan.object[which(scan.object$chr == chr),4] <- area.summed
	scan.object[which(scan.object$chr == chr),5] <- area.mat
	scan.object[which(scan.object$chr == chr),6] <- area.pat
	scan.object[which(scan.object$chr == chr),7] <- null.LL

	}
	names(scan.object)[3:7] <- c("AreaBoth","AreaSummed","AreaMat","AreaPat", "AreaNull")
		
	return(scan.object)
}

date()
scan.out <- AreaMZscan2(MatGeno, PatGeno, scan.holder)
date()

scan.out[,8:12] <- NA

scan.out[,8:12] <- cbind(2*(scan.out$AreaBoth - scan.out$AreaNull), 2*(scan.out$AreaSummed - scan.out$AreaNull), 2*(scan.out$AreaBoth - scan.out$AreaSummed), 2*(scan.out$AreaBoth - scan.out$AreaMat), 2*(scan.out$AreaBoth - scan.out$AreaPat))
names(scan.out)[8:12] <- c("BothChisq", "SummedChisq", "BetasChisq","MatVBothChisq", "PatVBothChisq")

AreaScan2.results <- scan.out
rm(scan.out)

#### End of second forward search.

# Summary of results for each column
apply(AreaScan2.results[,c(3:6)], 2, function(x){AreaScan2.results[which(x == max(x)),]})
#$AreaBoth
#        chr      pos  AreaBoth AreaSummed   AreaMat   AreaPat  AreaNull BothChisq SummedChisq BetasChisq MatVBothChisq PatVBothChisq
#SNP49     6 2.087372 -39301.11  -39308.66 -39301.63 -39312.02 -39312.29  22.37141    7.256478   15.11493      1.056148      21.83554
#SNP93     6 2.295514 -39301.11  -39308.66 -39301.63 -39312.02 -39312.29  22.37141    7.256478   15.11493      1.056148      21.83554
#SNP1310   6 2.496541 -39301.11  -39308.66 -39301.63 -39312.02 -39312.29  22.37141    7.256478   15.11493      1.056148      21.83554
#$AreaSummed
#        chr      pos  AreaBoth AreaSummed   AreaMat   AreaPat  AreaNull BothChisq SummedChisq BetasChisq MatVBothChisq PatVBothChisq
#SNP776    6 18.08709 -39303.17  -39306.62 -39303.25 -39312.17 -39312.29  18.23756    11.33884   6.898722     0.1559622      17.99197
#SNP1000   6 18.29302 -39303.17  -39306.62 -39303.25 -39312.17 -39312.29  18.23756    11.33884   6.898722     0.1559622      17.99197
#$AreaMat
#        chr pos  AreaBoth AreaSummed   AreaMat   AreaPat  AreaNull BothChisq SummedChisq BetasChisq MatVBothChisq PatVBothChisq
#SNP1170   6   0 -39301.33  -39307.79 -39301.54 -39312.13 -39312.29  21.91817    8.996824   12.92134     0.4150447      21.60546
#$AreaPat
#        chr pos  AreaBoth AreaSummed   AreaMat   AreaPat  AreaNull BothChisq SummedChisq BetasChisq MatVBothChisq PatVBothChisq
#SNP1249   X   0 -39303.66  -39312.25 -39310.01 -39305.31 -39312.29  17.26379  0.07845681   17.18533      12.70076      3.311691

# permutations for autosomes and X, areascan2

perm.combo <- c(MatGeno,PatGeno)
combo.dups <- findDupMarkers(perm.combo, adjacent.only = F)
MGnd <- drop.markers(MatGeno, unlist(combo.dups))
PGnd <- drop.markers(PatGeno, unlist(combo.dups))

# How many columns of results?
n.results <- 5
# How many permutations?
n.perms <- 200

#Prepare data for permutation by parent
cross.rows <- row.names(unique(data.frame(MatGeno$pheno$Male_ID, MatGeno$pheno$Female_ID)))
#Make mini cross objects that include each cross only once. 
#mini.G3s.nodups <- subset(G3s.nodups, ind = as.numeric(cross.rows))
mini.MGnd <- subset(MGnd, ind = as.numeric(cross.rows))
#make a mini.Gpat cross object:
mini.PGnd <- subset(MGnd, ind = as.numeric(cross.rows))

perm.scan.holder <- scanone(mini.MGnd, pheno = "nChaetae")
perm.scan.holder$lod <- 0
perm.scan.holder[,3:(2+n.results)] <- 0
perm.results <- matrix(nrow = n.perms, ncol = 2*n.results) # matrix(nrow = n.perms, ncol = n.results)

date()
for(m in 1:n.perms){
	
	male.perm <- data.frame(levels(MatGeno$pheno$Male_ID), levels(MatGeno$pheno$Male_ID))
	# Permutations are stratified by the F1 mother of the F2 fathers, i.e., 1A, 3C, 6F, 8H.
	male.perm[1:10,2] <- sample(male.perm[1:10,1])	
	male.perm[11:24,2] <- sample(male.perm[11:24,1])	
	male.perm[25:32,2] <- sample(male.perm[25:32,1])	
	male.perm[33:53,2] <- sample(male.perm[33:53,1])	

	female.perm <- data.frame(levels(MatGeno$pheno$Female_ID), levels(MatGeno$pheno$Female_ID)) 
	# Permutations are stratified by the F1 mother of the F2 mothers, i.e., 1A, 3C, 6F, 8H.
	female.perm[1:23,2] <- sample(female.perm[1:23,1])	
	female.perm[24:60,2] <- sample(female.perm[24:60,1])		
	female.perm[61:81,2] <- sample(female.perm[61:81,1])	
	female.perm[82:115,2] <- sample(female.perm[82:115,1])	
	
#	perm.G3s <- G3s.nodups
	perm.Gmat <- MGnd
	perm.Gpat <- PGnd
	
	for(i in 1:dim(MatGeno$pheno)[1]){
		permdad <- male.perm[which(male.perm[,1] == MatGeno$pheno$Male_ID[i]),2]
		permmom <- female.perm[which(female.perm[,1] == MatGeno$pheno$Female_ID[i]),2]
		for(j in 1:11){
			perm.Gpat$geno[[j]]$data[i,] <- mini.PGnd$geno[[j]]$data[which(mini.MGnd$pheno$Male_ID == permdad)[1],]
			perm.Gmat$geno[[j]]$data[i,] <- mini.MGnd$geno[[j]]$data[which(mini.MGnd$pheno$Female_ID == permmom)[1],]
		}
	}
	
permscan <- AreaMZscan2(perm.Gmat, perm.Gpat, perm.scan.holder)

perm.results[m,1:n.results] <- apply(as.matrix(permscan[permscan$chr != "X",3:(2+n.results)]), 2, max)
perm.results[m, (n.results+1):(2*n.results)] <- apply(as.matrix(permscan[permscan$chr == "X",3:(2+n.results)]), 2, max)
perm.out <- data.frame(perm.results)
names(perm.out)  <- colnames(permscan)[3:(2+n.results)]
print(m); print(date());
}


rm(perm.results)
perm.out[,11:20] <- NA

perm.out[,11:20] <- cbind(2*(perm.out$AreaBoth - perm.out$AreaNull), 2*(perm.out$AreaSummed - perm.out$AreaNull), 2*(perm.out$AreaBoth - perm.out$AreaSummed), 2*(perm.out$AreaBoth - perm.out$AreaMat), 2*(perm.out$AreaBoth - perm.out$AreaPat), 2*(perm.out[,6] - perm.out$AreaNull), 2*(perm.out[,7] - perm.out$AreaNull), 2*(perm.out[,6] - perm.out[,7]), 2*(perm.out[,6]- perm.out[,8]), 2*(perm.out[,6] - perm.out[,9]))
names(perm.out)[11:20] <- c("BothChisq", "SummedChisq", "BetasChisq","MatVBothChisq", "PatVBothChisq", "XBothChisq", "XSummedChisq", "XBetasChisq","XMatVBothChisq", "XPatVBothChisq")

apply(perm.out[,11:20], 2, function(x){quantile(x,0.95)})
#     BothChisq    SummedChisq     BetasChisq  MatVBothChisq  PatVBothChisq     XBothChisq   XSummedChisq    XBetasChisq 
#     12.807721      10.030570       6.422031       5.670141       7.763151      11.340917       6.499351       5.928858 
#XMatVBothChisq XPatVBothChisq 
#      5.991032       6.552809 

perm.out -> AreaMZScan2Perms
save(AreaMZScan2Perms, file = "AreaScan2.200perms.R")
rm(perm.out)

# At peak marker 6:SNP93 (or 49 or 1310) the 2-beta model is prefered to the 1-beta (chisq = 15.11). The paternal-only model is strongly rejected and the maternal-only model is not rejected.
# SNP93 is accepted as a maternal-effect QTL.

##############################

area.qtl.6.93 <- MatGeno$geno[[6]]$data[,"SNP93"] - 1

last.column <- dim(MatGeno$pheno)[2]
MatGeno$pheno[,last.column+1] <- area.qtl.6.93
names(MatGeno$pheno)[last.column + 1] <- "area.qtl.6.93"

AreaMZscan3 <- function(geno.mat, geno.pat, scan.object) {
	area.pheno <- geno.mat$pheno$area
	dadID <- geno.mat$pheno$Dad_G2_Name
	momID <-  geno.mat$pheno$Mom_G2_Name
	null.LL <- logLik(lmer(area.pheno  ~ 1 + geno.mat$pheno$area.qtl.7.68 + geno.mat$pheno$area.qtl.6.93 + (1|momID), REML = F))[1]
		for(j in 1: 10){	# This function is only for the autosomes
		chr <- names(nmar(geno.mat))[j]
		area.both <- NULL
		area.summed <- NULL
		area.mat <- NULL
		area.pat <- NULL

			for(i in 1:(nmar(geno.mat)[chr])){ 							
				qtlMom <- geno.mat$geno[[chr]]$data[,i] -1 # Count of lecithotroph alleles: 0, 1, or 2.
				qtlDad <- geno.pat$geno[[chr]]$data[,i] -1 # Count of lecithotroph alleles: 0, 1, or 2.
				qtlAll <- qtlMom + qtlDad				
		# Fit mixed models		
				area.both[i] <- logLik(lmer(area.pheno  ~ qtlMom + qtlDad +  geno.mat$pheno$area.qtl.7.68 + geno.mat$pheno$area.qtl.6.93 + (1|momID), REML = F))[1]
				area.summed[i] <- logLik(lmer(area.pheno ~ qtlAll +  geno.mat$pheno$area.qtl.7.68 + geno.mat$pheno$area.qtl.6.93 + (1|momID), REML = F))[1]
				area.mat[i] <- logLik(lmer(area.pheno  ~ qtlMom +  geno.mat$pheno$area.qtl.7.68 + geno.mat$pheno$area.qtl.6.93 + (1|momID), REML = F))[1]
				area.pat[i] <- logLik(lmer(area.pheno  ~ qtlDad +  geno.mat$pheno$area.qtl.7.68 + geno.mat$pheno$area.qtl.6.93 + (1|momID), REML = F))[1]			
		}				
	scan.object[which(scan.object$chr == chr),3] <- area.both
	scan.object[which(scan.object$chr == chr),4] <- area.summed
	scan.object[which(scan.object$chr == chr),5] <- area.mat
	scan.object[which(scan.object$chr == chr),6] <- area.pat
	scan.object[which(scan.object$chr == chr),7] <- null.LL
	}

			for(j in 11){	# X only
		chr <- names(nmar(geno.mat))[j]
		area.both <- NULL
		area.summed <- NULL
		area.mat <- NULL
		area.pat <- NULL

			for(i in 1:(nmar(geno.mat)[chr])){ 	
						
#				print(paste(chr, i))	
						
				qtlMom <- geno.mat$geno[[chr]]$data[,i] -1 # Count of lecithotroph alleles: 0, or 1.
				qtlDad <- geno.pat$geno[[chr]]$data[,i] -1 # Count of lecithotroph alleles: 0, or 1.
				qtlAll <- (1/4)*qtlDad + (3/8)*qtlMom # X-specific G3 additive effect parameterization
		# Fit mixed models		
				area.both[i] <- logLik(lmer(area.pheno  ~ qtlMom + qtlDad + geno.mat$pheno$area.qtl.7.68 + geno.mat$pheno$area.qtl.6.93 + (1|momID), REML = F))[1]
				area.summed[i] <- logLik(lmer(area.pheno ~ qtlAll + geno.mat$pheno$area.qtl.7.68 + geno.mat$pheno$area.qtl.6.93 + (1|momID), REML = F))[1]
				area.mat[i] <- logLik(lmer(area.pheno  ~ qtlMom + geno.mat$pheno$area.qtl.7.68 + geno.mat$pheno$area.qtl.6.93 + (1|momID), REML = F))[1]
				area.pat[i] <- logLik(lmer(area.pheno  ~ qtlDad + geno.mat$pheno$area.qtl.7.68 + geno.mat$pheno$area.qtl.6.93 + (1|momID), REML = F))[1]
		}				
	scan.object[which(scan.object$chr == chr),3] <- area.both
	scan.object[which(scan.object$chr == chr),4] <- area.summed
	scan.object[which(scan.object$chr == chr),5] <- area.mat
	scan.object[which(scan.object$chr == chr),6] <- area.pat
	scan.object[which(scan.object$chr == chr),7] <- null.LL

	}
	names(scan.object)[3:7] <- c("AreaBoth","AreaSummed","AreaMat","AreaPat", "AreaNull")
		
	return(scan.object)
}

date()
scan.out <- AreaMZscan3(MatGeno, PatGeno, scan.holder)
date()

scan.out[,8:12] <- NA

scan.out[,8:12] <- cbind(2*(scan.out$AreaBoth - scan.out$AreaNull), 2*(scan.out$AreaSummed - scan.out$AreaNull), 2*(scan.out$AreaBoth - scan.out$AreaSummed), 2*(scan.out$AreaBoth - scan.out$AreaMat), 2*(scan.out$AreaBoth - scan.out$AreaPat))
names(scan.out)[8:12] <- c("BothChisq", "SummedChisq", "BetasChisq","MatVBothChisq", "PatVBothChisq")

AreaScan3.results <- scan.out
rm(scan.out)

#### End of third forward search.

# Summary of results for each column
apply(AreaScan3.results[,c(3:6)], 2, function(x){AreaScan3.results[which(x == max(x)),]})
#$AreaBoth
#        chr pos AreaBoth AreaSummed   AreaMat   AreaPat  AreaNull BothChisq SummedChisq BetasChisq MatVBothChisq PatVBothChisq
#SNP1249   X   0 -39292.8  -39301.63 -39298.98 -39294.79 -39301.63  17.67026  0.01325674     17.657      12.37213      3.980629
#$AreaSummed
#       chr      pos  AreaBoth AreaSummed   AreaMat   AreaPat  AreaNull BothChisq SummedChisq BetasChisq MatVBothChisq PatVBothChisq
#SNP687   X 33.85628 -39296.72  -39297.82 -39296.75 -39301.55 -39301.63  9.822914    7.618038   2.204877    0.04660091      9.660682
#$AreaMat
#        chr      pos  AreaBoth AreaSummed   AreaMat   AreaPat  AreaNull BothChisq SummedChisq BetasChisq MatVBothChisq PatVBothChisq
#SNP1073   X 28.62778 -39296.31  -39299.15 -39296.62 -39301.44 -39301.63   10.6453    4.967191   5.678106     0.6229535      10.25391
#$AreaPat
#        chr pos AreaBoth AreaSummed   AreaMat   AreaPat  AreaNull BothChisq SummedChisq BetasChisq MatVBothChisq PatVBothChisq
#SNP1249   X   0 -39292.8  -39301.63 -39298.98 -39294.79 -39301.63  17.67026  0.01325674     17.657      12.37213      3.980629

# permutations for autosomes and X, areascan3

perm.combo <- c(MatGeno,PatGeno)
combo.dups <- findDupMarkers(perm.combo, adjacent.only = F)
MGnd <- drop.markers(MatGeno, unlist(combo.dups))
PGnd <- drop.markers(PatGeno, unlist(combo.dups))

# How many columns of results?
n.results <- 5
# How many permutations?
n.perms <- 200

#Prepare data for permutation by parent
cross.rows <- row.names(unique(data.frame(MatGeno$pheno$Male_ID, MatGeno$pheno$Female_ID)))
#Make mini cross objects that include each cross only once. 
#mini.G3s.nodups <- subset(G3s.nodups, ind = as.numeric(cross.rows))
mini.MGnd <- subset(MGnd, ind = as.numeric(cross.rows))
#make a mini.Gpat cross object:
mini.PGnd <- subset(MGnd, ind = as.numeric(cross.rows))

perm.scan.holder <- scanone(mini.MGnd, pheno = "nChaetae")
perm.scan.holder$lod <- 0
perm.scan.holder[,3:(2+n.results)] <- 0
perm.results <- matrix(nrow = n.perms, ncol = 2*n.results) # matrix(nrow = n.perms, ncol = n.results)

date()
for(m in 1:n.perms){
	
	male.perm <- data.frame(levels(MatGeno$pheno$Male_ID), levels(MatGeno$pheno$Male_ID))
	# Permutations are stratified by the F1 mother of the F2 fathers, i.e., 1A, 3C, 6F, 8H.
	male.perm[1:10,2] <- sample(male.perm[1:10,1])	
	male.perm[11:24,2] <- sample(male.perm[11:24,1])	
	male.perm[25:32,2] <- sample(male.perm[25:32,1])	
	male.perm[33:53,2] <- sample(male.perm[33:53,1])	

	female.perm <- data.frame(levels(MatGeno$pheno$Female_ID), levels(MatGeno$pheno$Female_ID)) 
	# Permutations are stratified by the F1 mother of the F2 mothers, i.e., 1A, 3C, 6F, 8H.
	female.perm[1:23,2] <- sample(female.perm[1:23,1])	
	female.perm[24:60,2] <- sample(female.perm[24:60,1])		
	female.perm[61:81,2] <- sample(female.perm[61:81,1])	
	female.perm[82:115,2] <- sample(female.perm[82:115,1])	
	
#	perm.G3s <- G3s.nodups
	perm.Gmat <- MGnd
	perm.Gpat <- PGnd
	
	for(i in 1:dim(MatGeno$pheno)[1]){
		permdad <- male.perm[which(male.perm[,1] == MatGeno$pheno$Male_ID[i]),2]
		permmom <- female.perm[which(female.perm[,1] == MatGeno$pheno$Female_ID[i]),2]
		for(j in 1:11){
			perm.Gpat$geno[[j]]$data[i,] <- mini.PGnd$geno[[j]]$data[which(mini.MGnd$pheno$Male_ID == permdad)[1],]
			perm.Gmat$geno[[j]]$data[i,] <- mini.MGnd$geno[[j]]$data[which(mini.MGnd$pheno$Female_ID == permmom)[1],]
		}
	}
	
permscan <- AreaMZscan3(perm.Gmat, perm.Gpat, perm.scan.holder)

perm.results[m,1:n.results] <- apply(as.matrix(permscan[permscan$chr != "X",3:(2+n.results)]), 2, max)
perm.results[m, (n.results+1):(2*n.results)] <- apply(as.matrix(permscan[permscan$chr == "X",3:(2+n.results)]), 2, max)
perm.out <- data.frame(perm.results)
names(perm.out)  <- colnames(permscan)[3:(2+n.results)]
print(m); print(date());
}

rm(perm.results)
perm.out[,11:20] <- NA

perm.out[,11:20] <- cbind(2*(perm.out$AreaBoth - perm.out$AreaNull), 2*(perm.out$AreaSummed - perm.out$AreaNull), 2*(perm.out$AreaBoth - perm.out$AreaSummed), 2*(perm.out$AreaBoth - perm.out$AreaMat), 2*(perm.out$AreaBoth - perm.out$AreaPat), 2*(perm.out[,6] - perm.out$AreaNull), 2*(perm.out[,7] - perm.out$AreaNull), 2*(perm.out[,6] - perm.out[,7]), 2*(perm.out[,6]- perm.out[,8]), 2*(perm.out[,6] - perm.out[,9]))
names(perm.out)[11:20] <- c("BothChisq", "SummedChisq", "BetasChisq","MatVBothChisq", "PatVBothChisq", "XBothChisq", "XSummedChisq", "XBetasChisq","XMatVBothChisq", "XPatVBothChisq")

apply(perm.out[,11:20], 2, function(x){quantile(x,0.95)})

#     BothChisq    SummedChisq     BetasChisq  MatVBothChisq  PatVBothChisq     XBothChisq   XSummedChisq    XBetasChisq 
#     14.782931      11.357572       7.733979       8.396555       7.535856      10.811711       7.522500       6.627170 
#XMatVBothChisq XPatVBothChisq 
#      7.048794       5.927768 

perm.out -> AreaMZScan3Perms
save(AreaMZScan3Perms, file = "AreaScan3.200perms.R")
rm(perm.out)

# Peak marker X:SNP1249 is a genome-wide significant QTL exceeding autosomal and X-specific thresholds for the 2-beta model.
# At peak marker X:SNP1249 the 2-beta model is prefered to the 1-beta (chisq = 17.66). The maternal-only model is strongly rejected and the paternal-only model is nominally rejected too (pchisq(3.980629, df = 1, lower = F) = 0.04602637). 
# SNP1249 is accepted as a paternal- and maternal-effect QTL.

##############################

area.qtl.X.1249P <- PatGeno$geno[["X"]]$data[,"SNP1249"] - 1
area.qtl.X.1249M <- MatGeno$geno[["X"]]$data[,"SNP1249"] - 1

last.column <- dim(MatGeno$pheno)[2]
MatGeno$pheno[,last.column+1] <- area.qtl.X.1249P
MatGeno$pheno[,last.column+2] <- area.qtl.X.1249M

names(MatGeno$pheno)[last.column + 1] <- "area.qtl.X.1249P"
names(MatGeno$pheno)[last.column + 2] <- "area.qtl.X.1249M"

AreaMZscan4 <- function(geno.mat, geno.pat, scan.object) {
	area.pheno <- geno.mat$pheno$area
	dadID <- geno.mat$pheno$Dad_G2_Name
	momID <-  geno.mat$pheno$Mom_G2_Name
	null.LL <- logLik(lmer(area.pheno  ~ 1 + geno.mat$pheno$area.qtl.7.68 + geno.mat$pheno$area.qtl.6.93 + geno.mat$pheno$area.qtl.X.1249P + geno.mat$pheno$area.qtl.X.1249M + (1|momID), REML = F))[1]
		for(j in 1: 10){	# This function is only for the autosomes
		chr <- names(nmar(geno.mat))[j]
		area.both <- NULL
		area.summed <- NULL
		area.mat <- NULL
		area.pat <- NULL

			for(i in 1:(nmar(geno.mat)[chr])){ 							
				qtlMom <- geno.mat$geno[[chr]]$data[,i] -1 # Count of lecithotroph alleles: 0, 1, or 2.
				qtlDad <- geno.pat$geno[[chr]]$data[,i] -1 # Count of lecithotroph alleles: 0, 1, or 2.
				qtlAll <- qtlMom + qtlDad				
		# Fit mixed models		
				area.both[i] <- logLik(lmer(area.pheno  ~ qtlMom + qtlDad +  geno.mat$pheno$area.qtl.7.68 + geno.mat$pheno$area.qtl.6.93 + geno.mat$pheno$area.qtl.X.1249P + geno.mat$pheno$area.qtl.X.1249M + (1|momID), REML = F))[1]
				area.summed[i] <- logLik(lmer(area.pheno ~ qtlAll +  geno.mat$pheno$area.qtl.7.68 + geno.mat$pheno$area.qtl.6.93 + geno.mat$pheno$area.qtl.X.1249P + geno.mat$pheno$area.qtl.X.1249M +  (1|momID), REML = F))[1]
				area.mat[i] <- logLik(lmer(area.pheno  ~ qtlMom +  geno.mat$pheno$area.qtl.7.68 + geno.mat$pheno$area.qtl.6.93 + geno.mat$pheno$area.qtl.X.1249P + geno.mat$pheno$area.qtl.X.1249M +  (1|momID), REML = F))[1]
				area.pat[i] <- logLik(lmer(area.pheno  ~ qtlDad +  geno.mat$pheno$area.qtl.7.68 + geno.mat$pheno$area.qtl.6.93 + geno.mat$pheno$area.qtl.X.1249P + geno.mat$pheno$area.qtl.X.1249M +  (1|momID), REML = F))[1]			
		}				
	scan.object[which(scan.object$chr == chr),3] <- area.both
	scan.object[which(scan.object$chr == chr),4] <- area.summed
	scan.object[which(scan.object$chr == chr),5] <- area.mat
	scan.object[which(scan.object$chr == chr),6] <- area.pat
	scan.object[which(scan.object$chr == chr),7] <- null.LL
	}

			for(j in 11){	# X only
		chr <- names(nmar(geno.mat))[j]
		area.both <- NULL
		area.summed <- NULL
		area.mat <- NULL
		area.pat <- NULL

			for(i in 1:(nmar(geno.mat)[chr])){ 	
						
#				print(paste(chr, i))	
						
				qtlMom <- geno.mat$geno[[chr]]$data[,i] -1 # Count of lecithotroph alleles: 0, or 1.
				qtlDad <- geno.pat$geno[[chr]]$data[,i] -1 # Count of lecithotroph alleles: 0, or 1.
				qtlAll <- (1/4)*qtlDad + (3/8)*qtlMom # X-specific G3 additive effect parameterization
		# Fit mixed models		
				area.both[i] <- logLik(lmer(area.pheno  ~ qtlMom + qtlDad + geno.mat$pheno$area.qtl.7.68 + geno.mat$pheno$area.qtl.6.93 + geno.mat$pheno$area.qtl.X.1249P + geno.mat$pheno$area.qtl.X.1249M +  (1|momID), REML = F))[1]
				area.summed[i] <- logLik(lmer(area.pheno ~ qtlAll + geno.mat$pheno$area.qtl.7.68 + geno.mat$pheno$area.qtl.6.93 + geno.mat$pheno$area.qtl.X.1249P + geno.mat$pheno$area.qtl.X.1249M +  (1|momID), REML = F))[1]
				area.mat[i] <- logLik(lmer(area.pheno  ~ qtlMom + geno.mat$pheno$area.qtl.7.68 + geno.mat$pheno$area.qtl.6.93 + geno.mat$pheno$area.qtl.X.1249P + geno.mat$pheno$area.qtl.X.1249M +  (1|momID), REML = F))[1]
				area.pat[i] <- logLik(lmer(area.pheno  ~ qtlDad + geno.mat$pheno$area.qtl.7.68 + geno.mat$pheno$area.qtl.6.93 + geno.mat$pheno$area.qtl.X.1249P + geno.mat$pheno$area.qtl.X.1249M +  (1|momID), REML = F))[1]
		}				
	scan.object[which(scan.object$chr == chr),3] <- area.both
	scan.object[which(scan.object$chr == chr),4] <- area.summed
	scan.object[which(scan.object$chr == chr),5] <- area.mat
	scan.object[which(scan.object$chr == chr),6] <- area.pat
	scan.object[which(scan.object$chr == chr),7] <- null.LL

	}
	names(scan.object)[3:7] <- c("AreaBoth","AreaSummed","AreaMat","AreaPat", "AreaNull")
		
	return(scan.object)
}

date()
scan.out <- AreaMZscan4(MatGeno, PatGeno, scan.holder)
date()

scan.out[,8:12] <- NA

scan.out[,8:12] <- cbind(2*(scan.out$AreaBoth - scan.out$AreaNull), 2*(scan.out$AreaSummed - scan.out$AreaNull), 2*(scan.out$AreaBoth - scan.out$AreaSummed), 2*(scan.out$AreaBoth - scan.out$AreaMat), 2*(scan.out$AreaBoth - scan.out$AreaPat))
names(scan.out)[8:12] <- c("BothChisq", "SummedChisq", "BetasChisq","MatVBothChisq", "PatVBothChisq")

AreaScan4.results <- scan.out
rm(scan.out)

#### End of fourth forward search.

# Summary of results for each column
apply(AreaScan4.results[,c(3:6)], 2, function(x){AreaScan4.results[which(x == max(x)),]})
# $AreaBoth
        # chr     pos  AreaBoth AreaSummed   AreaMat   AreaPat AreaNull BothChisq SummedChisq BetasChisq MatVBothChisq PatVBothChisq
# SNP1244   9 5.88542 -39284.75   -39292.7 -39290.12 -39287.48 -39292.8  16.08966   0.1878753   15.90179      10.74214      5.445384
# $AreaSummed
       # chr      pos  AreaBoth AreaSummed   AreaMat   AreaPat AreaNull BothChisq SummedChisq BetasChisq MatVBothChisq PatVBothChisq
# SNP687   X 33.85628 -39289.68  -39289.83 -39290.17 -39292.19 -39292.8  6.229371    5.939144  0.2902278     0.9696615      5.014014
# $AreaMat
      # chr      pos  AreaBoth AreaSummed   AreaMat   AreaPat AreaNull BothChisq SummedChisq BetasChisq MatVBothChisq PatVBothChisq
# SNP90   4 5.685495 -39289.15  -39290.68 -39289.16 -39292.77 -39292.8  7.296756    4.237764   3.058992    0.01481658       7.24235
# $AreaPat
       # chr      pos  AreaBoth AreaSummed   AreaMat   AreaPat AreaNull BothChisq SummedChisq BetasChisq MatVBothChisq PatVBothChisq
# SNP190   9 7.127093 -39285.05  -39292.39 -39291.22 -39286.89 -39292.8  15.48668   0.8099083   14.67677      12.32512       3.66103

# At peak marker 9:SNP1244 the 2-beta model is prefered to the 1-beta (chisq = 15.90). Both the maternal-only and paternal-only models are rejected. 
# SNP1244 is accepted as a paternal- and maternal-effect QTL.


# permutations for autosomes and X, areascan4

perm.combo <- c(MatGeno,PatGeno)
combo.dups <- findDupMarkers(perm.combo, adjacent.only = F)
MGnd <- drop.markers(MatGeno, unlist(combo.dups))
PGnd <- drop.markers(PatGeno, unlist(combo.dups))

# How many columns of results?
n.results <- 5
# How many permutations?
n.perms <- 500

#Prepare data for permutation by parent
cross.rows <- row.names(unique(data.frame(MatGeno$pheno$Male_ID, MatGeno$pheno$Female_ID)))
#Make mini cross objects that include each cross only once. 
#mini.G3s.nodups <- subset(G3s.nodups, ind = as.numeric(cross.rows))
mini.MGnd <- subset(MGnd, ind = as.numeric(cross.rows))
#make a mini.Gpat cross object:
mini.PGnd <- subset(MGnd, ind = as.numeric(cross.rows))

perm.scan.holder <- scanone(mini.MGnd, pheno = "nChaetae")
perm.scan.holder$lod <- 0
perm.scan.holder[,3:(2+n.results)] <- 0
perm.results <- matrix(nrow = n.perms, ncol = 2*n.results) # matrix(nrow = n.perms, ncol = n.results)

date()
for(m in 1:n.perms){
	
	male.perm <- data.frame(levels(MatGeno$pheno$Male_ID), levels(MatGeno$pheno$Male_ID))
	# Permutations are stratified by the F1 mother of the F2 fathers, i.e., 1A, 3C, 6F, 8H.
	male.perm[1:10,2] <- sample(male.perm[1:10,1])	
	male.perm[11:24,2] <- sample(male.perm[11:24,1])	
	male.perm[25:32,2] <- sample(male.perm[25:32,1])	
	male.perm[33:53,2] <- sample(male.perm[33:53,1])	

	female.perm <- data.frame(levels(MatGeno$pheno$Female_ID), levels(MatGeno$pheno$Female_ID)) 
	# Permutations are stratified by the F1 mother of the F2 mothers, i.e., 1A, 3C, 6F, 8H.
	female.perm[1:23,2] <- sample(female.perm[1:23,1])	
	female.perm[24:60,2] <- sample(female.perm[24:60,1])		
	female.perm[61:81,2] <- sample(female.perm[61:81,1])	
	female.perm[82:115,2] <- sample(female.perm[82:115,1])	
	
#	perm.G3s <- G3s.nodups
	perm.Gmat <- MGnd
	perm.Gpat <- PGnd
	
	for(i in 1:dim(MatGeno$pheno)[1]){
		permdad <- male.perm[which(male.perm[,1] == MatGeno$pheno$Male_ID[i]),2]
		permmom <- female.perm[which(female.perm[,1] == MatGeno$pheno$Female_ID[i]),2]
		for(j in 1:11){
			perm.Gpat$geno[[j]]$data[i,] <- mini.PGnd$geno[[j]]$data[which(mini.MGnd$pheno$Male_ID == permdad)[1],]
			perm.Gmat$geno[[j]]$data[i,] <- mini.MGnd$geno[[j]]$data[which(mini.MGnd$pheno$Female_ID == permmom)[1],]
		}
	}
	
permscan <- AreaMZscan4(perm.Gmat, perm.Gpat, perm.scan.holder)

perm.results[m,1:n.results] <- apply(as.matrix(permscan[permscan$chr != "X",3:(2+n.results)]), 2, max)
perm.results[m, (n.results+1):(2*n.results)] <- apply(as.matrix(permscan[permscan$chr == "X",3:(2+n.results)]), 2, max)
perm.out <- data.frame(perm.results)
names(perm.out)  <- colnames(permscan)[3:(2+n.results)]
print(m); print(date());
}
rm(perm.results)
perm.out[,11:20] <- NA

perm.out[,11:20] <- cbind(2*(perm.out$AreaBoth - perm.out$AreaNull), 2*(perm.out$AreaSummed - perm.out$AreaNull), 2*(perm.out$AreaBoth - perm.out$AreaSummed), 2*(perm.out$AreaBoth - perm.out$AreaMat), 2*(perm.out$AreaBoth - perm.out$AreaPat), 2*(perm.out[,6] - perm.out$AreaNull), 2*(perm.out[,7] - perm.out$AreaNull), 2*(perm.out[,6] - perm.out[,7]), 2*(perm.out[,6]- perm.out[,8]), 2*(perm.out[,6] - perm.out[,9]))
names(perm.out)[11:20] <- c("BothChisq", "SummedChisq", "BetasChisq","MatVBothChisq", "PatVBothChisq", "XBothChisq", "XSummedChisq", "XBetasChisq","XMatVBothChisq", "XPatVBothChisq")

apply(perm.out[,11:20], 2, function(x){quantile(x,0.95)})
#     BothChisq    SummedChisq     BetasChisq  MatVBothChisq  PatVBothChisq     XBothChisq   XSummedChisq    XBetasChisq 
#     13.638117       9.872829       7.192198       6.054210       8.164856       9.278591       6.730313       5.449783 
#XMatVBothChisq XPatVBothChisq 
#      5.492105       6.213634 

perm.out -> AreaMZScan4Perms
#save(AreaMZScan4Perms, file = "AreaScan4.500perms.R")
#rm(perm.out)

##############################

area.qtl.9.1244P <- PatGeno$geno[[9]]$data[,"SNP1244"] - 1
area.qtl.9.1244M <- MatGeno$geno[[9]]$data[,"SNP1244"] - 1

last.column <- dim(MatGeno$pheno)[2]
MatGeno$pheno[,last.column+1] <- area.qtl.9.1244P
MatGeno$pheno[,last.column+2] <- area.qtl.9.1244M
names(MatGeno$pheno)[last.column + 1] <- "area.qtl.9.1244P"
names(MatGeno$pheno)[last.column + 2] <- "area.qtl.9.1244M"

AreaMZscan5 <- function(geno.mat, geno.pat, scan.object) {
	area.pheno <- geno.mat$pheno$area
	dadID <- geno.mat$pheno$Dad_G2_Name
	momID <-  geno.mat$pheno$Mom_G2_Name
	null.LL <- logLik(lmer(area.pheno  ~ 1 + geno.mat$pheno$area.qtl.7.68 + geno.mat$pheno$area.qtl.6.93 + geno.mat$pheno$area.qtl.X.1249P + geno.mat$pheno$area.qtl.X.1249M + geno.mat$pheno$area.qtl.9.1244P + geno.mat$pheno$area.qtl.9.1244M + (1|momID), REML = F))[1]
		for(j in 1: 10){	# This function is only for the autosomes
		chr <- names(nmar(geno.mat))[j]
		area.both <- NULL
		area.summed <- NULL
		area.mat <- NULL
		area.pat <- NULL

			for(i in 1:(nmar(geno.mat)[chr])){ 							
				qtlMom <- geno.mat$geno[[chr]]$data[,i] -1 # Count of lecithotroph alleles: 0, 1, or 2.
				qtlDad <- geno.pat$geno[[chr]]$data[,i] -1 # Count of lecithotroph alleles: 0, 1, or 2.
				qtlAll <- qtlMom + qtlDad				
		# Fit mixed models		
				area.both[i] <- logLik(lmer(area.pheno  ~ qtlMom + qtlDad +  geno.mat$pheno$area.qtl.7.68 + geno.mat$pheno$area.qtl.6.93 + geno.mat$pheno$area.qtl.X.1249P + geno.mat$pheno$area.qtl.X.1249M + geno.mat$pheno$area.qtl.9.1244P + geno.mat$pheno$area.qtl.9.1244M + (1|momID), REML = F))[1]
				area.summed[i] <- logLik(lmer(area.pheno ~ qtlAll +  geno.mat$pheno$area.qtl.7.68 + geno.mat$pheno$area.qtl.6.93 + geno.mat$pheno$area.qtl.X.1249P + geno.mat$pheno$area.qtl.X.1249M +  geno.mat$pheno$area.qtl.9.1244P + geno.mat$pheno$area.qtl.9.1244M + (1|momID), REML = F))[1]
				area.mat[i] <- logLik(lmer(area.pheno  ~ qtlMom +  geno.mat$pheno$area.qtl.7.68 + geno.mat$pheno$area.qtl.6.93 + geno.mat$pheno$area.qtl.X.1249P + geno.mat$pheno$area.qtl.X.1249M +  geno.mat$pheno$area.qtl.9.1244P + geno.mat$pheno$area.qtl.9.1244M + (1|momID), REML = F))[1]
				area.pat[i] <- logLik(lmer(area.pheno  ~ qtlDad +  geno.mat$pheno$area.qtl.7.68 + geno.mat$pheno$area.qtl.6.93 + geno.mat$pheno$area.qtl.X.1249P + geno.mat$pheno$area.qtl.X.1249M +  geno.mat$pheno$area.qtl.9.1244P + geno.mat$pheno$area.qtl.9.1244M + (1|momID), REML = F))[1]			
		}				
	scan.object[which(scan.object$chr == chr),3] <- area.both
	scan.object[which(scan.object$chr == chr),4] <- area.summed
	scan.object[which(scan.object$chr == chr),5] <- area.mat
	scan.object[which(scan.object$chr == chr),6] <- area.pat
	scan.object[which(scan.object$chr == chr),7] <- null.LL
	}

			for(j in 11){	# X only
		chr <- names(nmar(geno.mat))[j]
		area.both <- NULL
		area.summed <- NULL
		area.mat <- NULL
		area.pat <- NULL

			for(i in 1:(nmar(geno.mat)[chr])){ 	
						
#				print(paste(chr, i))	
						
				qtlMom <- geno.mat$geno[[chr]]$data[,i] -1 # Count of lecithotroph alleles: 0, or 1.
				qtlDad <- geno.pat$geno[[chr]]$data[,i] -1 # Count of lecithotroph alleles: 0, or 1.
				qtlAll <- (1/4)*qtlDad + (3/8)*qtlMom # X-specific G3 additive effect parameterization
		# Fit mixed models		
				area.both[i] <- logLik(lmer(area.pheno  ~ qtlMom + qtlDad + geno.mat$pheno$area.qtl.7.68 + geno.mat$pheno$area.qtl.6.93 + geno.mat$pheno$area.qtl.X.1249P + geno.mat$pheno$area.qtl.X.1249M +  geno.mat$pheno$area.qtl.9.1244P + geno.mat$pheno$area.qtl.9.1244M + (1|momID), REML = F))[1]
				area.summed[i] <- logLik(lmer(area.pheno ~ qtlAll + geno.mat$pheno$area.qtl.7.68 + geno.mat$pheno$area.qtl.6.93 + geno.mat$pheno$area.qtl.X.1249P + geno.mat$pheno$area.qtl.X.1249M +  geno.mat$pheno$area.qtl.9.1244P + geno.mat$pheno$area.qtl.9.1244M + (1|momID), REML = F))[1]
				area.mat[i] <- logLik(lmer(area.pheno  ~ qtlMom + geno.mat$pheno$area.qtl.7.68 + geno.mat$pheno$area.qtl.6.93 + geno.mat$pheno$area.qtl.X.1249P + geno.mat$pheno$area.qtl.X.1249M +  geno.mat$pheno$area.qtl.9.1244P + geno.mat$pheno$area.qtl.9.1244M + (1|momID), REML = F))[1]
				area.pat[i] <- logLik(lmer(area.pheno  ~ qtlDad + geno.mat$pheno$area.qtl.7.68 + geno.mat$pheno$area.qtl.6.93 + geno.mat$pheno$area.qtl.X.1249P + geno.mat$pheno$area.qtl.X.1249M +  geno.mat$pheno$area.qtl.9.1244P + geno.mat$pheno$area.qtl.9.1244M + (1|momID), REML = F))[1]
		}				
	scan.object[which(scan.object$chr == chr),3] <- area.both
	scan.object[which(scan.object$chr == chr),4] <- area.summed
	scan.object[which(scan.object$chr == chr),5] <- area.mat
	scan.object[which(scan.object$chr == chr),6] <- area.pat
	scan.object[which(scan.object$chr == chr),7] <- null.LL

	}
	names(scan.object)[3:7] <- c("AreaBoth","AreaSummed","AreaMat","AreaPat", "AreaNull")
		
	return(scan.object)
}

date()
scan.out <- AreaMZscan5(MatGeno, PatGeno, scan.holder)
date()

scan.out[,8:12] <- NA

scan.out[,8:12] <- cbind(2*(scan.out$AreaBoth - scan.out$AreaNull), 2*(scan.out$AreaSummed - scan.out$AreaNull), 2*(scan.out$AreaBoth - scan.out$AreaSummed), 2*(scan.out$AreaBoth - scan.out$AreaMat), 2*(scan.out$AreaBoth - scan.out$AreaPat))
names(scan.out)[8:12] <- c("BothChisq", "SummedChisq", "BetasChisq","MatVBothChisq", "PatVBothChisq")

AreaScan5.results <- scan.out
rm(scan.out)

#### End of fifth forward search.

# Summary of results for each column
apply(AreaScan5.results[,c(3:6)], 2, function(x){AreaScan5.results[which(x == max(x)),]})

# $AreaBoth
        # chr      pos  AreaBoth AreaSummed   AreaMat   AreaPat  AreaNull BothChisq SummedChisq BetasChisq MatVBothChisq PatVBothChisq
# SNP1342   8 4.365605 -39278.78  -39284.75 -39280.89 -39282.06 -39284.75  11.95179  0.01425842   11.93753      4.231432       6.56231
# $AreaSummed
      # chr      pos  AreaBoth AreaSummed   AreaMat   AreaPat  AreaNull BothChisq SummedChisq BetasChisq MatVBothChisq PatVBothChisq
# SNP90   4 5.685495 -39280.87  -39281.61 -39281.17 -39284.35 -39284.75   7.77401    6.276739    1.49727     0.6109428      6.974069
# $AreaMat
        # chr      pos  AreaBoth AreaSummed   AreaMat   AreaPat  AreaNull BothChisq SummedChisq BetasChisq MatVBothChisq PatVBothChisq
# SNP1342   8 4.365605 -39278.78  -39284.75 -39280.89 -39282.06 -39284.75  11.95179 0.014258416   11.93753      4.231432      6.562310
# SNP856    8 5.210916 -39278.92  -39284.75 -39280.89 -39282.17 -39284.75  11.66639 0.004843627   11.66155      3.946034      6.503097
# SNP1364   8 5.418872 -39278.92  -39284.75 -39280.89 -39282.17 -39284.75  11.66639 0.004843627   11.66155      3.946034      6.503097
# $AreaPat
        # chr       pos  AreaBoth AreaSummed   AreaMat   AreaPat  AreaNull BothChisq  SummedChisq BetasChisq MatVBothChisq PatVBothChisq
# SNP114    8 0.4133690 -39279.53  -39284.75 -39281.83 -39282.01 -39284.75  10.44762 5.299200e-03   10.44232      4.594966      4.958840
# SNP160    8 0.8311002 -39279.20  -39284.75 -39281.40 -39282.01 -39284.75  11.11512 1.047974e-03   11.11408      4.406203      5.626347
# SNP220    8 1.0392637 -39279.20  -39284.75 -39281.40 -39282.01 -39284.75  11.11512 1.047974e-03   11.11408      4.406203      5.626347
# SNP1076   8 1.2474274 -39279.20  -39284.75 -39281.40 -39282.01 -39284.75  11.11512 1.047974e-03   11.11408      4.406203      5.626347
# SNP432    8 2.2882286 -39278.92  -39284.75 -39281.24 -39282.01 -39284.75  11.65722 3.769894e-03   11.65345      4.630635      6.168440
# SNP725    8 2.4963860 -39278.96  -39284.75 -39281.22 -39282.01 -39284.75  11.57950 2.519553e-03   11.57698      4.520061      6.090724
# SNP661    8 2.7045392 -39279.11  -39284.75 -39281.43 -39282.01 -39284.75  11.28535 3.086657e-04   11.28504      4.640484      5.796576
# SNP372    8 2.9127218 -39279.02  -39284.75 -39281.42 -39282.01 -39284.75  11.47024 1.211085e-03   11.46903      4.801907      5.981462
# SNP287    8 3.1208707 -39279.21  -39284.75 -39281.58 -39282.01 -39284.75  11.07913 3.767243e-06   11.07913      4.733194      5.590355
# SNP799    8 3.3290241 -39279.30  -39284.75 -39281.59 -39282.01 -39284.75  10.89756 2.466613e-04   10.89731      4.579107      5.408783
# SNP431    8 3.5371840 -39279.17  -39284.75 -39281.40 -39282.01 -39284.75  11.17334 2.760650e-04   11.17306      4.460232      5.684560
# SNP1038   8 3.7453436 -39279.35  -39284.75 -39281.66 -39282.01 -39284.75  10.80643 4.382936e-04   10.80599      4.618541      5.317653
# SNP1322   8 3.9534987 -39279.17  -39284.75 -39281.40 -39282.01 -39284.75  11.17334 2.760650e-04   11.17306      4.460232      5.684560

# permutations for autosomes and X, areascan5

perm.combo <- c(MatGeno,PatGeno)
combo.dups <- findDupMarkers(perm.combo, adjacent.only = F)
MGnd <- drop.markers(MatGeno, unlist(combo.dups))
PGnd <- drop.markers(PatGeno, unlist(combo.dups))

# How many columns of results?
n.results <- 5
# How many permutations?
n.perms <- 500

#Prepare data for permutation by parent
cross.rows <- row.names(unique(data.frame(MatGeno$pheno$Male_ID, MatGeno$pheno$Female_ID)))
#Make mini cross objects that include each cross only once. 
#mini.G3s.nodups <- subset(G3s.nodups, ind = as.numeric(cross.rows))
mini.MGnd <- subset(MGnd, ind = as.numeric(cross.rows))
#make a mini.Gpat cross object:
mini.PGnd <- subset(MGnd, ind = as.numeric(cross.rows))

perm.scan.holder <- scanone(mini.MGnd, pheno = "nChaetae")
perm.scan.holder$lod <- 0
perm.scan.holder[,3:(2+n.results)] <- 0
perm.results <- matrix(nrow = n.perms, ncol = 2*n.results) # matrix(nrow = n.perms, ncol = 2*n.results)

date()
for(m in 1:n.perms){
	
	male.perm <- data.frame(levels(MatGeno$pheno$Male_ID), levels(MatGeno$pheno$Male_ID))
	# Permutations are stratified by the F1 mother of the F2 fathers, i.e., 1A, 3C, 6F, 8H.
	male.perm[1:10,2] <- sample(male.perm[1:10,1])	
	male.perm[11:24,2] <- sample(male.perm[11:24,1])	
	male.perm[25:32,2] <- sample(male.perm[25:32,1])	
	male.perm[33:53,2] <- sample(male.perm[33:53,1])	

	female.perm <- data.frame(levels(MatGeno$pheno$Female_ID), levels(MatGeno$pheno$Female_ID)) 
	# Permutations are stratified by the F1 mother of the F2 mothers, i.e., 1A, 3C, 6F, 8H.
	female.perm[1:23,2] <- sample(female.perm[1:23,1])	
	female.perm[24:60,2] <- sample(female.perm[24:60,1])		
	female.perm[61:81,2] <- sample(female.perm[61:81,1])	
	female.perm[82:115,2] <- sample(female.perm[82:115,1])	
	
#	perm.G3s <- G3s.nodups
	perm.Gmat <- MGnd
	perm.Gpat <- PGnd
	
	for(i in 1:dim(MatGeno$pheno)[1]){
		permdad <- male.perm[which(male.perm[,1] == MatGeno$pheno$Male_ID[i]),2]
		permmom <- female.perm[which(female.perm[,1] == MatGeno$pheno$Female_ID[i]),2]
		for(j in 1:11){
			perm.Gpat$geno[[j]]$data[i,] <- mini.PGnd$geno[[j]]$data[which(mini.MGnd$pheno$Male_ID == permdad)[1],]
			perm.Gmat$geno[[j]]$data[i,] <- mini.MGnd$geno[[j]]$data[which(mini.MGnd$pheno$Female_ID == permmom)[1],]
		}
	}
	
permscan <- AreaMZscan5(perm.Gmat, perm.Gpat, perm.scan.holder)

perm.results[m,1:n.results] <- apply(as.matrix(permscan[permscan$chr != "X",3:(2+n.results)]), 2, max)
perm.results[m, (n.results+1):(2*n.results)] <- apply(as.matrix(permscan[permscan$chr == "X",3:(2+n.results)]), 2, max)
perm.out <- data.frame(perm.results)
names(perm.out)  <- colnames(permscan)[3:(2+n.results)]
print(m); print(date());
}

perm.out[,11:20] <- NA

perm.out[,11:20] <- cbind(2*(perm.out$AreaBoth - perm.out$AreaNull), 2*(perm.out$AreaSummed - perm.out$AreaNull), 2*(perm.out$AreaBoth - perm.out$AreaSummed), 2*(perm.out$AreaBoth - perm.out$AreaMat), 2*(perm.out$AreaBoth - perm.out$AreaPat), 2*(perm.out[,6] - perm.out$AreaNull), 2*(perm.out[,7] - perm.out$AreaNull), 2*(perm.out[,6] - perm.out[,7]), 2*(perm.out[,6]- perm.out[,8]), 2*(perm.out[,6] - perm.out[,9]))
names(perm.out)[11:20] <- c("BothChisq", "SummedChisq", "BetasChisq","MatVBothChisq", "PatVBothChisq", "XBothChisq", "XSummedChisq", "XBetasChisq","XMatVBothChisq", "XPatVBothChisq")

apply(perm.out[,11:20], 2, function(x){quantile(x,0.95)})

#     BothChisq    SummedChisq     BetasChisq  MatVBothChisq  PatVBothChisq     XBothChisq   XSummedChisq    XBetasChisq 
#     13.779889      10.134564       7.103213       4.895046       9.297969       9.436422       6.651440       4.668213 
#XMatVBothChisq XPatVBothChisq 
#      4.053284       6.995066 

perm.out -> AreaMZScan5Perms
save(AreaMZScan5Perms, file = "AreaScan5.500perms.R")
rm(perm.out)

#No peak exceeds the p=0.05 threshold for significance for the full or additive model genome scans.

############################
#Tests for dominance
 
 	dadID <- MatGeno$pheno$Dad_G2_Name
	momID <-  MatGeno$pheno$Mom_G2_Name
	area.pheno <- MatGeno$pheno$area
	
	# Area QTL are parental-effect loci, so dominance is encoded differently
	qtl1.area <- MatGeno$geno[[7]]$data[,"SNP68"] - 1
	qtl2.area <- MatGeno$geno[[6]]$data[,"SNP93"] - 1
	qtl3.areaP <- PatGeno$geno[["X"]]$data[,"SNP1249"] - 1.  # X-linked and no opportunity for dominance
	qtl3.areaM <- MatGeno$geno[["X"]]$data[,"SNP1249"] - 1.  # X-linked and no opportunity for dominance
	qtl4.areaP <- PatGeno$geno[[9]]$data[,"SNP1244"] - 1
	qtl4.areaM <- MatGeno$geno[[9]]$data[,"SNP1244"] - 1

	qtl1.area.h <- qtl1.area
	qtl1.area.h[qtl1.area == 2] <- 0
	qtl2.area.h <- qtl2.area
	qtl2.area.h[qtl2.area == 2] <- 0
	qtl4.area.hP <- qtl4.areaP
	qtl4.area.hP[qtl4.areaP == 2] <- 0
	qtl4.area.hM <- qtl4.areaM
	qtl4.area.hM[qtl4.areaM == 2] <- 0

area.add <- lmer(area.pheno  ~ 1 + qtl1.area + qtl2.area + qtl3.areaP + qtl3.areaM + qtl4.areaP + qtl4.areaM + (1|momID), REML = F)
area.dom1 <- lmer(area.pheno  ~ 1 + qtl1.area + qtl2.area + qtl3.areaP + qtl3.areaM + qtl4.areaP + qtl4.areaM + qtl1.area.h + (1|momID), REML = F)
area.dom2 <- lmer(area.pheno  ~ 1 + qtl1.area + qtl2.area + qtl3.areaP + qtl3.areaM + qtl4.areaP + qtl4.areaM + qtl2.area.h + (1|momID), REML = F)
area.dom4P <- lmer(area.pheno  ~ 1 + qtl1.area + qtl2.area + qtl3.areaP + qtl3.areaM + qtl4.areaP + qtl4.areaM + qtl4.area.hP + (1|momID), REML = F)
area.dom4M <- lmer(area.pheno  ~ 1 + qtl1.area + qtl2.area + qtl3.areaP + qtl3.areaM + qtl4.areaP + qtl4.areaM + qtl4.area.hM + (1|momID), REML = F)
area.dom24P <- lmer(area.pheno  ~ 1 + qtl1.area + qtl2.area + qtl3.areaP + qtl3.areaM + qtl4.areaP + qtl4.areaM + qtl2.area.h + qtl4.area.hP + (1|momID), REML = F)
area.dom24M <- lmer(area.pheno  ~ 1 + qtl1.area + qtl2.area + qtl3.areaP + qtl3.areaM + qtl4.areaP + qtl4.areaM + qtl2.area.h + qtl4.area.hM + (1|momID), REML = F)
area.full <- lmer(area.pheno  ~ 1 + qtl1.area + qtl2.area + qtl3.areaP + qtl3.areaM + qtl4.areaP + qtl4.areaM + qtl1.area.h + qtl2.area.h + qtl4.area.hP + qtl4.area.hM + (1|momID), REML = F)

anova(area.dom1, area.add)
#area.add: area.pheno ~ 1 + qtl1.area + qtl2.area + qtl3.areaP + qtl3.areaM + qtl4.areaP + qtl4.areaM + (1 | momID)
#area.dom1: area.pheno ~ 1 + qtl1.area + qtl2.area + qtl3.areaP + qtl3.areaM + qtl4.areaP + qtl4.areaM + qtl1.area.h + (1 | momID)
#          npar   AIC   BIC logLik deviance  Chisq Df Pr(>Chisq)
#area.add     9 78588 78644 -39285    78570                     
#area.dom1   10 78589 78652 -39284    78569 0.7498  1     0.3866

anova(area.dom2, area.add)
#area.add: area.pheno ~ 1 + qtl1.area + qtl2.area + qtl3.areaP + qtl3.areaM + qtl4.areaP + qtl4.areaM + (1 | momID)
#area.dom2: area.pheno ~ 1 + qtl1.area + qtl2.area + qtl3.areaP + qtl3.areaM + qtl4.areaP + qtl4.areaM + qtl2.area.h + (1 | momID)
#          npar   AIC   BIC logLik deviance  Chisq Df Pr(>Chisq)  
#area.add     9 78588 78644 -39285    78570                       
#area.dom2   10 78586 78648 -39283    78566 3.9221  1    0.04766 *

anova(area.dom24P, area.dom2)
#area.dom2: area.pheno ~ 1 + qtl1.area + qtl2.area + qtl3.areaP + qtl3.areaM + qtl4.areaP + qtl4.areaM + qtl2.area.h + (1|momID)
#area.dom24P: area.pheno ~ 1 + qtl1.area + qtl2.area + qtl3.areaP + qtl3.areaM + qtl4.areaP + qtl4.areaM + qtl2.area.h + qtl4.area.hP + (1|momID)
#            npar   AIC   BIC logLik deviance  Chisq Df Pr(>Chisq)
#area.dom2     10 78586 78648 -39283    78566                     
#area.dom24P   11 78587 78656 -39283    78565 0.3036  1     0.5816

anova(area.dom24M, area.dom2)
#area.dom2: area.pheno ~ 1 + qtl1.area + qtl2.area + qtl3.areaP + qtl3.areaM + qtl4.areaP + qtl4.areaM + qtl2.area.h + (1|momID)
#area.dom24M: area.pheno ~ 1 + qtl1.area + qtl2.area + qtl3.areaP + qtl3.areaM + qtl4.areaP + qtl4.areaM + qtl2.area.h + qtl4.area.hM + (1|momID)
#            npar   AIC   BIC logLik deviance Chisq Df Pr(>Chisq)  
#area.dom2     10 78586 78648 -39283    78566                      
#area.dom24M   11 78584 78653 -39281    78562 3.503  1    0.06126 .

# Model:
#area.pheno ~ 1 + qtl1.area + qtl2.area + qtl3.areaP + qtl3.areaM + qtl4.areaP + qtl4.areaM + qtl2.area.h + (1|momID)


############################
# Tests for polygenic effects
# Autosomes only
get.lecitho.proportion <- function(genotypes, chromosomes = c(1:10)){ 
allelecountbychr <- matrix(nrow = dim(genotypes$pheno)[1], ncol = length(chromosomes))	
		for(j in 1: length(chromosomes)){
		allelecountbychr[,j] <- apply(genotypes$geno[[j]]$data -1, 1, sum)
		}
TotalAlleleCount <- apply(allelecountbychr, 1, sum)
TotalAlleleCount/(2*sum(nmar(genotypes)[c(chromosomes)]))
}

area.dom2 <- lmer(area.pheno  ~ 1 + qtl1.area + qtl2.area + qtl3.areaP + qtl3.areaM + qtl4.areaP + qtl4.areaM + qtl2.area.h + (1|momID), REML = F)

LpropM.area <- get.lecitho.proportion(MatGeno, c(1:5,8,10)) # excludes 6, 7, 9, and X
LpropP.area <- get.lecitho.proportion(PatGeno, c(1:8,10)) # excludes 9 and X

MatLprop <- get.lecitho.proportion(MatGeno)
PatLprop <- get.lecitho.proportion(PatGeno)
ZygLprop <- (MatLprop+PatLprop)/2 # includes all the autosomes

area.polyMx <- lmer(area.pheno  ~ 1 + qtl1.area + qtl2.area + qtl3.areaP + qtl3.areaM + qtl4.areaP + qtl4.areaM + qtl2.area.h + LpropM.area + (1|momID), REML = F)
area.polyPx <- lmer(area.pheno  ~ 1 + qtl1.area + qtl2.area + qtl3.areaP + qtl3.areaM + qtl4.areaP + qtl4.areaM + qtl2.area.h + LpropP.area + (1|momID), REML = F)
area.polyZ <- lmer(area.pheno  ~ 1 + qtl1.area + qtl2.area + qtl3.areaP + qtl3.areaM + qtl4.areaP + qtl4.areaM + qtl2.area.h + ZygLprop + (1|momID), REML = F)

anova(area.polyMx, area.dom2)
#area.dom2: area.pheno ~ 1 + qtl1.area + qtl2.area + qtl3.areaP + qtl3.areaM + qtl4.areaP + qtl4.areaM + qtl2.area.h + (1 | momID)
#area.polyMx: area.pheno ~ 1 + qtl1.area + qtl2.area + qtl3.areaP + qtl3.areaM + qtl4.areaP + qtl4.areaM + qtl2.area.h + LpropM.area + (1|momID)
#            npar   AIC   BIC logLik deviance  Chisq Df Pr(>Chisq)  
#area.dom2     10 78586 78648 -39283    78566                       
#area.polyMx   11 78583 78652 -39281    78561 4.3347  1    0.03734 *

anova(area.polyPx, area.dom2)
#area.dom2: area.pheno ~ 1 + qtl1.area + qtl2.area + qtl3.areaP + qtl3.areaM + qtl4.areaP + qtl4.areaM + qtl2.area.h + (1 | momID)
#area.polyPx: area.pheno ~ 1 + qtl1.area + qtl2.area + qtl3.areaP + qtl3.areaM + qtl4.areaP + qtl4.areaM + qtl2.area.h + LpropP.area + (1|momID)
#            npar   AIC   BIC logLik deviance  Chisq Df Pr(>Chisq)
#area.dom2     10 78586 78648 -39283    78566                     
#area.polyPx   11 78586 78655 -39282    78564 1.5203  1     0.2176

anova(area.polyZ, area.dom2)
#area.dom2: area.pheno ~ 1 + qtl1.area + qtl2.area + qtl3.areaP + qtl3.areaM + qtl4.areaP + qtl4.areaM + qtl2.area.h + (1 | momID)
#area.polyZ: area.pheno ~ 1 + qtl1.area + qtl2.area + qtl3.areaP + qtl3.areaM + qtl4.areaP + qtl4.areaM + qtl2.area.h + ZygLprop + (1 | momID)
#           npar   AIC   BIC logLik deviance  Chisq Df Pr(>Chisq)
#area.dom2    10 78586 78648 -39283    78566                     
#area.polyZ   11 78586 78655 -39282    78564 2.0798  1     0.1493

# Quadratic or cubic effects
area.polyMx2 <- lmer(area.pheno  ~ 1 + qtl1.area + qtl2.area + qtl3.areaP + qtl3.areaM + qtl4.areaP + qtl4.areaM + qtl2.area.h + poly(LpropM.area, 2, raw = T) + (1|momID), REML = F)
area.polyMx3 <- lmer(area.pheno  ~ 1 + qtl1.area + qtl2.area + qtl3.areaP + qtl3.areaM + qtl4.areaP + qtl4.areaM + qtl2.area.h + poly(LpropM.area, 3, raw = T) + (1|momID), REML = F)
# no improvement over additive polygenic effect
area.polyPx2 <- lmer(area.pheno  ~ 1 + qtl1.area + qtl2.area + qtl3.areaP + qtl3.areaM + qtl4.areaP + qtl4.areaM + qtl2.area.h + poly(LpropP.area, 2, raw = T) + (1|momID), REML = F)
area.polyPx3 <- lmer(area.pheno  ~ 1 + qtl1.area + qtl2.area + qtl3.areaP + qtl3.areaM + qtl4.areaP + qtl4.areaM + qtl2.area.h + poly(LpropP.area, 3, raw = T) + (1|momID), REML = F)
# no improvement over no polygenic effect
area.polyZ2 <- lmer(area.pheno  ~ 1 + qtl1.area + qtl2.area + qtl3.areaP + qtl3.areaM + qtl4.areaP + qtl4.areaM + qtl2.area.h + poly(ZygLprop, 2, raw = T) + (1|momID), REML = F)
area.polyZ3 <- lmer(area.pheno  ~ 1 + qtl1.area + qtl2.area + qtl3.areaP + qtl3.areaM + qtl4.areaP + qtl4.areaM + qtl2.area.h + poly(ZygLprop, 3, raw = T) + (1|momID), REML = F)
# no improvement over no polygenic effect


############################
# Tests for Epistasis

# no-interaction null model
area.polyMx <- lmer(area.pheno  ~ qtl1.area + qtl2.area + qtl3.areaP + qtl3.areaM + qtl4.areaP + qtl4.areaM + qtl2.area.h + LpropM.area + (1|momID), REML = F)

# all pairwise additive-by-additive epistasis:
Pairwise.area.polyMx <- lmer(area.pheno  ~ (qtl1.area + qtl2.area + qtl3.areaP + qtl3.areaM + qtl4.areaP + qtl4.areaM)^2 + qtl2.area.h + LpropM.area + (1|momID), REML = F)

anova(area.polyMx, Pairwise.area.polyMx)
#                    npar   AIC   BIC logLik deviance  Chisq Df Pr(>Chisq)
#area.polyMx            11 78583 78652 -39281    78561                     
#Pairwise.area.polyMx   26 78596 78759 -39272    78544 17.249 15     0.3042

# intraparental pairwise additive-by-additive epistasis
Parental.Pairwise.area.polyMx <- lmer(area.pheno  ~ (qtl1.area + qtl2.area + qtl3.areaM + qtl4.areaM)^2+ (qtl3.areaP + qtl4.areaP)^2 + qtl2.area.h + LpropM.area + (1|momID), REML = F)

anova(Parental.Pairwise.area.polyMx, area.polyMx)
#                              npar   AIC   BIC logLik deviance Chisq Df Pr(>Chisq)
#area.polyMx                     11 78583 78652 -39281    78561                    
#Parental.Pairwise.area.polyMx   18 78590 78703 -39277    78554 6.999  7      0.429

# maternal pairwise additive-by-additive epistasis:
Maternal.Pairwise.area.polyMx <- lmer(area.pheno  ~ (qtl1.area + qtl2.area + qtl3.areaM + qtl4.areaM)^2+ qtl3.areaP + qtl4.areaP + qtl2.area.h + LpropM.area + (1|momID), REML = F)

anova(Maternal.Pairwise.area.polyMx, area.polyMx)
#                              npar   AIC   BIC logLik deviance  Chisq Df Pr(>Chisq)
#area.polyMx                     11 78583 78652 -39281    78561                     
#Maternal.Pairwise.area.polyMx   17 78592 78699 -39279    78558 2.9102  6       0.82

# paternal pairwise additive-by-additive epistasis:
Paternal.Pairwise.area.polyMx <- lmer(area.pheno  ~ qtl1.area + qtl2.area + qtl3.areaM + qtl4.areaM+ (qtl3.areaP + qtl4.areaP)^2 + qtl2.area.h + LpropM.area + (1|momID), REML = F)

anova(Paternal.Pairwise.area.polyMx, area.polyMx)
#                              npar   AIC   BIC logLik deviance  Chisq Df Pr(>Chisq)  
#area.polyMx                     11 78583 78652 -39281    78561                       
#Paternal.Pairwise.area.polyMx   12 78582 78658 -39279    78558 3.0526  1    0.08061 .

# Interaction between the major-effect maternal QTL
E1x2.area.polyMx <- lmer(area.pheno  ~ qtl1.area * qtl2.area + qtl3.areaP + qtl3.areaM + qtl4.areaP + qtl4.areaM + qtl2.area.h + LpropM.area + (1|momID), REML = F)

anova(E1x2.area.polyMx, area.polyMx)
#                 npar   AIC   BIC logLik deviance  Chisq Df Pr(>Chisq)
#area.polyMx        11 78583 78652 -39281    78561                     
#E1x2.area.polyMx   12 78585 78660 -39280    78561 0.5722  1     0.4494

#Interactions between parents, QTL on X:
PMX.area.polyMx <- lmer(area.pheno  ~ qtl1.area + qtl2.area + qtl3.areaP * qtl3.areaM + qtl4.areaP + qtl4.areaM + qtl2.area.h + LpropM.area + (1|momID), REML = F)
 
anova(PMX.area.polyMx, area.polyMx)
#                npar   AIC   BIC logLik deviance  Chisq Df Pr(>Chisq)
#area.polyMx       11 78583 78652 -39281    78561                     
#PMX.area.polyMx   12 78585 78660 -39281    78561 0.1059  1     0.7449


#Interactions between parents, QTL on 9:
PM9.area.polyMx <- lmer(area.pheno  ~ qtl1.area + qtl2.area + qtl3.areaP + qtl3.areaM + qtl4.areaP * qtl4.areaM + qtl2.area.h + LpropM.area + (1|momID), REML = F)
 
anova(PM9.area.polyMx, area.polyMx)
#                npar   AIC   BIC logLik deviance  Chisq Df Pr(>Chisq)
#area.polyMx       11 78583 78652 -39281    78561                     
#PM9.area.polyMx   12 78583 78659 -39280    78559 1.8702  1     0.1715


# Final model
summary(area.polyMx)

###############################
# Tests for direct effects of the M+P loci in the G2 generation

Sb.slim <- subset(Sb.deduped, ind = !is.na(Sb.deduped$pheno$G2.area))
# Data for 240 G2 larvae

# Locus on the X chromosome. 
# Must include terms for sex and sex-by-genotype because males are LY and BY and females are LB and BB. 
xqtl <- Sb.slim $geno[["X"]]$data[,"SNP1249"] -1
sexXqtl <- lmer(Sb.slim$pheno$G2.area ~ xqtl* Sb.slim$pheno$Sex + (1| Sb.slim$pheno$Family))
sexonly <- lmer(Sb.slim$pheno$G2.area ~ Sb.slim$pheno$Sex + (1| Sb.slim$pheno$Family))

anova(sexXqtl, sexonly)
#refitting model(s) with ML (instead of REML)
#        npar    AIC    BIC  logLik deviance  Chisq Df Pr(>Chisq)
#sexonly    4 4944.1 4958.1 -2468.1   4936.1                     
#sexXqtl    6 4947.9 4968.8 -2467.9   4935.9 0.2559  2     0.8799

# No evidence of any direct (zygotic) effect of the X-linked locus

# Locus on chromosome 9
qtl9 <- Sb.slim$geno[[9]]$data[,"SNP1244"] - 1
withqtl9 <- lmer(Sb.slim$pheno$G2.area ~ qtl9 + (1| Sb.slim$pheno$Family))
null9 <- lmer(Sb.slim$pheno$G2.area ~ 1 + (1| Sb.slim$pheno$Family))
anova(withqtl9, null9)
#refitting model(s) with ML (instead of REML)
#         npar    AIC    BIC  logLik deviance  Chisq Df Pr(>Chisq)
#null9       3 4944.6 4955.1 -2469.3   4938.6                     
#withqtl9    4 4945.1 4959.0 -2468.5   4937.1 1.5646  1      0.211

# No evidence of any direct (zygotic) effect of the locus on chromosome 9
```
