## Supplementary File 1. Data and Code for "Baby makes three: maternal, paternal, and zygotic genetic effects shape larval phenotypic evolution": ChaetaeAndCirriTestsForDominanceAndEpistasis.R

```
# Tests for QTL dominance and epistasis using expected heterozygosity of G3 broods
# Cirri and Chaetae traits

load(file = "G3AnalysisObjects.R")
# [1] "dups"       "MatGeno"    "PatGeno"    "Sb.deduped"

library(qtl)
library(lme4)

# function to get expected G3 heterozygosity from the genotypes of G2 parents.
get.expected.heterozygosity <- function(geno.mat, geno.pat, chrom, snpname){
	snp.M <- geno.mat$geno[[chrom]]$data[,snpname] - 2
	snp.P <- geno.pat$geno[[chrom]]$data[,snpname] - 2
	snp <- snp.M + snp.P
	snp.het <- snp
	snp.het[snp %in% c(-2,2)] <- 0
	snp.het[snp.M == 0] <- 0.5
	snp.het[snp.P == 0] <- 0.5
	snp.het[snp.M == -1 & snp.P == 1] <- 1
	snp.het[snp.M == 1 & snp.P == -1] <- 1
	return(snp.het)
}	

	dadID <- MatGeno$pheno$Dad_G2_Name
	momID <-  MatGeno$pheno$Mom_G2_Name
	chnum.bin.pheno <- MatGeno$pheno$nChaetae
	chnum.bin.pheno[chnum.bin.pheno > 0] <- 1
	bc.chnum0 <- ((MatGeno$pheno$nChaetae^0.29)-1)/0.29 # uses the maximum likelihood Box-Cox coefficient for these data
	bc.chnum0[MatGeno$pheno$nChaetae == 0] <- NA

	# QTLs for chaetae and anal cirri
	qtl1.chlen <- MatGeno$geno[[9]]$data[,"SNP766"] + PatGeno$geno[[9]]$data[,"SNP766"] - 2
	qtl2.chlen <- MatGeno$geno[[3]]$data[,"SNP555"] + PatGeno$geno[[3]]$data[,"SNP555"] - 2
	qtl1.chnum.bin <- MatGeno$geno[[8]]$data[,"SNP230"] + PatGeno$geno[[8]]$data[,"SNP230"] - 2
	qtl1.bc.chnum0 <- MatGeno$geno[[8]]$data[,"SNP768"] + PatGeno$geno[[8]]$data[,"SNP768"] - 2
	qtl2.chnum <- MatGeno$geno[[3]]$data[,"SNP779"] + PatGeno$geno[[3]]$data[,"SNP779"] - 2
	qtl1.chnum.combined <- MatGeno$geno[[8]]$data[,"SNP725"] + PatGeno$geno[[8]]$data[,"SNP725"] - 2
	qtl1.cirri <- MatGeno$geno[[5]]$data[,"SNP1072"] +  PatGeno$geno[[5]]$data[,"SNP1072"] - 2

# now fit the models 

# ANAL CIRRI
cirri.h <- get.expected.heterozygosity(MatGeno, PatGeno, 5, "SNP1072")
cirri.add <- glmer(MatGeno$pheno$cirri ~  qtl1.cirri + (1|momID), family = "binomial")
cirri.dom <- glmer(MatGeno$pheno$cirri ~  qtl1.cirri + cirri.h + (1|momID), family = "binomial")

anova(cirri.dom, cirri.add)
#Models:
#cirri.add: MatGeno$pheno$cirri ~ qtl1.cirri + (1 | momID)
#cirri.dom: MatGeno$pheno$cirri ~ qtl1.cirri + cirri.h + (1 | momID)
#          npar    AIC    BIC  logLik deviance Chisq Df Pr(>Chisq)  
#cirri.add    3 4072.5 4091.3 -2033.2   4066.5                      
#cirri.dom    4 4071.7 4096.8 -2031.8   4063.7 2.784  1    0.09521 .

# CHAETAE LENGTH
chlenqtl1.h <- get.expected.heterozygosity(MatGeno, PatGeno, 9, "SNP766")
chlenqtl2.h <- get.expected.heterozygosity(MatGeno, PatGeno, 3, "SNP555")
chlen.add <- lmer(MatGeno$pheno$ChLen ~ qtl1.chlen + qtl2.chlen + (1|dadID) + (1|momID), REML = F)
chlen.dom1 <- lmer(MatGeno$pheno$ChLen ~ qtl1.chlen + qtl2.chlen + chlenqtl1.h + (1|dadID) + (1|momID), REML = F)
chlen.dom2 <- lmer(MatGeno$pheno$ChLen ~ qtl1.chlen + qtl2.chlen + chlenqtl2.h + (1|dadID) + (1|momID), REML = F)
chlen.dom12 <- lmer(MatGeno$pheno$ChLen ~ qtl1.chlen + qtl2.chlen + chlenqtl1.h + chlenqtl2.h + (1|dadID) + (1|momID), REML = F)

anova(chlen.dom1, chlen.add)
#Models:
#chlen.add: MatGeno$pheno$ChLen ~ qtl1.chlen + qtl2.chlen + (1 | dadID) + (1 | momID)
#chlen.dom1: MatGeno$pheno$ChLen ~ qtl1.chlen + qtl2.chlen + chlenqtl1.h + (1 | dadID) + (1 | momID)
#           npar   AIC   BIC logLik deviance  Chisq Df Pr(>Chisq)  
#chlen.add     6 29091 29127 -14540    29079                       
#chlen.dom1    7 29090 29132 -14538    29076 2.9678  1    0.08494 .

anova(chlen.dom2, chlen.add)
#chlen.add: MatGeno$pheno$ChLen ~ qtl1.chlen + qtl2.chlen + (1 | dadID) + (1 | momID)
#chlen.dom2: MatGeno$pheno$ChLen ~ qtl1.chlen + qtl2.chlen + chlenqtl2.h + (1 | dadID) + (1 | momID)
#           npar   AIC   BIC logLik deviance  Chisq Df Pr(>Chisq)   
#chlen.add     6 29091 29127 -14540    29079                        
#chlen.dom2    7 29086 29128 -14536    29072 7.2755  1    0.00699 **

# CHAETAE NUMBER - COMBINED 
chnumqtl1.h <- get.expected.heterozygosity(MatGeno, PatGeno, 8, "SNP725")
chnumqtl2.h <- get.expected.heterozygosity(MatGeno, PatGeno, 3, "SNP779")

chnum.combined.add <- logLik(glmer(chnum.bin.pheno ~ qtl1.chnum.combined + qtl2.chnum + (1|momID), family = "binomial"))[1] + logLik(lmer(bc.chnum0 ~ qtl1.chnum.combined + qtl2.chnum + (1|dadID) + (1|momID), REML = F))[1]
chnum.combined.dom1 <- logLik(glmer(chnum.bin.pheno ~ qtl1.chnum.combined + qtl2.chnum + chnumqtl1.h + (1|momID), family = "binomial"))[1] + logLik(lmer(bc.chnum0 ~ qtl1.chnum.combined + qtl2.chnum + chnumqtl1.h + (1|dadID) + (1|momID), REML = F))[1]
chnum.combined.dom2 <- logLik(glmer(chnum.bin.pheno ~ qtl1.chnum.combined + qtl2.chnum + chnumqtl2.h + (1|momID), family = "binomial"))[1] + logLik(lmer(bc.chnum0 ~ qtl1.chnum.combined + qtl2.chnum + chnumqtl2.h + (1|dadID) + (1|momID), REML = F))[1]
chnum.combined.dom12 <- logLik(glmer(chnum.bin.pheno ~ qtl1.chnum.combined + qtl2.chnum + chnumqtl1.h + chnumqtl2.h + (1|momID), family = "binomial"))[1] + logLik(lmer(bc.chnum0 ~ qtl1.chnum.combined + qtl2.chnum + chnumqtl1.h + chnumqtl2.h + (1|dadID) + (1|momID), REML = F))[1]

pchisq(2*(chnum.combined.dom1 - chnum.combined.add), 2, lower = F)
# [1] 0.0001378026
pchisq(2*(chnum.combined.dom12 - chnum.combined.dom1), 2, lower = F)
# [1] 0.02528473

# CHAETAE NUMBER - count 

chnum.bc.add <- lmer(bc.chnum0 ~ qtl1.chnum.combined + qtl2.chnum + (1|dadID) + (1|momID), REML = F)
chnum.bc.dom1 <- lmer(bc.chnum0 ~ qtl1.chnum.combined + qtl2.chnum + chnumqtl1.h + (1|dadID) + (1|momID), REML = F)
chnum.bc.dom2 <- lmer(bc.chnum0 ~ qtl1.chnum.combined + qtl2.chnum + chnumqtl2.h + (1|dadID) + (1|momID), REML = F)
chnum.bc.dom12 <- lmer(bc.chnum0 ~ qtl1.chnum.combined + qtl2.chnum + chnumqtl1.h + chnumqtl2.h + (1|dadID) + (1|momID), REML = F)

anova(chnum.bc.dom1, chnum.bc.add)
#chnum.bc.add: bc.chnum0 ~ qtl1.chnum.combined + qtl2.chnum + (1 | dadID) + (1 | momID)
#chnum.bc.dom1: bc.chnum0 ~ qtl1.chnum.combined + qtl2.chnum + chnumqtl1.h + (1 | dadID) + (1 | momID)
#              npar    AIC    BIC  logLik deviance  Chisq Df Pr(>Chisq)
#chnum.bc.add     6 8376.1 8412.0 -4182.0   8364.1                     
#chnum.bc.dom1    7 8375.8 8417.7 -4180.9   8361.8 2.3055  1     0.1289

anova(chnum.bc.dom2, chnum.bc.add)
#chnum.bc.add: bc.chnum0 ~ qtl1.chnum.combined + qtl2.chnum + (1 | dadID) + (1 | momID)
#chnum.bc.dom2: bc.chnum0 ~ qtl1.chnum.combined + qtl2.chnum + chnumqtl2.h + (1 | dadID) + (1 | momID)
#              npar    AIC  BIC logLik deviance  Chisq Df Pr(>Chisq)
#chnum.bc.add     6 8376.1 8412  -4182   8364.1                     
#chnum.bc.dom2    7 8378.1 8420  -4182   8364.1 0.0041  1     0.9488


# CHAETAE NUMBER - presence 

chnum.bin.add <- glmer(chnum.bin.pheno ~ qtl1.chnum.combined + qtl2.chnum + (1|momID), family = "binomial")
chnum.bin.dom1 <- glmer(chnum.bin.pheno ~ qtl1.chnum.combined + qtl2.chnum + chnumqtl1.h + (1|momID), family = "binomial")
chnum.bin.dom2 <- glmer(chnum.bin.pheno ~ qtl1.chnum.combined + qtl2.chnum + chnumqtl2.h + (1|momID), family = "binomial")
chnum.bin.dom12 <- glmer(chnum.bin.pheno ~ qtl1.chnum.combined + qtl2.chnum + chnumqtl1.h + chnumqtl2.h + (1|momID), family = "binomial")

anova(chnum.bin.dom1, chnum.bin.add)
#chnum.bin.add: chnum.bin.pheno ~ qtl1.chnum.combined + qtl2.chnum + (1 | momID)
#chnum.bin.dom1: chnum.bin.pheno ~ qtl1.chnum.combined + qtl2.chnum + chnumqtl1.h + (1 | momID)
#               npar    AIC    BIC  logLik deviance  Chisq Df Pr(>Chisq)    
#chnum.bin.add     4 3669.9 3695.0 -1830.9   3661.9                         
#chnum.bin.dom1    5 3656.4 3687.8 -1823.2   3646.4 15.474  1  8.365e-05 ***

anova(chnum.bin.dom12, chnum.bin.dom1)
#chnum.bin.dom1: chnum.bin.pheno ~ qtl1.chnum.combined + qtl2.chnum + chnumqtl1.h + (1 | momID)
#chnum.bin.dom12: chnum.bin.pheno ~ qtl1.chnum.combined + qtl2.chnum + chnumqtl1.h + chnumqtl2.h + (1 | momID)
#                npar    AIC    BIC  logLik deviance  Chisq Df Pr(>Chisq)   
#chnum.bin.dom1     5 3656.4 3687.8 -1823.2   3646.4                        
#chnum.bin.dom12    6 3651.1 3688.8 -1819.5   3639.1 7.3289  1   0.006785 **

##################################

# Tests for epistasis
chlen.dom2e <- lmer(MatGeno$pheno$ChLen ~ qtl1.chlen * (qtl2.chlen + chlenqtl2.h) + (1|dadID) + (1|momID), REML = F)
anova(chlen.dom2e, chlen.dom2)
#            npar   AIC   BIC logLik deviance  Chisq Df Pr(>Chisq)
#chlen.dom2     7 29086 29128 -14536    29072                     
#chlen.dom2e    9 29089 29143 -14535    29071 1.2561  2     0.5336

chnum.bc.e <- lmer(bc.chnum0 ~ qtl1.chnum.combined * qtl2.chnum + (1|dadID) + (1|momID), REML = F)
anova(chnum.bc.e, chnum.bc.add)
#             npar    AIC    BIC  logLik deviance  Chisq Df Pr(>Chisq)  
#chnum.bc.add    6 8376.1 8412.0 -4182.0   8364.1                       
#chnum.bc.e      7 8373.1 8415.1 -4179.5   8359.1 4.9786  1    0.02566 *

chnum.bin.dom12e <- glmer(chnum.bin.pheno ~ (qtl1.chnum.combined + chnumqtl1.h )* (qtl2.chnum + chnumqtl2.h) + (1|momID), family = "binomial")
anova(chnum.bin.dom12e, chnum.bin.dom12)
#                 npar    AIC    BIC  logLik deviance  Chisq Df Pr(>Chisq)
#chnum.bin.dom12     6 3651.1 3688.8 -1819.5   3639.1                     
#chnum.bin.dom12e   10 3653.1 3715.9 -1816.5   3633.1 5.9988  4     0.1992

chnum.combined.epi.LL <- logLik(glmer(chnum.bin.pheno ~ (qtl1.chnum.combined + chnumqtl1.h )* (qtl2.chnum + chnumqtl2.h) + (1|momID), family = "binomial"))[1] + logLik(lmer(bc.chnum0 ~ qtl1.chnum.combined * qtl2.chnum + (1|dadID) + (1|momID), REML = F))[1]
[1] -5996.073
chnum.combined.noepi.LL <- logLik(glmer(chnum.bin.pheno ~ qtl1.chnum.combined + chnumqtl1.h  + qtl2.chnum + chnumqtl2.h + (1|momID), family = "binomial"))[1] + logLik(lmer(bc.chnum0 ~ qtl1.chnum.combined + qtl2.chnum + (1|dadID) + (1|momID), REML = F))[1]
[1] -6001.562
# -2(LL.epi - LL.noepi) = 5.489, 5 df, p = 0.36.

##################################
# Favored models
cirri.add <- glmer(MatGeno$pheno$cirri ~  qtl1.cirri + (1|momID), family = "binomial")
chlen.dom2 <- lmer(MatGeno$pheno$ChLen ~ qtl1.chlen + qtl2.chlen + chlenqtl2.h + (1|dadID) + (1|momID), REML = F)
chnum.bc.e <- lmer(bc.chnum0 ~ qtl1.chnum.combined * qtl2.chnum + (1|dadID) + (1|momID), REML = F)
chnum.bin.dom12 <- glmer(chnum.bin.pheno ~ qtl1.chnum.combined + qtl2.chnum + chnumqtl1.h + chnumqtl2.h + (1|momID), family = "binomial")
```
