## Supplementary File 1. Data and Code for "Baby makes three: maternal, paternal, and zygotic genetic effects shape larval phenotypic evolution": ChaetaeAndCirriTestsForPolygenicEffects.R

```
# Tests for polygenic background effects

load(file = "G3AnalysisObjects.R")
# [1] "dups"       "MatGeno"    "PatGeno"    "Sb.deduped"

library(qtl)
library(lme4)

get.expected.heterozygosity <- function(geno.mat, geno.pat, chrom, snpname){
	snp.M <- geno.mat$geno[[chrom]]$data[,snpname] - 2
	snp.P <- geno.pat$geno[[chrom]]$data[,snpname] - 2
	snp <- snp.M + snp.P
	snp.het <- snp
	snp.het[snp %in% c(-2,2)] <- 0
	snp.het[snp.M == 0] <- 0.5
	snp.het[snp.P == 0] <- 0.5
	snp.het[snp.M == -1 & snp.P == 1] <- 1
	snp.het[snp.M == 1 & snp.P == -1] <- 1
	return(snp.het)
}	

	qtl1.chlen <- MatGeno$geno[[9]]$data[,"SNP766"] + PatGeno$geno[[9]]$data[,"SNP766"] - 2
	qtl1.chnum.bin <- MatGeno$geno[[8]]$data[,"SNP230"] + PatGeno$geno[[8]]$data[,"SNP230"] - 2
	qtl1.bc.chnum0 <- MatGeno$geno[[8]]$data[,"SNP768"] + PatGeno$geno[[8]]$data[,"SNP768"] - 2
	qtl1.chnum.combined <- MatGeno$geno[[8]]$data[,"SNP725"] + PatGeno$geno[[8]]$data[,"SNP725"] - 2
	qtl1.cirri <- MatGeno$geno[[5]]$data[,"SNP1072"] +  PatGeno$geno[[5]]$data[,"SNP1072"] - 2

	qtl2.chlen <- MatGeno$geno[[3]]$data[,"SNP555"] + PatGeno$geno[[3]]$data[,"SNP555"] - 2
	chlenqtl2.h <- get.expected.heterozygosity(MatGeno, PatGeno, 3, "SNP555")

	qtl2.chnum <- MatGeno$geno[[3]]$data[,"SNP779"] + PatGeno$geno[[3]]$data[,"SNP779"] - 2
	chnumqtl1.h <- get.expected.heterozygosity(MatGeno, PatGeno, 8, "SNP725")
	chnumqtl2.h <- get.expected.heterozygosity(MatGeno, PatGeno, 3, "SNP779")
	
	# Area QTL are parental-effect loci, so dominance is encoded differently
	qtl1.area <- MatGeno$geno[[7]]$data[,"SNP68"] - 1
	qtl2.area <- MatGeno$geno[[6]]$data[,"SNP93"] - 1
	qtl3.area <- PatGeno$geno[["X"]]$data[,"SNP1249"] - 1.  
	qtl4.area <- PatGeno$geno[[9]]$data[,"SNP1244"] - 1
	qtl1.area.h <- qtl1.area
	qtl1.area.h[qtl1.area == 2] <- 0
	qtl2.area.h <- qtl2.area
	qtl2.area.h[qtl2.area == 2] <- 0
	qtl4.area.h <- qtl4.area
	qtl4.area.h[qtl4.area == 2] <- 0


	area.pheno <- MatGeno$pheno$area
	chnum.bin.pheno <- MatGeno$pheno$nChaetae
	chnum.bin.pheno[chnum.bin.pheno > 0] <- 1
	bc.chnum0 <- ((MatGeno$pheno$nChaetae^0.29)-1)/0.29 # uses the maximum likelihood Box-Cox coefficient for these data
	bc.chnum0[MatGeno$pheno$nChaetae == 0] <- NA
	dadID <- MatGeno$pheno$Dad_G2_Name
	momID <-  MatGeno$pheno$Mom_G2_Name

# Function to estimate the proportion of genome carrying lecithotroph alleles
get.lecitho.proportion <- function(genotypes, chromosomes = c(1:10)){ 
allelecountbychr <- matrix(nrow = dim(genotypes$pheno)[1], ncol = length(chromosomes))	
		for(j in 1: length(chromosomes)){
		allelecountbychr[,j] <- apply(genotypes$geno[[j]]$data -1, 1, sum)
		}
TotalAlleleCount <- apply(allelecountbychr, 1, sum)
TotalAlleleCount/(2*sum(nmar(genotypes)[c(chromosomes)]))
}

MatLprop <- get.lecitho.proportion(MatGeno)
PatLprop <- get.lecitho.proportion(PatGeno)
ZygLprop <- (MatLprop+PatLprop)/2


summary(MatLprop)
#   Min. 1st Qu.  Median    Mean 3rd Qu.    Max. 
# 0.1867  0.3996  0.4659  0.4768  0.5462  0.7410 
summary(PatLprop)
#   Min. 1st Qu.  Median    Mean 3rd Qu.    Max. 
# 0.3012  0.4177  0.4699  0.4907  0.5663  0.6948 
summary(ZygLprop)
#   Min. 1st Qu.  Median    Mean 3rd Qu.    Max. 
# 0.3283  0.4317  0.4839  0.4837  0.5361  0.6476 

# QTL models:
cirri.add <- glmer(MatGeno$pheno$cirri ~  qtl1.cirri + (1|momID), family = "binomial")
chlen.dom2 <- lmer(MatGeno$pheno$ChLen ~ qtl1.chlen + qtl2.chlen + chlenqtl2.h + (1|dadID) + (1|momID), REML = F)
chnum.bc.add <- lmer(bc.chnum0 ~ qtl1.chnum.combined + qtl2.chnum + (1|dadID) + (1|momID), REML = F)
chnum.bin.dom12 <- glmer(chnum.bin.pheno ~ qtl1.chnum.combined + qtl2.chnum + chnumqtl1.h + chnumqtl2.h + (1|momID), family = "binomial")

# CIRRI 
# additive QTL on chromosome 5
cirri.add <- glmer(MatGeno$pheno$cirri ~  qtl1.cirri + (1|momID), family = "binomial")

LpropM.cirri <- get.lecitho.proportion(MatGeno, c(1:4,6:10))
LpropP.cirri <- get.lecitho.proportion(PatGeno, c(1:4,6:10))
LpropZ.cirri <- (LpropM.cirri + LpropP.cirri)/2

cirri.polyZ <- glmer(MatGeno$pheno$cirri ~  qtl1.cirri + LpropZ.cirri + (1|momID), family = "binomial")
cirri.polyM <- glmer(MatGeno$pheno$cirri ~  qtl1.cirri + LpropM.cirri + (1|momID), family = "binomial")
cirri.polyP <- glmer(MatGeno$pheno$cirri ~  qtl1.cirri + LpropP.cirri + (1|momID), family = "binomial")

anova(cirri.polyZ, cirri.add)
#            npar    AIC    BIC  logLik deviance  Chisq Df Pr(>Chisq)
#cirri.add      3 4072.5 4091.3 -2033.2   4066.5                     
#cirri.polyZ    4 4073.1 4098.2 -2032.5   4065.1 1.4108  1     0.2349

anova(cirri.polyM, cirri.add)
#            npar    AIC    BIC  logLik deviance Chisq Df Pr(>Chisq)
#cirri.add      3 4072.5 4091.3 -2033.2   4066.5                    
#cirri.polyM    4 4074.5 4099.6 -2033.2   4066.5 0.001  1     0.9751

anova(cirri.polyP, cirri.add)
#            npar    AIC    BIC  logLik deviance  Chisq Df Pr(>Chisq)
#cirri.add      3 4072.5 4091.3 -2033.2   4066.5                     
#cirri.polyP    4 4071.9 4097.0 -2031.9   4063.9 2.5912  1     0.1075

# CHAETAE LENGTH
# loci on chromosomes 9 and 3
chlen.dom2 <- lmer(MatGeno$pheno$ChLen ~ qtl1.chlen + qtl2.chlen + chlenqtl2.h + (1|dadID) + (1|momID), REML = F)

LpropM.chlen <- get.lecitho.proportion(MatGeno, c(1, 2, 4:8, 10))
LpropP.chlen <- get.lecitho.proportion(PatGeno, c(1, 2, 4:8, 10))
LpropZ.chlen <- (LpropM.chlen + LpropP.chlen)/2

chlen.polyZ <- lmer(MatGeno$pheno$ChLen ~ qtl1.chlen + qtl2.chlen + chlenqtl2.h + LpropZ.chlen + (1|dadID) + (1|momID), REML = F)
chlen.polyM <- lmer(MatGeno$pheno$ChLen ~ qtl1.chlen + qtl2.chlen + chlenqtl2.h + LpropM.chlen + (1|dadID) + (1|momID), REML = F)
chlen.polyP <- lmer(MatGeno$pheno$ChLen ~ qtl1.chlen + qtl2.chlen + chlenqtl2.h + LpropP.chlen + (1|dadID) + (1|momID), REML = F)

anova(chlen.polyZ, chlen.dom2)
#            npar   AIC   BIC logLik deviance  Chisq Df Pr(>Chisq)
#chlen.dom2     7 29086 29128 -14536    29072                     
#chlen.polyZ    8 29088 29136 -14536    29072 0.1698  1     0.6803

anova(chlen.polyM, chlen.dom2)
#            npar   AIC   BIC logLik deviance  Chisq Df Pr(>Chisq)
#chlen.dom2     7 29086 29128 -14536    29072                     
#chlen.polyM    8 29088 29136 -14536    29072 0.3663  1      0.545


anova(chlen.polyP, chlen.dom2)
#            npar   AIC   BIC logLik deviance  Chisq Df Pr(>Chisq)
#chlen.dom2     7 29086 29128 -14536    29072                     
#chlen.polyP    8 29086 29134 -14535    29070 2.0183  1     0.1554

# CHAETAE NUMBER - count
# loci on chromosomes 3 and 8
chnum.bc.add <- lmer(bc.chnum0 ~ qtl1.chnum.combined + qtl2.chnum + (1|dadID) + (1|momID), REML = F)

LpropM.chnum <- get.lecitho.proportion(MatGeno, c(1, 2, 4:7, 9, 10))
LpropP.chnum <- get.lecitho.proportion(PatGeno, c(1, 2, 4:7, 9, 10))
LpropZ.chnum <- (LpropM.chnum + LpropP.chnum)/2

chnum.bc.polyZ <- lmer(bc.chnum0 ~ qtl1.chnum.combined + qtl2.chnum + LpropZ.chnum + (1|dadID) + (1|momID), REML = F)
chnum.bc.polyM <- lmer(bc.chnum0 ~ qtl1.chnum.combined + qtl2.chnum + LpropM.chnum + (1|dadID) + (1|momID), REML = F)
chnum.bc.polyP <- lmer(bc.chnum0 ~ qtl1.chnum.combined + qtl2.chnum + LpropP.chnum + (1|dadID) + (1|momID), REML = F)

anova(chnum.bc.polyZ, chnum.bc.add)
#               npar    AIC    BIC  logLik deviance  Chisq Df Pr(>Chisq)
#chnum.bc.add      6 8376.1 8412.0 -4182.0   8364.1                     
#chnum.bc.polyZ    7 8377.6 8419.6 -4181.8   8363.6 0.4351  1     0.5095

anova(chnum.bc.polyM, chnum.bc.add)
#               npar    AIC    BIC  logLik deviance  Chisq Df Pr(>Chisq)
#chnum.bc.add      6 8376.1 8412.0 -4182.0   8364.1                     
#chnum.bc.polyM    7 8377.7 8419.7 -4181.9   8363.7 0.3565  1     0.5505

anova(chnum.bc.polyP, chnum.bc.add)
#               npar    AIC  BIC logLik deviance  Chisq Df Pr(>Chisq)
#chnum.bc.add      6 8376.1 8412  -4182   8364.1                     
#chnum.bc.polyP    7 8378.0 8420  -4182   8364.0 0.0415  1     0.8385

# CHAETAE NUMBER - presence
# loci on chromosomes 3 and 8; same as count trait above.
chnum.bin.dom12 <- glmer(chnum.bin.pheno ~ qtl1.chnum.combined + qtl2.chnum + chnumqtl1.h + chnumqtl2.h + (1|momID), family = "binomial")

chnum.bin.polyZ <- glmer(chnum.bin.pheno ~ qtl1.chnum.combined + qtl2.chnum + chnumqtl1.h + chnumqtl2.h + LpropZ.chnum + (1|momID), family = "binomial")
chnum.bin.polyM <- glmer(chnum.bin.pheno ~ qtl1.chnum.combined + qtl2.chnum + chnumqtl1.h + chnumqtl2.h + LpropM.chnum + (1|momID), family = "binomial")
chnum.bin.polyP <- glmer(chnum.bin.pheno ~ qtl1.chnum.combined + qtl2.chnum + chnumqtl1.h + chnumqtl2.h + LpropP.chnum + (1|momID), family = "binomial")

anova(chnum.bin.polyZ, chnum.bin.dom12)
#                npar    AIC    BIC  logLik deviance  Chisq Df Pr(>Chisq)  
#chnum.bin.dom12    6 3651.1 3688.8 -1819.5   3639.1                       
#chnum.bin.polyZ    7 3650.3 3694.3 -1818.2   3636.3 2.7308  1    0.09843 .


anova(chnum.bin.polyM, chnum.bin.dom12)
#                npar    AIC    BIC  logLik deviance  Chisq Df Pr(>Chisq)
#chnum.bin.dom12    6 3651.1 3688.8 -1819.5   3639.1                     
#chnum.bin.polyM    7 3651.4 3695.4 -1818.7   3637.4 1.6275  1      0.202

anova(chnum.bin.polyP, chnum.bin.dom12)
#                npar    AIC    BIC  logLik deviance  Chisq Df Pr(>Chisq)
#chnum.bin.dom12    6 3651.1 3688.8 -1819.5   3639.1                     
#chnum.bin.polyP    7 3652.5 3696.4 -1819.2   3638.5 0.6151  1     0.4329


## TEST FOR QUADRATIC OR CUBIC EFFECTS
# CIRRI 
# additive QTL on chromosome 5
cirri.add <- glmer(MatGeno$pheno$cirri ~  qtl1.cirri + (1|momID), family = "binomial")

LpropM.cirri <- get.lecitho.proportion(MatGeno, c(1:4,6:10))
LpropP.cirri <- get.lecitho.proportion(PatGeno, c(1:4,6:10))
LpropZ.cirri <- (LpropM.cirri + LpropP.cirri)/2

cirri.polyZ <- glmer(MatGeno$pheno$cirri ~  qtl1.cirri + LpropZ.cirri + (1|momID), family = "binomial")
cirri.polyM <- glmer(MatGeno$pheno$cirri ~  qtl1.cirri + LpropM.cirri + (1|momID), family = "binomial")
cirri.polyP <- glmer(MatGeno$pheno$cirri ~  qtl1.cirri + LpropP.cirri + (1|momID), family = "binomial")

cirri.polyZ2 <- glmer(MatGeno$pheno$cirri ~  qtl1.cirri + poly(LpropZ.cirri, 2, raw = T) + (1|momID), family = "binomial")
cirri.polyM2 <- glmer(MatGeno$pheno$cirri ~  qtl1.cirri + poly(LpropM.cirri, 2, raw = T) + (1|momID), family = "binomial")
cirri.polyP2 <- glmer(MatGeno$pheno$cirri ~  qtl1.cirri + poly(LpropP.cirri, 2, raw = T) + (1|momID), family = "binomial")

cirri.polyZ3 <- glmer(MatGeno$pheno$cirri ~  qtl1.cirri + poly(LpropZ.cirri, 3, raw = T) + (1|momID), family = "binomial")
cirri.polyM3 <- glmer(MatGeno$pheno$cirri ~  qtl1.cirri + poly(LpropM.cirri, 3, raw = T) + (1|momID), family = "binomial")
cirri.polyP3 <- glmer(MatGeno$pheno$cirri ~  qtl1.cirri + poly(LpropP.cirri, 3, raw = T) + (1|momID), family = "binomial")

# One nominally significant result: cubic fit for zygotic polygenic background. 
#cirri.add: MatGeno$pheno$cirri ~ qtl1.cirri + (1 | momID)
#cirri.polyZ3raw: MatGeno$pheno$cirri ~ qtl1.cirri + poly(LpropZ.cirri, 3, raw = T) + (1 | momID)
#                npar    AIC    BIC  logLik deviance  Chisq Df Pr(>Chisq)   
#cirri.add          3 4072.5 4091.3 -2033.2   4066.5                        
#cirri.polyZ3raw    6 4065.6 4103.3 -2026.8   4053.6 12.873  3   0.004921 **


# CHAETAE LENGTH
# loci on chromosomes 9 and 3
chlen.dom2 <- lmer(MatGeno$pheno$ChLen ~ qtl1.chlen + qtl2.chlen + chlenqtl2.h + (1|dadID) + (1|momID), REML = F)

LpropM.chlen <- get.lecitho.proportion(MatGeno, c(1, 2, 4:8, 10))
LpropP.chlen <- get.lecitho.proportion(PatGeno, c(1, 2, 4:8, 10))
LpropZ.chlen <- (LpropM.chlen + LpropP.chlen)/2

chlen.polyZ <- lmer(MatGeno$pheno$ChLen ~ qtl1.chlen + qtl2.chlen + chlenqtl2.h + LpropZ.chlen + (1|dadID) + (1|momID), REML = F)
chlen.polyM <- lmer(MatGeno$pheno$ChLen ~ qtl1.chlen + qtl2.chlen + chlenqtl2.h + LpropM.chlen + (1|dadID) + (1|momID), REML = F)
chlen.polyP <- lmer(MatGeno$pheno$ChLen ~ qtl1.chlen + qtl2.chlen + chlenqtl2.h + LpropP.chlen + (1|dadID) + (1|momID), REML = F)

chlen.polyZ2 <- lmer(MatGeno$pheno$ChLen ~ qtl1.chlen + qtl2.chlen + chlenqtl2.h + poly(LpropZ.chlen, 2, raw = T) + (1|dadID) + (1|momID), REML = F)
chlen.polyM2 <- lmer(MatGeno$pheno$ChLen ~ qtl1.chlen + qtl2.chlen + chlenqtl2.h + poly(LpropM.chlen, 2, raw = T) + (1|dadID) + (1|momID), REML = F)
chlen.polyP2 <- lmer(MatGeno$pheno$ChLen ~ qtl1.chlen + qtl2.chlen + chlenqtl2.h + poly(LpropP.chlen, 2, raw = T) + (1|dadID) + (1|momID), REML = F)

chlen.polyZ3 <- lmer(MatGeno$pheno$ChLen ~ qtl1.chlen + qtl2.chlen + chlenqtl2.h + poly(LpropZ.chlen, 3, raw = T) + (1|dadID) + (1|momID), REML = F)
chlen.polyM3 <- lmer(MatGeno$pheno$ChLen ~ qtl1.chlen + qtl2.chlen + chlenqtl2.h + poly(LpropM.chlen, 3, raw = T) + (1|dadID) + (1|momID), REML = F)
chlen.polyP3 <- lmer(MatGeno$pheno$ChLen ~ qtl1.chlen + qtl2.chlen + chlenqtl2.h + poly(LpropP.chlen, 3, raw = T) + (1|dadID) + (1|momID), REML = F)

# CHAETAE NUMBER - count
# loci on chromosomes 3 and 8
chnum.bc.add <- lmer(bc.chnum0 ~ qtl1.chnum.combined + qtl2.chnum + (1|dadID) + (1|momID), REML = F)

LpropM.chnum <- get.lecitho.proportion(MatGeno, c(1, 2, 4:7, 9, 10))
LpropP.chnum <- get.lecitho.proportion(PatGeno, c(1, 2, 4:7, 9, 10))
LpropZ.chnum <- (LpropM.chnum + LpropP.chnum)/2

chnum.bc.polyZ <- lmer(bc.chnum0 ~ qtl1.chnum.combined + qtl2.chnum + LpropZ.chnum + (1|dadID) + (1|momID), REML = F)
chnum.bc.polyM <- lmer(bc.chnum0 ~ qtl1.chnum.combined + qtl2.chnum + LpropM.chnum + (1|dadID) + (1|momID), REML = F)
chnum.bc.polyP <- lmer(bc.chnum0 ~ qtl1.chnum.combined + qtl2.chnum + LpropP.chnum + (1|dadID) + (1|momID), REML = F)

chnum.bc.polyZ2 <- lmer(bc.chnum0 ~ qtl1.chnum.combined + qtl2.chnum + poly(LpropZ.chnum, 2, raw = T) + (1|dadID) + (1|momID), REML = F)
chnum.bc.polyM2 <- lmer(bc.chnum0 ~ qtl1.chnum.combined + qtl2.chnum + poly(LpropM.chnum, 2, raw = T) + (1|dadID) + (1|momID), REML = F)
chnum.bc.polyP2 <- lmer(bc.chnum0 ~ qtl1.chnum.combined + qtl2.chnum + poly(LpropP.chnum, 2, raw = T) + (1|dadID) + (1|momID), REML = F)

chnum.bc.polyZ3 <- lmer(bc.chnum0 ~ qtl1.chnum.combined + qtl2.chnum + poly(LpropZ.chnum, 3, raw = T) + (1|dadID) + (1|momID), REML = F)
chnum.bc.polyM3 <- lmer(bc.chnum0 ~ qtl1.chnum.combined + qtl2.chnum + poly(LpropM.chnum, 3, raw = T) + (1|dadID) + (1|momID), REML = F)
chnum.bc.polyP3 <- lmer(bc.chnum0 ~ qtl1.chnum.combined + qtl2.chnum + poly(LpropP.chnum, 3, raw = T) + (1|dadID) + (1|momID), REML = F)

# CHAETAE NUMBER - presence
# loci on chromosomes 3 and 8; same as count trait above.
chnum.bin.dom12 <- glmer(chnum.bin.pheno ~ qtl1.chnum.combined + qtl2.chnum + chnumqtl1.h + chnumqtl2.h + (1|momID), family = "binomial")

chnum.bin.polyZ <- glmer(chnum.bin.pheno ~ qtl1.chnum.combined + qtl2.chnum + chnumqtl1.h + chnumqtl2.h + LpropZ.chnum + (1|momID), family = "binomial")
chnum.bin.polyM <- glmer(chnum.bin.pheno ~ qtl1.chnum.combined + qtl2.chnum + chnumqtl1.h + chnumqtl2.h + LpropM.chnum + (1|momID), family = "binomial")
chnum.bin.polyP <- glmer(chnum.bin.pheno ~ qtl1.chnum.combined + qtl2.chnum + chnumqtl1.h + chnumqtl2.h + LpropP.chnum + (1|momID), family = "binomial")

chnum.bin.polyZ2 <- glmer(chnum.bin.pheno ~ qtl1.chnum.combined + qtl2.chnum + chnumqtl1.h + chnumqtl2.h + poly(LpropZ.chnum, 2, raw = T) + (1|momID), family = "binomial")
chnum.bin.polyM2 <- glmer(chnum.bin.pheno ~ qtl1.chnum.combined + qtl2.chnum + chnumqtl1.h + chnumqtl2.h + poly(LpropM.chnum, 2, raw = T) + (1|momID), family = "binomial")
chnum.bin.polyP2 <- glmer(chnum.bin.pheno ~ qtl1.chnum.combined + qtl2.chnum + chnumqtl1.h + chnumqtl2.h + poly(LpropP.chnum, 2, raw = T) + (1|momID), family = "binomial")

chnum.bin.polyZ3 <- glmer(chnum.bin.pheno ~ qtl1.chnum.combined + qtl2.chnum + chnumqtl1.h + chnumqtl2.h + poly(LpropZ.chnum, 3, raw = T) + (1|momID), family = "binomial")
chnum.bin.polyM3 <- glmer(chnum.bin.pheno ~ qtl1.chnum.combined + qtl2.chnum + chnumqtl1.h + chnumqtl2.h + poly(LpropM.chnum, 3, raw = T) + (1|momID), family = "binomial")
chnum.bin.polyP3 <- glmer(chnum.bin.pheno ~ qtl1.chnum.combined + qtl2.chnum + chnumqtl1.h + chnumqtl2.h + poly(LpropP.chnum, 3, raw = T) + (1|momID), family = "binomial")
```
