## Supplementary File 1. Data and Code for "Baby makes three: maternal, paternal, and zygotic genetic effects shape larval phenotypic evolution": ChLenScan.R

```
# Chaetae Length trait scans for G3 phenotypes

library(qtl)
library(lme4)

load(file = "G3AnalysisObjects.R")

# Create an empty scanone object to hold the results
scan.holder <- scanone(MatGeno, pheno = "nChaetae")
# This will give a "calc.genoprob" warning; this can be ignored because we are just making an empty scanone object to fill in later.
scan.holder$lod <- 0
scan.holder[,4:5] <- 0


MZscan1 <- function(geno.mat, geno.pat, scan.object) {
	chlen.pheno <- geno.mat$pheno$ChLen
	dadID <- geno.mat$pheno$Dad_G2_Name
	momID <-  geno.mat$pheno$Mom_G2_Name
	null.LL <- logLik(lmer(chlen.pheno ~ 1 + (1|dadID) + (1|momID), REML = F))[1]

		for(j in 1: 10){	# This function is only for the autosomes
		chr <- names(nmar(geno.mat))[j]
		chlen.both <- NULL
		chlen.summed <- NULL

			for(i in 1:(nmar(geno.mat)[chr])){ 							
				qtlMom <- geno.mat$geno[[chr]]$data[,i] -1 # Count of lecithotroph alleles: 0, 1, or 2.
				qtlDad <- geno.pat$geno[[chr]]$data[,i] -1 # Count of lecithotroph alleles: 0, 1, or 2.
				qtlAll <- qtlMom + qtlDad				
		# Fit mixed models		
				chlen.both[i] <- logLik(lmer(chlen.pheno  ~ qtlMom + qtlDad + (1|dadID) + (1|momID), REML = F))[1]
				chlen.summed[i] <- logLik(lmer(chlen.pheno ~ qtlAll + (1|dadID) + (1|momID), REML = F))[1]
		}				
	scan.object[which(scan.object$chr == chr),3] <- chlen.both
	scan.object[which(scan.object$chr == chr),4] <- chlen.summed
	scan.object[which(scan.object$chr == chr),5] <- null.LL
	}

			for(j in 11){	# X only
		chr <- names(nmar(geno.mat))[j]
		chlen.both <- NULL
		chlen.summed <- NULL

			for(i in 1:(nmar(geno.mat)[chr])){ 	
						
#				print(paste(chr, i))	
						
				qtlMom <- geno.mat$geno[[chr]]$data[,i] -1 # Count of lecithotroph alleles: 0, or 1.
				qtlDad <- geno.pat$geno[[chr]]$data[,i] -1 # Count of lecithotroph alleles: 0, or 1.
				qtlAll <- (1/4)*qtlDad + (3/8)*qtlMom # X-specific G3 additive effect parameterization
		# Fit mixed models		
				chlen.both[i] <- logLik(lmer(chlen.pheno  ~ qtlMom + qtlDad + (1|dadID) + (1|momID), REML = F))[1]
				chlen.summed[i] <- logLik(lmer(chlen.pheno ~ qtlAll + (1|dadID) + (1|momID), REML = F))[1]
		}				
	scan.object[which(scan.object$chr == chr),3] <- chlen.both
	scan.object[which(scan.object$chr == chr),4] <- chlen.summed
	scan.object[which(scan.object$chr == chr),5] <- null.LL

	}
	names(scan.object)[3:5] <- c("Both","Summed","Null")
		
	return(scan.object)
}


date()
scan.out <- MZscan1(MatGeno, PatGeno, scan.holder)
date()

scan.out[,6:8] <- NA

scan.out[,6:8] <- cbind(2*(scan.out$Both - scan.out$Null), 2*(scan.out$Summed - scan.out$Null), 2*(scan.out$Both - scan.out$Summed))
names(scan.out)[6:8] <- c("BothChisq", "SummedChisq", "BetasChisq")

ChlenScan1.results <- scan.out
rm(scan.out)

#### End of first forward search.
apply(ChlenScan1.results[,c(3:4)], 2, function(x){ChlenScan1.results[which(x == max(x)),]})
# $Both
       # chr      pos      Both    Summed      Null BothChisq SummedChisq BetasChisq
# SNP272   9 1.457207 -14558.84 -14558.91 -14577.81  37.94488    37.80343  0.1414565
# SNP324   9 1.665381 -14558.84 -14558.91 -14577.81  37.94488    37.80343  0.1414565
# SNP836   9 1.873507 -14558.84 -14558.91 -14577.81  37.94488    37.80343  0.1414565
# SNP766   9 2.081634 -14558.84 -14558.91 -14577.81  37.94488    37.80343  0.1414565
# SNP328   9 2.289805 -14558.84 -14558.91 -14577.81  37.94488    37.80343  0.1414565

# $Summed
       # chr      pos      Both    Summed      Null BothChisq SummedChisq BetasChisq
# SNP272   9 1.457207 -14558.84 -14558.91 -14577.81  37.94488    37.80343  0.1414565
# SNP324   9 1.665381 -14558.84 -14558.91 -14577.81  37.94488    37.80343  0.1414565
# SNP836   9 1.873507 -14558.84 -14558.91 -14577.81  37.94488    37.80343  0.1414565
# SNP766   9 2.081634 -14558.84 -14558.91 -14577.81  37.94488    37.80343  0.1414565
# SNP328   9 2.289805 -14558.84 -14558.91 -14577.81  37.94488    37.80343  0.1414565


# permutations for autosomes and X

perm.combo <- c(MatGeno,PatGeno)
combo.dups <- findDupMarkers(perm.combo, adjacent.only = F)
MGnd <- drop.markers(MatGeno, unlist(combo.dups))
PGnd <- drop.markers(PatGeno, unlist(combo.dups))

# How many columns of results?
n.results <- 3
# How many permutations?
n.perms <- 200

#Prepare data for permutation by parent
cross.rows <- row.names(unique(data.frame(MatGeno$pheno$Male_ID, MatGeno$pheno$Female_ID)))
#Make mini cross objects that include each cross only once. 
mini.MGnd <- subset(MGnd, ind = as.numeric(cross.rows))
#make a mini.Gpat cross object:
mini.PGnd <- subset(MGnd, ind = as.numeric(cross.rows))

perm.scan.holder <- scanone(mini.MGnd, pheno = "nChaetae")
perm.scan.holder$lod <- 0
perm.scan.holder[,3:(2+n.results)] <- 0
perm.results <- matrix(nrow = n.perms, ncol = 2*n.results) # matrix(nrow = n.perms, ncol = n.results)

date()
for(m in 1:n.perms){
	
	male.perm <- data.frame(levels(MatGeno$pheno$Male_ID), levels(MatGeno$pheno$Male_ID))
	# Permutations are stratified by the F1 mother of the F2 fathers, i.e., 1A, 3C, 6F, 8H.
	male.perm[1:10,2] <- sample(male.perm[1:10,1])	
	male.perm[11:24,2] <- sample(male.perm[11:24,1])	
	male.perm[25:32,2] <- sample(male.perm[25:32,1])	
	male.perm[33:53,2] <- sample(male.perm[33:53,1])	

	female.perm <- data.frame(levels(MatGeno$pheno$Female_ID), levels(MatGeno$pheno$Female_ID)) 
	# Permutations are stratified by the F1 mother of the F2 mothers, i.e., 1A, 3C, 6F, 8H.
	female.perm[1:23,2] <- sample(female.perm[1:23,1])	
	female.perm[24:60,2] <- sample(female.perm[24:60,1])		
	female.perm[61:81,2] <- sample(female.perm[61:81,1])	
	female.perm[82:115,2] <- sample(female.perm[82:115,1])	
	
	perm.Gmat <- MGnd
	perm.Gpat <- PGnd
	
	for(i in 1:dim(MatGeno$pheno)[1]){
		permdad <- male.perm[which(male.perm[,1] == MatGeno$pheno$Male_ID[i]),2]
		permmom <- female.perm[which(female.perm[,1] == MatGeno$pheno$Female_ID[i]),2]
		for(j in 1:11){
			perm.Gpat$geno[[j]]$data[i,] <- mini.PGnd$geno[[j]]$data[which(mini.MGnd$pheno$Male_ID == permdad)[1],]
			perm.Gmat$geno[[j]]$data[i,] <- mini.MGnd$geno[[j]]$data[which(mini.MGnd$pheno$Female_ID == permmom)[1],]
		}
	}
	
permscan <- MZscan1(perm.Gmat, perm.Gpat, perm.scan.holder)

perm.results[m,1:n.results] <- apply(as.matrix(permscan[permscan$chr != "X",3:(2+n.results)]), 2, max)
perm.results[m, (n.results+1):(2*n.results)] <- apply(as.matrix(permscan[permscan$chr == "X",3:(2+n.results)]), 2, max)
perm.out <- data.frame(perm.results)
names(perm.out)  <- colnames(permscan)[3:(2+n.results)]
print(m); print(date());
}

rm(perm.results)
perm.out[,7:12] <- NA

perm.out[,7:12] <- cbind(2*(perm.out$Both - perm.out$Null), 2*(perm.out$Summed - perm.out$Null), 2*(perm.out$Both - perm.out$Summed), 2*(perm.out[,4] - perm.out$Null), 2*(perm.out[,5] - perm.out$Null), 2*(perm.out[,4] - perm.out[,5]))
names(perm.out)[7:12] <- c("BothChisq", "SummedChisq", "BetasChisq", "XBothChisq", "XSummedChisq", "XBetasChisq")

apply(perm.out[,7:12], 2, function(x){quantile(x,0.95)})
#   BothChisq  SummedChisq   BetasChisq   XBothChisq XSummedChisq  XBetasChisq 
#   14.230032    10.269908     7.210915     9.151639     7.321112     5.798716 


perm.out -> ChlenMZScan1Perms
save(ChlenMZScan1Perms, file = "ChlenScan1.200perms.R")
rm(perm.out)

# SNP272 is an additive-effect QTL

#################

chlen.qtl.9.272 <- MatGeno$geno[[9]]$data[,"SNP272"] + PatGeno$geno[[9]]$data[,"SNP272"]
last.column <- dim(MatGeno$pheno)[2]
MatGeno$pheno[,last.column+1] <- chlen.qtl.9.272
names(MatGeno$pheno)[last.column + 1] <- "chlen.qtl.9.272"

MZscan2 <- function(geno.mat, geno.pat, scan.object) {
	chlen.pheno <- geno.mat$pheno$ChLen
	dadID <- geno.mat$pheno$Dad_G2_Name
	momID <-  geno.mat$pheno$Mom_G2_Name
	null.LL <- logLik(lmer(chlen.pheno ~ geno.mat$pheno$chlen.qtl.9.272 + (1|dadID) + (1|momID), REML = F))[1]

		for(j in 1: 10){	# This function is only for the autosomes
		chr <- names(nmar(geno.mat))[j]
		chlen.both <- NULL
		chlen.summed <- NULL

			for(i in 1:(nmar(geno.mat)[chr])){ 							
				qtlMom <- geno.mat$geno[[chr]]$data[,i] -1 # Count of lecithotroph alleles: 0, 1, or 2.
				qtlDad <- geno.pat$geno[[chr]]$data[,i] -1 # Count of lecithotroph alleles: 0, 1, or 2.
				qtlAll <- qtlMom + qtlDad				
		# Fit mixed models		
				chlen.both[i] <- logLik(lmer(chlen.pheno  ~ qtlMom + qtlDad + geno.mat$pheno$chlen.qtl.9.272+ (1|dadID) + (1|momID), REML = F))[1]
				chlen.summed[i] <- logLik(lmer(chlen.pheno ~ qtlAll + geno.mat$pheno$chlen.qtl.9.272 + (1|dadID) + (1|momID), REML = F))[1]
		}				
	scan.object[which(scan.object$chr == chr),3] <- chlen.both
	scan.object[which(scan.object$chr == chr),4] <- chlen.summed
	scan.object[which(scan.object$chr == chr),5] <- null.LL
	}

			for(j in 11){	# X only
		chr <- names(nmar(geno.mat))[j]
		chlen.both <- NULL
		chlen.summed <- NULL

			for(i in 1:(nmar(geno.mat)[chr])){ 	
												
				qtlMom <- geno.mat$geno[[chr]]$data[,i] -1 # Count of lecithotroph alleles: 0, or 1.
				qtlDad <- geno.pat$geno[[chr]]$data[,i] -1 # Count of lecithotroph alleles: 0, or 1.
				qtlAll <- (1/4)*qtlDad + (3/8)*qtlMom # X-specific G3 additive effect parameterization
		# Fit mixed models		
				chlen.both[i] <- logLik(lmer(chlen.pheno  ~ qtlMom + qtlDad + geno.mat$pheno$chlen.qtl.9.272 + (1|dadID) + (1|momID), REML = F))[1]
				chlen.summed[i] <- logLik(lmer(chlen.pheno ~ qtlAll + geno.mat$pheno$chlen.qtl.9.272 + (1|dadID) + (1|momID), REML = F))[1]
		}				
	scan.object[which(scan.object$chr == chr),3] <- chlen.both
	scan.object[which(scan.object$chr == chr),4] <- chlen.summed
	scan.object[which(scan.object$chr == chr),5] <- null.LL

	}
	names(scan.object)[3:5] <- c("Both","Summed","Null")
		
	return(scan.object)
}


date()
scan.out <- MZscan2(MatGeno, PatGeno, scan.holder)
date()

scan.out[,6:8] <- NA

scan.out[,6:8] <- cbind(2*(scan.out$Both - scan.out$Null), 2*(scan.out$Summed - scan.out$Null), 2*(scan.out$Both - scan.out$Summed))
names(scan.out)[6:8] <- c("BothChisq", "SummedChisq", "BetasChisq")

ChlenScan2.results <- scan.out
rm(scan.out)

#### End of second forward search.

apply(ChlenScan2.results[,c(3:4)], 2, function(x){ChlenScan2.results[which(x == max(x)),]})
# $Both
       # chr      pos      Both   Summed      Null BothChisq SummedChisq BetasChisq
# SNP555   3 7.123157 -14539.63 -14539.7 -14558.91  38.56898    38.43529  0.1336933

# $Summed
       # chr      pos      Both   Summed      Null BothChisq SummedChisq BetasChisq
# SNP555   3 7.123157 -14539.63 -14539.7 -14558.91  38.56898    38.43529  0.1336933

# permutations for autosomes and X
perm.combo <- c(MatGeno,PatGeno)
combo.dups <- findDupMarkers(perm.combo, adjacent.only = F)
MGnd <- drop.markers(MatGeno, unlist(combo.dups))
PGnd <- drop.markers(PatGeno, unlist(combo.dups))

# How many columns of results?
n.results <- 3
# How many permutations?
n.perms <- 200

#Prepare data for permutation by parent
cross.rows <- row.names(unique(data.frame(MatGeno$pheno$Male_ID, MatGeno$pheno$Female_ID)))
#Make mini cross objects that include each cross only once. 
mini.MGnd <- subset(MGnd, ind = as.numeric(cross.rows))
#make a mini.Gpat cross object:
mini.PGnd <- subset(MGnd, ind = as.numeric(cross.rows))

perm.scan.holder <- scanone(mini.MGnd, pheno = "nChaetae")
perm.scan.holder$lod <- 0
perm.scan.holder[,3:(2+n.results)] <- 0
perm.results <- matrix(nrow = n.perms, ncol = 2*n.results) # matrix(nrow = n.perms, ncol = n.results)

date()
for(m in 1:n.perms){
	
	male.perm <- data.frame(levels(MatGeno$pheno$Male_ID), levels(MatGeno$pheno$Male_ID))
	# Permutations are stratified by the F1 mother of the F2 fathers, i.e., 1A, 3C, 6F, 8H.
	male.perm[1:10,2] <- sample(male.perm[1:10,1])	
	male.perm[11:24,2] <- sample(male.perm[11:24,1])	
	male.perm[25:32,2] <- sample(male.perm[25:32,1])	
	male.perm[33:53,2] <- sample(male.perm[33:53,1])	

	female.perm <- data.frame(levels(MatGeno$pheno$Female_ID), levels(MatGeno$pheno$Female_ID)) 
	# Permutations are stratified by the F1 mother of the F2 mothers, i.e., 1A, 3C, 6F, 8H.
	female.perm[1:23,2] <- sample(female.perm[1:23,1])	
	female.perm[24:60,2] <- sample(female.perm[24:60,1])		
	female.perm[61:81,2] <- sample(female.perm[61:81,1])	
	female.perm[82:115,2] <- sample(female.perm[82:115,1])	
	
	perm.Gmat <- MGnd
	perm.Gpat <- PGnd
	
	for(i in 1:dim(MatGeno$pheno)[1]){
		permdad <- male.perm[which(male.perm[,1] == MatGeno$pheno$Male_ID[i]),2]
		permmom <- female.perm[which(female.perm[,1] == MatGeno$pheno$Female_ID[i]),2]
		for(j in 1:11){
			perm.Gpat$geno[[j]]$data[i,] <- mini.PGnd$geno[[j]]$data[which(mini.MGnd$pheno$Male_ID == permdad)[1],]
			perm.Gmat$geno[[j]]$data[i,] <- mini.MGnd$geno[[j]]$data[which(mini.MGnd$pheno$Female_ID == permmom)[1],]
		}
	}
	
permscan <- MZscan2(perm.Gmat, perm.Gpat, perm.scan.holder)

perm.results[m,1:n.results] <- apply(as.matrix(permscan[permscan$chr != "X",3:(2+n.results)]), 2, max)
perm.results[m, (n.results+1):(2*n.results)] <- apply(as.matrix(permscan[permscan$chr == "X",3:(2+n.results)]), 2, max)
perm.out <- data.frame(perm.results)
names(perm.out)  <- colnames(permscan)[3:(2+n.results)]
print(m); print(date());
}

rm(perm.results)
perm.out[,7:12] <- NA

perm.out[,7:12] <- cbind(2*(perm.out$Both - perm.out$Null), 2*(perm.out$Summed - perm.out$Null), 2*(perm.out$Both - perm.out$Summed), 2*(perm.out[,4] - perm.out$Null), 2*(perm.out[,5] - perm.out$Null), 2*(perm.out[,4] - perm.out[,5]))
names(perm.out)[7:12] <- c("BothChisq", "SummedChisq", "BetasChisq", "XBothChisq", "XSummedChisq", "XBetasChisq")

apply(perm.out[,7:12], 2, function(x){quantile(x,0.95)})
#   BothChisq  SummedChisq   BetasChisq   XBothChisq XSummedChisq  XBetasChisq 
   14.164734    11.810617     6.334571     9.129530     6.449983     5.510697 

perm.out -> ChlenMZScan2Perms
save(ChlenMZScan2Perms, file = "ChlenScan2.200perms.R")
rm(perm.out)

# SNP555 is an additive-effect QTL

#################

chlen.qtl.3.555 <- MatGeno$geno[[3]]$data[,"SNP555"] + PatGeno$geno[[3]]$data[,"SNP555"]
last.column <- dim(MatGeno$pheno)[2]
MatGeno$pheno[,last.column+1] <- chlen.qtl.3.555
names(MatGeno$pheno)[last.column + 1] <- "chlen.qtl.3.555"

MZscan3 <- function(geno.mat, geno.pat, scan.object) {
	chlen.pheno <- geno.mat$pheno$ChLen
	dadID <- geno.mat$pheno$Dad_G2_Name
	momID <-  geno.mat$pheno$Mom_G2_Name
	null.LL <- logLik(lmer(chlen.pheno ~ geno.mat$pheno$chlen.qtl.9.272 + geno.mat$pheno$chlen.qtl.3.555 + (1|dadID) + (1|momID), REML = F))[1]

		for(j in 1: 10){	#  autosomes
		chr <- names(nmar(geno.mat))[j]
		chlen.both <- NULL
		chlen.summed <- NULL

			for(i in 1:(nmar(geno.mat)[chr])){ 							
				qtlMom <- geno.mat$geno[[chr]]$data[,i] -1 # Count of lecithotroph alleles: 0, 1, or 2.
				qtlDad <- geno.pat$geno[[chr]]$data[,i] -1 # Count of lecithotroph alleles: 0, 1, or 2.
				qtlAll <- qtlMom + qtlDad				
		# Fit mixed models		
				chlen.both[i] <- logLik(lmer(chlen.pheno  ~ qtlMom + qtlDad + geno.mat$pheno$chlen.qtl.9.272 + geno.mat$pheno$chlen.qtl.3.555 + (1|dadID) + (1|momID), REML = F))[1]
				chlen.summed[i] <- logLik(lmer(chlen.pheno ~ qtlAll + geno.mat$pheno$chlen.qtl.9.272 + geno.mat$pheno$chlen.qtl.3.555 + (1|dadID) + (1|momID), REML = F))[1]
		}				
	scan.object[which(scan.object$chr == chr),3] <- chlen.both
	scan.object[which(scan.object$chr == chr),4] <- chlen.summed
	scan.object[which(scan.object$chr == chr),5] <- null.LL
	}

			for(j in 11){	# X only
		chr <- names(nmar(geno.mat))[j]
		chlen.both <- NULL
		chlen.summed <- NULL

			for(i in 1:(nmar(geno.mat)[chr])){ 	
						
#				print(paste(chr, i))	
						
				qtlMom <- geno.mat$geno[[chr]]$data[,i] -1 # Count of lecithotroph alleles: 0, or 1.
				qtlDad <- geno.pat$geno[[chr]]$data[,i] -1 # Count of lecithotroph alleles: 0, or 1.
				qtlAll <- (1/4)*qtlDad + (3/8)*qtlMom # X-specific G3 additive effect parameterization
		# Fit mixed models		
				chlen.both[i] <- logLik(lmer(chlen.pheno  ~ qtlMom + qtlDad + geno.mat$pheno$chlen.qtl.9.272 + geno.mat$pheno$chlen.qtl.3.555 + (1|dadID) + (1|momID), REML = F))[1]
				chlen.summed[i] <- logLik(lmer(chlen.pheno ~ qtlAll + geno.mat$pheno$chlen.qtl.9.272 + geno.mat$pheno$chlen.qtl.3.555 + (1|dadID) + (1|momID), REML = F))[1]
		}				
	scan.object[which(scan.object$chr == chr),3] <- chlen.both
	scan.object[which(scan.object$chr == chr),4] <- chlen.summed
	scan.object[which(scan.object$chr == chr),5] <- null.LL

	}
	names(scan.object)[3:5] <- c("Both","Summed","Null")
		
	return(scan.object)
}

date()
scan.out <- MZscan3(MatGeno, PatGeno, scan.holder)
date()

scan.out[,6:8] <- NA

scan.out[,6:8] <- cbind(2*(scan.out$Both - scan.out$Null), 2*(scan.out$Summed - scan.out$Null), 2*(scan.out$Both - scan.out$Summed))
names(scan.out)[6:8] <- c("BothChisq", "SummedChisq", "BetasChisq")

ChlenScan3.results <- scan.out
rm(scan.out)

#### End of third forward search.

apply(ChlenScan3.results[,c(3:4)], 2, function(x){ChlenScan3.results[which(x == max(x)),]})
# $Both
       # chr      pos      Both    Summed     Null BothChisq SummedChisq BetasChisq
# SNP422   5 10.59194 -14535.53 -14536.97 -14539.7  8.325541    5.456097   2.869444

# $Summed
       # chr      pos      Both    Summed     Null BothChisq SummedChisq BetasChisq
# SNP372   8 2.912722 -14536.25 -14536.86 -14539.7  6.888705    5.679166   1.209539


# permutations for autosomes and X
perm.combo <- c(MatGeno,PatGeno)
combo.dups <- findDupMarkers(perm.combo, adjacent.only = F)
MGnd <- drop.markers(MatGeno, unlist(combo.dups))
PGnd <- drop.markers(PatGeno, unlist(combo.dups))

# How many columns of results?
n.results <- 3
# How many permutations?
n.perms <- 1000

#Prepare data for permutation by parent
cross.rows <- row.names(unique(data.frame(MatGeno$pheno$Male_ID, MatGeno$pheno$Female_ID)))
#Make mini cross objects that include each cross only once. 
mini.MGnd <- subset(MGnd, ind = as.numeric(cross.rows))
#make a mini.Gpat cross object:
mini.PGnd <- subset(MGnd, ind = as.numeric(cross.rows))

perm.scan.holder <- scanone(mini.MGnd, pheno = "nChaetae")
perm.scan.holder$lod <- 0
perm.scan.holder[,3:(2+n.results)] <- 0
perm.results <- matrix(nrow = n.perms, ncol = 2*n.results) # matrix(nrow = n.perms, ncol = n.results)

date()
for(m in 1:n.perms){
	
	male.perm <- data.frame(levels(MatGeno$pheno$Male_ID), levels(MatGeno$pheno$Male_ID))
	# Permutations are stratified by the F1 mother of the F2 fathers, i.e., 1A, 3C, 6F, 8H.
	male.perm[1:10,2] <- sample(male.perm[1:10,1])	
	male.perm[11:24,2] <- sample(male.perm[11:24,1])	
	male.perm[25:32,2] <- sample(male.perm[25:32,1])	
	male.perm[33:53,2] <- sample(male.perm[33:53,1])	

	female.perm <- data.frame(levels(MatGeno$pheno$Female_ID), levels(MatGeno$pheno$Female_ID)) 
	# Permutations are stratified by the F1 mother of the F2 mothers, i.e., 1A, 3C, 6F, 8H.
	female.perm[1:23,2] <- sample(female.perm[1:23,1])	
	female.perm[24:60,2] <- sample(female.perm[24:60,1])		
	female.perm[61:81,2] <- sample(female.perm[61:81,1])	
	female.perm[82:115,2] <- sample(female.perm[82:115,1])	
	
	perm.Gmat <- MGnd
	perm.Gpat <- PGnd
	
	for(i in 1:dim(MatGeno$pheno)[1]){
		permdad <- male.perm[which(male.perm[,1] == MatGeno$pheno$Male_ID[i]),2]
		permmom <- female.perm[which(female.perm[,1] == MatGeno$pheno$Female_ID[i]),2]
		for(j in 1:11){
			perm.Gpat$geno[[j]]$data[i,] <- mini.PGnd$geno[[j]]$data[which(mini.MGnd$pheno$Male_ID == permdad)[1],]
			perm.Gmat$geno[[j]]$data[i,] <- mini.MGnd$geno[[j]]$data[which(mini.MGnd$pheno$Female_ID == permmom)[1],]
		}
	}
	
permscan <- MZscan3(perm.Gmat, perm.Gpat, perm.scan.holder)

perm.results[m,1:n.results] <- apply(as.matrix(permscan[permscan$chr != "X",3:(2+n.results)]), 2, max)
perm.results[m, (n.results+1):(2*n.results)] <- apply(as.matrix(permscan[permscan$chr == "X",3:(2+n.results)]), 2, max)
perm.out <- data.frame(perm.results)
names(perm.out)  <- colnames(permscan)[3:(2+n.results)]
print(m); print(date());
}

rm(perm.results)
perm.out[,7:12] <- NA

perm.out[,7:12] <- cbind(2*(perm.out$Both - perm.out$Null), 2*(perm.out$Summed - perm.out$Null), 2*(perm.out$Both - perm.out$Summed), 2*(perm.out[,4] - perm.out$Null), 2*(perm.out[,5] - perm.out$Null), 2*(perm.out[,4] - perm.out[,5]))
names(perm.out)[7:12] <- c("BothChisq", "SummedChisq", "BetasChisq", "XBothChisq", "XSummedChisq", "XBetasChisq")

apply(perm.out[,7:12], 2, function(x){quantile(x,0.95)})
#   BothChisq  SummedChisq   BetasChisq   XBothChisq XSummedChisq  XBetasChisq 
#   13.904683    11.350412     6.962770    10.534858     7.494182     5.458950 

perm.out -> ChlenMZScan3Perms
save(ChlenMZScan3Perms, file = "ChlenScan3.1000perms.R")
rm(perm.out)

# No additional significant QTL in the third scan.
```
