## Supplementary File 1. Data and Code for "Baby makes three: maternal, paternal, and zygotic genetic effects shape larval phenotypic evolution": ChNumScan.R

```
# Chaetae Number trait scans for the G3 phenotypes

library(qtl)
library(lme4)

load(file = "G3AnalysisObjects.R")

# Create an empty scanone object to hold the results
scan.holder <- scanone(MatGeno, pheno = "nChaetae")
# This will give a "calc.genoprob" warning; this can be ignored because we are just making an empty scanone object to fill in later.
scan.holder$lod <- 0
scan.holder[,4:8] <- 0

# Redefine the phenotypes as a binary trait and a Box-Cox-adjusted continuous trait
bin.pheno <- MatGeno$pheno$nChaetae
bin.pheno[bin.pheno > 0] <- 1
bc.pheno <- ((MatGeno$pheno$nChaetae^0.29)-1)/0.29 # uses the maximum likelihood Box-Cox coefficient for these data
bc.pheno[MatGeno$pheno$nChaetae == 0] <- NA
last.column <- dim(MatGeno$pheno)[2]
MatGeno$pheno[,(last.column + 1)] <- bin.pheno
MatGeno$pheno[,(last.column + 2)] <- bc.pheno
names(MatGeno$pheno)[c((last.column +1) : (last.column +2))] <- c("chnum.bin.pheno","chnum.bc.pheno")


MZscan1 <- function(geno.mat, geno.pat, scan.object) {
	dadID <- geno.mat$pheno$Dad_G2_Name
	momID <-  geno.mat$pheno$Mom_G2_Name
	binpheno <- geno.mat$pheno$chnum.bin.pheno
	bcpheno <- geno.mat$pheno$chnum.bc.pheno
		
	bin.null <- logLik(glmer(binpheno ~ 1 + (1|dadID) + (1|momID), family = "binomial"))[1]
	bc.null <- 	logLik(lmer(bcpheno ~ 1 + (1|dadID) + (1|momID), REML = F))[1]
	
		for(j in 1: 10){	# This function is only for the autosomes
		chr <- names(nmar(geno.mat))[j]
		bin.both <- NULL
		bc.both <- NULL
		bin.summed <- NULL
		bc.summed <- NULL
			for(i in 1:(nmar(geno.mat)[chr])){ 			
	
				qtlMom <- geno.mat$geno[[chr]]$data[,i] -1 # Count of lecithotroph alleles: 0, 1, or 2.
				qtlDad <- geno.pat$geno[[chr]]$data[,i] -1 # Count of lecithotroph alleles: 0, 1, or 2.
				qtlAll <- qtlMom + qtlDad 				

				bin.both[i] <- logLik(glmer(binpheno  ~ qtlMom + qtlDad + (1|dadID) + (1|momID), family = "binomial"))[1]
				bc.both[i] <- logLik(lmer(bcpheno ~ qtlMom + qtlDad + (1|dadID) + (1|momID), REML = F))[1]


				bin.summed[i] <- logLik(glmer(binpheno ~ qtlAll + (1|dadID) + (1|momID), family = "binomial"))[1]
				bc.summed[i] <- logLik(lmer(bcpheno ~ qtlAll + (1|dadID) + (1|momID), REML = F))[1]


		}				
	scan.object[which(scan.object$chr == chr),3] <- bin.both		
	scan.object[which(scan.object$chr == chr),4] <- bin.summed
	scan.object[which(scan.object$chr == chr),5] <- bin.null
	scan.object[which(scan.object$chr == chr),6] <- bc.both
	scan.object[which(scan.object$chr == chr),7] <- bc.summed
	scan.object[which(scan.object$chr == chr),8] <- bc.null
	}
			for(j in 11){	# X only
		chr <- names(nmar(geno.mat))[j]
		bin.summed <- NULL
		bc.summed <- NULL

			for(i in 1:(nmar(geno.mat)[chr])){ 													
				qtlMom <- geno.mat$geno[[chr]]$data[,i] -1 # Count of lecithotroph alleles: 0, or 1.
				qtlDad <- geno.pat$geno[[chr]]$data[,i] -1 # Count of lecithotroph alleles: 0, or 1.
				qtlAll <- (1/4)*qtlDad + (3/8)*qtlMom # X-specific G3 additive effect parameterization

				bin.both[i] <- logLik(glmer(binpheno  ~ qtlMom + qtlDad + (1|dadID) + (1|momID), family = "binomial"))[1]
				bc.both[i] <- logLik(lmer(bcpheno ~ qtlMom + qtlDad + (1|dadID) + (1|momID), REML = F))[1]
				bin.summed[i] <- logLik(glmer(binpheno ~ qtlAll + (1|dadID) + (1|momID), family = "binomial"))[1]
				bc.summed[i] <- logLik(lmer(bcpheno ~ qtlAll + (1|dadID) + (1|momID), REML = F))[1]

		}				
	scan.object[which(scan.object$chr == chr),3] <- bin.both		
	scan.object[which(scan.object$chr == chr),4] <- bin.summed
	scan.object[which(scan.object$chr == chr),5] <- bin.null
	scan.object[which(scan.object$chr == chr),6] <- bc.both
	scan.object[which(scan.object$chr == chr),7] <- bc.summed
	scan.object[which(scan.object$chr == chr),8] <- bc.null
	}
	names(scan.object)[c(3:8)] <- c("binBoth","binSummed","binNull","bcBoth","bcSummed","bcNull")
		
	return(scan.object)
}

date()
scan.out <- MZscan1(MatGeno, PatGeno, scan.holder)
date()

scan.out[,9:11] <- NA
scan.out[,9:11] <- c(scan.out$binBoth+scan.out$bcBoth, scan.out$binSummed+scan.out$bcSummed, scan.out$binNull+scan.out$bcNull)
names(scan.out)[9:11] <- c("combBoth", "combSummed", "combNull")

scan.out[,12:20] <- NA

scan.out[,12:20] <- cbind(2*(scan.out$binBoth - scan.out$binNull), 2*(scan.out$binSummed - scan.out$binNull), 2*(scan.out$binBoth - scan.out$binSummed), 2*(scan.out$bcBoth - scan.out$bcNull), 2*(scan.out$bcSummed - scan.out$bcNull), 2*(scan.out$bcBoth - scan.out$bcSummed), 2*(scan.out$combBoth - scan.out$combNull), 2*(scan.out$combSummed - scan.out$combNull), 2*(scan.out$combBoth - scan.out$combSummed))

names(scan.out)[12:20] <- c("binBothChisq", "binSummedChisq", "binBetasChisq", "bcBothChisq", "bcSummedChisq", "bcBetasChisq", "combBothChisq", "combSummedChisq", "combBetasChisq")

ChnumScan1.results <- scan.out
rm(scan.out)

#### End of first forward search.
apply(ChnumScan1.results[,c(3:4,6:7,9:10)], 2, function(x){ChnumScan1.results[which(x == max(x)),]})
# $binBoth
       # chr      pos   binBoth binSummed   binNull    bcBoth  bcSummed    bcNull  combBoth combSummed  combNull binBothChisq binSummedChisq binBetasChisq
# SNP230   8 2.080073 -1847.804 -1847.938 -1865.096 -4195.353 -4195.467 -4207.522 -6043.157  -6043.405 -6072.618     34.58453       34.31673     0.2677996
       # bcBothChisq bcSummedChisq bcBetasChisq combBothChisq combSummedChisq combBetasChisq
# SNP230    24.33792      24.10895    0.2289752      58.92245        58.42568      0.4967748

# $binSummed
       # chr      pos   binBoth binSummed   binNull    bcBoth  bcSummed    bcNull  combBoth combSummed  combNull binBothChisq binSummedChisq binBetasChisq
# SNP230   8 2.080073 -1847.804 -1847.938 -1865.096 -4195.353 -4195.467 -4207.522 -6043.157  -6043.405 -6072.618     34.58453       34.31673     0.2677996
       # bcBothChisq bcSummedChisq bcBetasChisq combBothChisq combSummedChisq combBetasChisq
# SNP230    24.33792      24.10895    0.2289752      58.92245        58.42568      0.4967748

# $bcBoth
        # chr      pos   binBoth binSummed   binNull    bcBoth  bcSummed    bcNull  combBoth combSummed  combNull binBothChisq binSummedChisq binBetasChisq
# SNP1015   8 4.161655 -1850.716  -1850.85 -1865.096 -4194.303 -4194.554 -4207.522 -6045.019  -6045.405 -6072.618     28.76101       28.49197     0.2690429
        # bcBothChisq bcSummedChisq bcBetasChisq combBothChisq combSummedChisq combBetasChisq
# SNP1015    26.43752      25.93512    0.5023951      55.19853        54.42709       0.771438

# $bcSummed
       # chr pos   binBoth binSummed   binNull    bcBoth  bcSummed    bcNull  combBoth combSummed  combNull binBothChisq binSummedChisq binBetasChisq bcBothChisq
# SNP768   8   0 -1849.501 -1849.551 -1865.096 -4194.383 -4194.455 -4207.522 -6043.884  -6044.006 -6072.618      31.1899        31.0898     0.1000954    26.27817
       # bcSummedChisq bcBetasChisq combBothChisq combSummedChisq combBetasChisq
# SNP768      26.13478    0.1433933      57.46807        57.22458      0.2434887

# $combBoth
       # chr      pos  binBoth binSummed   binNull    bcBoth  bcSummed    bcNull  combBoth combSummed  combNull binBothChisq binSummedChisq binBetasChisq
# SNP725   8 2.496386 -1848.32 -1848.443 -1865.096 -4194.754 -4194.874 -4207.522 -6043.074  -6043.317 -6072.618     33.55215       33.30675     0.2453993
       # bcBothChisq bcSummedChisq bcBetasChisq combBothChisq combSummedChisq combBetasChisq
# SNP725    25.53643      25.29564    0.2407901      59.08858        58.60239      0.4861894

# $combSummed
       # chr      pos  binBoth binSummed   binNull    bcBoth  bcSummed    bcNull  combBoth combSummed  combNull binBothChisq binSummedChisq binBetasChisq
# SNP725   8 2.496386 -1848.32 -1848.443 -1865.096 -4194.754 -4194.874 -4207.522 -6043.074  -6043.317 -6072.618     33.55215       33.30675     0.2453993
       # bcBothChisq bcSummedChisq bcBetasChisq combBothChisq combSummedChisq combBetasChisq
# SNP725    25.53643      25.29564    0.2407901      59.08858        58.60239      0.4861894


# permutations for autosomes and X

perm.combo <- c(MatGeno,PatGeno)
combo.dups <- findDupMarkers(perm.combo, adjacent.only = F)
MGnd <- drop.markers(MatGeno, unlist(combo.dups))
PGnd <- drop.markers(PatGeno, unlist(combo.dups))

# How many columns of results?
n.results <- 6
# How many permutations?
n.perms <- 100

#Prepare data for permutation by parent
cross.rows <- row.names(unique(data.frame(MatGeno$pheno$Male_ID, MatGeno$pheno$Female_ID)))
#Make mini cross objects that include each cross only once. 
mini.MGnd <- subset(MGnd, ind = as.numeric(cross.rows))
#make a mini.Gpat cross object:
mini.PGnd <- subset(MGnd, ind = as.numeric(cross.rows))

perm.scan.holder <- scanone(mini.MGnd, pheno = "nChaetae")
perm.scan.holder$lod <- 0
perm.scan.holder[,3:(2+n.results)] <- 0
perm.results <- matrix(nrow = n.perms, ncol = 2*9) 

date()
for(m in 1:n.perms){
	
	male.perm <- data.frame(levels(MatGeno$pheno$Male_ID), levels(MatGeno$pheno$Male_ID))
	# Permutations are stratified by the F1 mother of the F2 fathers, i.e., 1A, 3C, 6F, 8H.
	male.perm[1:10,2] <- sample(male.perm[1:10,1])	
	male.perm[11:24,2] <- sample(male.perm[11:24,1])	
	male.perm[25:32,2] <- sample(male.perm[25:32,1])	
	male.perm[33:53,2] <- sample(male.perm[33:53,1])	

	female.perm <- data.frame(levels(MatGeno$pheno$Female_ID), levels(MatGeno$pheno$Female_ID)) 
	# Permutations are stratified by the F1 mother of the F2 mothers, i.e., 1A, 3C, 6F, 8H.
	female.perm[1:23,2] <- sample(female.perm[1:23,1])	
	female.perm[24:60,2] <- sample(female.perm[24:60,1])		
	female.perm[61:81,2] <- sample(female.perm[61:81,1])	
	female.perm[82:115,2] <- sample(female.perm[82:115,1])	
	
	perm.Gmat <- MGnd
	perm.Gpat <- PGnd
	
	for(i in 1:dim(MatGeno$pheno)[1]){
		permdad <- male.perm[which(male.perm[,1] == MatGeno$pheno$Male_ID[i]),2]
		permmom <- female.perm[which(female.perm[,1] == MatGeno$pheno$Female_ID[i]),2]
		for(j in 1:11){
			perm.Gpat$geno[[j]]$data[i,] <- mini.PGnd$geno[[j]]$data[which(mini.MGnd$pheno$Male_ID == permdad)[1],]
			perm.Gmat$geno[[j]]$data[i,] <- mini.MGnd$geno[[j]]$data[which(mini.MGnd$pheno$Female_ID == permmom)[1],]
		}
	}
	
permscan <- MZscan1(perm.Gmat, perm.Gpat, perm.scan.holder)
permscan[,9:11] <- NA
permscan[,9:11] <- c(permscan$binBoth+ permscan$bcBoth, permscan$binSummed+ permscan$bcSummed, permscan$binNull+ permscan$bcNull)
names(permscan)[9:11] <- c("combBoth", "combSummed", "combNull")

perm.results[m,1:9] <- apply(as.matrix(permscan[permscan$chr != "X",3:(2+9)]), 2, max)
perm.results[m, (9+1):(2*9)] <- apply(as.matrix(permscan[permscan$chr == "X",3:(2+9)]), 2, max)
perm.out <- data.frame(perm.results)
names(perm.out)  <- colnames(permscan)[3:(2+9)]
print(m); print(date());
}


rm(perm.results)
perm.out[,19:30] <- NA

perm.out[,19:30] <- cbind( 2*(perm.out[,1] - perm.out[,3]), 2*(perm.out[,2] - perm.out[,3]), 2*(perm.out[,4] - perm.out[,6]), 2*(perm.out[,5] - perm.out[,6]), 2*(perm.out[,7] - perm.out[,9]), 2*(perm.out[,8] - perm.out[,9]), 2*(perm.out[,10] - perm.out[,12]), 2*(perm.out[,11] - perm.out[,12]), 2*(perm.out[,13] - perm.out[,15]), 2*(perm.out[,14] - perm.out[,15]), 2*(perm.out[,16] - perm.out[,18]), 2*(perm.out[,17] - perm.out[,18]))

names(perm.out)[19:30] <- c("binBothChisq", "binSummedChisq", "bcBothChisq", "bcSummedChisq", "combBothChisq", "combSummedChisq", paste0("X",c("binBothChisq", "binSummedChisq", "bcBothChisq", "bcSummedChisq", "combBothChisq", "combSummedChisq")))


apply(perm.out[,19:30], 2, function(x){quantile(x,0.95)})
    binBothChisq   binSummedChisq      bcBothChisq    bcSummedChisq    combBothChisq  combSummedChisq    XbinBothChisq  XbinSummedChisq     XbcBothChisq 
       13.376625        10.518907        15.810075        10.580495        26.169313        17.283097        11.647169         8.632339         9.450947 
  XbcSummedChisq   XcombBothChisq XcombSummedChisq 
        5.331924        18.574212        12.455031 


perm.out -> ChnumMZScan1Perms
save(ChnumMZScan1Perms, file = "ChnumScan1.100perms.R")
rm(perm.out)

# Chromosome 8, SNP725 is an additive zygotic locius for chaetae number

#################

qtl1.comb <- MatGeno$geno[[8]]$data[,"SNP725"] + PatGeno$geno[[8]]$data[,"SNP725"] - 2
last.column <- dim(MatGeno$pheno)[2]
MatGeno$pheno[,last.column+1] <- qtl1.comb
names(MatGeno$pheno)[last.column + 1] <- "qtl1comb"

# After including QTL covariates, the logistic model has convergence problems or singular fits for the random effects, because the estimated paternal random effect is zero. For easier fitting, we remove the paternal random effect from these models in the analyses below. Note that the likelihoods are identical with the effects included, but the models take longer to run, which is inconvenient for the permutations. 

MZscan2 <- function(geno.mat, geno.pat, scan.object) {
	dadID <- geno.mat$pheno$Dad_G2_Name
	momID <-  geno.mat$pheno$Mom_G2_Name
	binpheno <- geno.mat$pheno$chnum.bin.pheno
	bcpheno <- geno.mat$pheno$chnum.bc.pheno
		
	bin.null <- logLik(glmer(binpheno ~ geno.mat$pheno$qtl1comb + (1|momID), family = "binomial"))[1]
	bc.null <- 	logLik(lmer(bcpheno ~ geno.mat$pheno$qtl1comb + (1|dadID) + (1|momID), REML = F))[1]
	
		for(j in 1: 10){	# This function is only for the autosomes
		chr <- names(nmar(geno.mat))[j]
		bin.both <- NULL
		bc.both <- NULL
		bin.summed <- NULL
		bc.summed <- NULL
			for(i in 1:(nmar(geno.mat)[chr])){ 			
	
				qtlMom <- geno.mat$geno[[chr]]$data[,i] -1 # Count of lecithotroph alleles: 0, 1, or 2.
				qtlDad <- geno.pat$geno[[chr]]$data[,i] -1 # Count of lecithotroph alleles: 0, 1, or 2.
				qtlAll <- qtlMom + qtlDad 				

				bin.both[i] <- logLik(glmer(binpheno  ~ qtlMom + qtlDad + geno.mat$pheno$qtl1comb + (1|momID), family = "binomial"))[1]
				bc.both[i] <- logLik(lmer(bcpheno ~ qtlMom + qtlDad + geno.mat$pheno$qtl1comb + (1|dadID) + (1|momID), REML = F))[1]

				bin.summed[i] <- logLik(glmer(binpheno ~ qtlAll + geno.mat$pheno$qtl1comb + (1|momID), family = "binomial"))[1]
				bc.summed[i] <- logLik(lmer(bcpheno ~ qtlAll + geno.mat$pheno$qtl1comb + (1|dadID) + (1|momID), REML = F))[1]

		}				
	scan.object[which(scan.object$chr == chr),3] <- bin.both		
	scan.object[which(scan.object$chr == chr),4] <- bin.summed
	scan.object[which(scan.object$chr == chr),5] <- bin.null
	scan.object[which(scan.object$chr == chr),6] <- bc.both
	scan.object[which(scan.object$chr == chr),7] <- bc.summed
	scan.object[which(scan.object$chr == chr),8] <- bc.null
	}
			for(j in 11){	# X only
		chr <- names(nmar(geno.mat))[j]
		bin.summed <- NULL
		bc.summed <- NULL

			for(i in 1:(nmar(geno.mat)[chr])){ 													
				qtlMom <- geno.mat$geno[[chr]]$data[,i] -1 # Count of lecithotroph alleles: 0, or 1.
				qtlDad <- geno.pat$geno[[chr]]$data[,i] -1 # Count of lecithotroph alleles: 0, or 1.
				qtlAll <- (1/4)*qtlDad + (3/8)*qtlMom # X-specific G3 additive effect parameterization

				bin.both[i] <- logLik(glmer(binpheno  ~ qtlMom + qtlDad + geno.mat$pheno$qtl1comb + (1|momID), family = "binomial"))[1]
				bc.both[i] <- logLik(lmer(bcpheno ~ qtlMom + qtlDad + geno.mat$pheno$qtl1comb + (1|dadID) + (1|momID), REML = F))[1]

				bin.summed[i] <- logLik(glmer(binpheno ~ qtlAll + geno.mat$pheno$qtl1comb + (1|momID), family = "binomial"))[1]
				bc.summed[i] <- logLik(lmer(bcpheno ~ qtlAll + geno.mat$pheno$qtl1comb + (1|dadID) + (1|momID), REML = F))[1]

		}				
	scan.object[which(scan.object$chr == chr),3] <- bin.both		
	scan.object[which(scan.object$chr == chr),4] <- bin.summed
	scan.object[which(scan.object$chr == chr),5] <- bin.null
	scan.object[which(scan.object$chr == chr),6] <- bc.both
	scan.object[which(scan.object$chr == chr),7] <- bc.summed
	scan.object[which(scan.object$chr == chr),8] <- bc.null
	}
	names(scan.object)[c(3:8)] <- c("binBoth","binSummed","binNull","bcBoth","bcSummed","bcNull")
		
	return(scan.object)
}

date()
scan.out <- MZscan2(MatGeno, PatGeno, scan.holder)
date()

scan.out[,9:11] <- NA
scan.out[,9:11] <- c(scan.out$binBoth+scan.out$bcBoth, scan.out$binSummed+scan.out$bcSummed, scan.out$binNull+scan.out$bcNull)
names(scan.out)[9:11] <- c("combBoth", "combSummed", "combNull")

scan.out[,12:20] <- NA
scan.out[,12:20] <- cbind(2*(scan.out$binBoth - scan.out$binNull), 2*(scan.out$binSummed - scan.out$binNull), 2*(scan.out$binBoth - scan.out$binSummed), 2*(scan.out$bcBoth - scan.out$bcNull), 2*(scan.out$bcSummed - scan.out$bcNull), 2*(scan.out$bcBoth - scan.out$bcSummed), 2*(scan.out$combBoth - scan.out$combNull), 2*(scan.out$combSummed - scan.out$combNull), 2*(scan.out$combBoth - scan.out$combSummed))

names(scan.out)[12:20] <- c("binBothChisq", "binSummedChisq", "binBetasChisq", "bcBothChisq", "bcSummedChisq", "bcBetasChisq", "combBothChisq", "combSummedChisq", "combBetasChisq")

ChnumScan2.results <- scan.out
rm(scan.out)

#### End of second forward search.

apply(ChnumScan2.results[,c(3:4,6:7,9:10)], 2, function(x){ChnumScan2.results[which(x == max(x)),]})
# $binBoth
       # chr      pos   binBoth binSummed   binNull   bcBoth  bcSummed    bcNull  combBoth combSummed  combNull binBothChisq binSummedChisq binBetasChisq
# SNP219   3 4.816849 -1829.519 -1830.969 -1848.443 -4183.08 -4183.333 -4194.874 -6012.599  -6014.301 -6043.317     37.84784       34.94867      2.899164
       # bcBothChisq bcSummedChisq bcBetasChisq combBothChisq combSummedChisq combBetasChisq
# SNP219    23.58794      23.08277    0.5051744      61.43578        58.03144       3.404338

# $binSummed
       # chr     pos   binBoth binSummed   binNull    bcBoth  bcSummed    bcNull  combBoth combSummed  combNull binBothChisq binSummedChisq binBetasChisq
# SNP779   3 5.23311 -1829.602 -1830.935 -1848.443 -4181.599 -4182.028 -4194.874 -6011.202  -6012.963 -6043.317     37.68168       35.01605      2.665621
       # bcBothChisq bcSummedChisq bcBetasChisq combBothChisq combSummedChisq combBetasChisq
# SNP779    26.54931       25.6914    0.8579188      64.23099        60.70745        3.52354

# $bcBoth
       # chr     pos   binBoth binSummed   binNull    bcBoth  bcSummed    bcNull  combBoth combSummed  combNull binBothChisq binSummedChisq binBetasChisq
# SNP779   3 5.23311 -1829.602 -1830.935 -1848.443 -4181.599 -4182.028 -4194.874 -6011.202  -6012.963 -6043.317     37.68168       35.01605      2.665621
       # bcBothChisq bcSummedChisq bcBetasChisq combBothChisq combSummedChisq combBetasChisq
# SNP779    26.54931       25.6914    0.8579188      64.23099        60.70745        3.52354

# $bcSummed
       # chr     pos   binBoth binSummed   binNull    bcBoth  bcSummed    bcNull  combBoth combSummed  combNull binBothChisq binSummedChisq binBetasChisq
# SNP779   3 5.23311 -1829.602 -1830.935 -1848.443 -4181.599 -4182.028 -4194.874 -6011.202  -6012.963 -6043.317     37.68168       35.01605      2.665621
       # bcBothChisq bcSummedChisq bcBetasChisq combBothChisq combSummedChisq combBetasChisq
# SNP779    26.54931       25.6914    0.8579188      64.23099        60.70745        3.52354

# $combBoth
       # chr     pos   binBoth binSummed   binNull    bcBoth  bcSummed    bcNull  combBoth combSummed  combNull binBothChisq binSummedChisq binBetasChisq
# SNP779   3 5.23311 -1829.602 -1830.935 -1848.443 -4181.599 -4182.028 -4194.874 -6011.202  -6012.963 -6043.317     37.68168       35.01605      2.665621
       # bcBothChisq bcSummedChisq bcBetasChisq combBothChisq combSummedChisq combBetasChisq
# SNP779    26.54931       25.6914    0.8579188      64.23099        60.70745        3.52354

# $combSummed
       # chr     pos   binBoth binSummed   binNull    bcBoth  bcSummed    bcNull  combBoth combSummed  combNull binBothChisq binSummedChisq binBetasChisq
# SNP779   3 5.23311 -1829.602 -1830.935 -1848.443 -4181.599 -4182.028 -4194.874 -6011.202  -6012.963 -6043.317     37.68168       35.01605      2.665621
       # bcBothChisq bcSummedChisq bcBetasChisq combBothChisq combSummedChisq combBetasChisq
# SNP779    26.54931       25.6914    0.8579188      64.23099        60.70745        3.52354


# permutations for autosomes and X

perm.combo <- c(MatGeno,PatGeno)
combo.dups <- findDupMarkers(perm.combo, adjacent.only = F)
MGnd <- drop.markers(MatGeno, unlist(combo.dups))
PGnd <- drop.markers(PatGeno, unlist(combo.dups))

# How many columns of results?
n.results <- 6
# How many permutations?
n.perms <- 100

#Prepare data for permutation by parent
cross.rows <- row.names(unique(data.frame(MatGeno$pheno$Male_ID, MatGeno$pheno$Female_ID)))
#Make mini cross objects that include each cross only once. 
mini.MGnd <- subset(MGnd, ind = as.numeric(cross.rows))
#make a mini.Gpat cross object:
mini.PGnd <- subset(MGnd, ind = as.numeric(cross.rows))

perm.scan.holder <- scanone(mini.MGnd, pheno = "nChaetae")
perm.scan.holder$lod <- 0
perm.scan.holder[,3:(2+n.results)] <- 0
perm.results <- matrix(nrow = n.perms, ncol = 2*9) 

date()
for(m in 1:n.perms){
	
	male.perm <- data.frame(levels(MatGeno$pheno$Male_ID), levels(MatGeno$pheno$Male_ID))
	# Permutations are stratified by the F1 mother of the F2 fathers, i.e., 1A, 3C, 6F, 8H.
	male.perm[1:10,2] <- sample(male.perm[1:10,1])	
	male.perm[11:24,2] <- sample(male.perm[11:24,1])	
	male.perm[25:32,2] <- sample(male.perm[25:32,1])	
	male.perm[33:53,2] <- sample(male.perm[33:53,1])	

	female.perm <- data.frame(levels(MatGeno$pheno$Female_ID), levels(MatGeno$pheno$Female_ID)) 
	# Permutations are stratified by the F1 mother of the F2 mothers, i.e., 1A, 3C, 6F, 8H.
	female.perm[1:23,2] <- sample(female.perm[1:23,1])	
	female.perm[24:60,2] <- sample(female.perm[24:60,1])		
	female.perm[61:81,2] <- sample(female.perm[61:81,1])	
	female.perm[82:115,2] <- sample(female.perm[82:115,1])	
	
	perm.Gmat <- MGnd
	perm.Gpat <- PGnd
	
	for(i in 1:dim(MatGeno$pheno)[1]){
		permdad <- male.perm[which(male.perm[,1] == MatGeno$pheno$Male_ID[i]),2]
		permmom <- female.perm[which(female.perm[,1] == MatGeno$pheno$Female_ID[i]),2]
		for(j in 1:11){
			perm.Gpat$geno[[j]]$data[i,] <- mini.PGnd$geno[[j]]$data[which(mini.MGnd$pheno$Male_ID == permdad)[1],]
			perm.Gmat$geno[[j]]$data[i,] <- mini.MGnd$geno[[j]]$data[which(mini.MGnd$pheno$Female_ID == permmom)[1],]
		}
	}
	
permscan <- MZscan2(perm.Gmat, perm.Gpat, perm.scan.holder)
permscan[,9:11] <- NA
permscan[,9:11] <- c(permscan$binBoth+ permscan$bcBoth, permscan$binSummed+ permscan$bcSummed, permscan$binNull+ permscan$bcNull)
names(permscan)[9:11] <- c("combBoth", "combSummed", "combNull")

perm.results[m,1:9] <- apply(as.matrix(permscan[permscan$chr != "X",3:(2+9)]), 2, max)
perm.results[m, (9+1):(2*9)] <- apply(as.matrix(permscan[permscan$chr == "X",3:(2+9)]), 2, max)
perm.out <- data.frame(perm.results)
names(perm.out)  <- colnames(permscan)[3:(2+9)]
print(m); print(date());
}

rm(perm.results)
perm.out[,19:30] <- NA

perm.out[,19:30] <- cbind( 2*(perm.out[,1] - perm.out[,3]), 2*(perm.out[,2] - perm.out[,3]), 2*(perm.out[,4] - perm.out[,6]), 2*(perm.out[,5] - perm.out[,6]), 2*(perm.out[,7] - perm.out[,9]), 2*(perm.out[,8] - perm.out[,9]), 2*(perm.out[,10] - perm.out[,12]), 2*(perm.out[,11] - perm.out[,12]), 2*(perm.out[,13] - perm.out[,15]), 2*(perm.out[,14] - perm.out[,15]), 2*(perm.out[,16] - perm.out[,18]), 2*(perm.out[,17] - perm.out[,18]))

names(perm.out)[19:30] <- c("binBothChisq", "binSummedChisq", "bcBothChisq", "bcSummedChisq", "combBothChisq", "combSummedChisq", paste0("X",c("binBothChisq", "binSummedChisq", "bcBothChisq", "bcSummedChisq", "combBothChisq", "combSummedChisq")))


apply(perm.out[,19:30], 2, function(x){quantile(x,0.95)})
    # binBothChisq   binSummedChisq      bcBothChisq    bcSummedChisq    combBothChisq  combSummedChisq    XbinBothChisq  XbinSummedChisq     XbcBothChisq 
       # 14.265282        10.920086        14.030322        11.440395        22.508464        18.164444         9.972714         6.531498        10.757514 
  # XbcSummedChisq   XcombBothChisq XcombSummedChisq 
        # 7.752003        15.463217        10.434717 

perm.out -> ChnumMZScan2Perms
save(ChnumMZScan2Perms, file = "ChnumScan2.100perms.R")
rm(perm.out)

# SNP779 is an additive-effect QTL

#################

qtl2.comb <- MatGeno$geno[[3]]$data[,"SNP779"] + PatGeno$geno[[3]]$data[,"SNP779"] - 2
last.column <- dim(MatGeno$pheno)[2]
MatGeno$pheno[,last.column+1] <- qtl2.comb
names(MatGeno$pheno)[last.column + 1] <- "qtl2comb"

MZscan3 <- function(geno.mat, geno.pat, scan.object) {
	dadID <- geno.mat$pheno$Dad_G2_Name
	momID <-  geno.mat$pheno$Mom_G2_Name
	binpheno <- geno.mat$pheno$chnum.bin.pheno
	bcpheno <- geno.mat$pheno$chnum.bc.pheno
		
	bin.null <- logLik(glmer(binpheno ~ geno.mat$pheno$qtl1comb + geno.mat$pheno$qtl2comb + (1|momID), family = "binomial"))[1]
	bc.null <- 	logLik(lmer(bcpheno ~ geno.mat$pheno$qtl1comb + geno.mat$pheno$qtl2comb + (1|dadID) + (1|momID), REML = F))[1]
	
		for(j in 1: 10){	# This function is only for the autosomes
		chr <- names(nmar(geno.mat))[j]
		bin.both <- NULL
		bc.both <- NULL
		bin.summed <- NULL
		bc.summed <- NULL
			for(i in 1:(nmar(geno.mat)[chr])){ 			
	
				qtlMom <- geno.mat$geno[[chr]]$data[,i] -1 # Count of lecithotroph alleles: 0, 1, or 2.
				qtlDad <- geno.pat$geno[[chr]]$data[,i] -1 # Count of lecithotroph alleles: 0, 1, or 2.
				qtlAll <- qtlMom + qtlDad 				

				bin.both[i] <- logLik(glmer(binpheno  ~ qtlMom + qtlDad + geno.mat$pheno$qtl1comb + geno.mat$pheno$qtl2comb + (1|momID), family = "binomial"))[1]
				bc.both[i] <- logLik(lmer(bcpheno ~ qtlMom + qtlDad + geno.mat$pheno$qtl1comb + geno.mat$pheno$qtl2comb + (1|dadID) + (1|momID), REML = F))[1]

				bin.summed[i] <- logLik(glmer(binpheno ~ qtlAll + geno.mat$pheno$qtl1comb + geno.mat$pheno$qtl2comb + (1|momID), family = "binomial"))[1]
				bc.summed[i] <- logLik(lmer(bcpheno ~ qtlAll + geno.mat$pheno$qtl1comb + geno.mat$pheno$qtl2comb + (1|dadID) + (1|momID), REML = F))[1]

		}				
	scan.object[which(scan.object$chr == chr),3] <- bin.both		
	scan.object[which(scan.object$chr == chr),4] <- bin.summed
	scan.object[which(scan.object$chr == chr),5] <- bin.null
	scan.object[which(scan.object$chr == chr),6] <- bc.both
	scan.object[which(scan.object$chr == chr),7] <- bc.summed
	scan.object[which(scan.object$chr == chr),8] <- bc.null
	}
			for(j in 11){	# X only
		chr <- names(nmar(geno.mat))[j]
		bin.summed <- NULL
		bc.summed <- NULL

			for(i in 1:(nmar(geno.mat)[chr])){ 													
				qtlMom <- geno.mat$geno[[chr]]$data[,i] -1 # Count of lecithotroph alleles: 0, or 1.
				qtlDad <- geno.pat$geno[[chr]]$data[,i] -1 # Count of lecithotroph alleles: 0, or 1.
				qtlAll <- (1/4)*qtlDad + (3/8)*qtlMom # X-specific G3 additive effect parameterization

				bin.both[i] <- logLik(glmer(binpheno  ~ qtlMom + qtlDad + geno.mat$pheno$qtl1comb + geno.mat$pheno$qtl2comb + (1|momID), family = "binomial"))[1]
				bc.both[i] <- logLik(lmer(bcpheno ~ qtlMom + qtlDad + geno.mat$pheno$qtl1comb + geno.mat$pheno$qtl2comb + (1|dadID) + (1|momID), REML = F))[1]

				bin.summed[i] <- logLik(glmer(binpheno ~ qtlAll + geno.mat$pheno$qtl1comb + geno.mat$pheno$qtl2comb + (1|momID), family = "binomial"))[1]
				bc.summed[i] <- logLik(lmer(bcpheno ~ qtlAll + geno.mat$pheno$qtl1comb + geno.mat$pheno$qtl2comb + (1|dadID) + (1|momID), REML = F))[1]

		}				
	scan.object[which(scan.object$chr == chr),3] <- bin.both		
	scan.object[which(scan.object$chr == chr),4] <- bin.summed
	scan.object[which(scan.object$chr == chr),5] <- bin.null
	scan.object[which(scan.object$chr == chr),6] <- bc.both
	scan.object[which(scan.object$chr == chr),7] <- bc.summed
	scan.object[which(scan.object$chr == chr),8] <- bc.null
	}
	names(scan.object)[c(3:8)] <- c("binBoth","binSummed","binNull","bcBoth","bcSummed","bcNull")
		
	return(scan.object)
}


date()
scan.out <- MZscan3(MatGeno, PatGeno, scan.holder)
date()

scan.out[,9:11] <- NA
scan.out[,9:11] <- c(scan.out$binBoth+scan.out$bcBoth, scan.out$binSummed+scan.out$bcSummed, scan.out$binNull+scan.out$bcNull)
names(scan.out)[9:11] <- c("combBoth", "combSummed", "combNull")

scan.out[,12:20] <- NA

scan.out[,12:20] <- cbind(2*(scan.out$binBoth - scan.out$binNull), 2*(scan.out$binSummed - scan.out$binNull), 2*(scan.out$binBoth - scan.out$binSummed), 2*(scan.out$bcBoth - scan.out$bcNull), 2*(scan.out$bcSummed - scan.out$bcNull), 2*(scan.out$bcBoth - scan.out$bcSummed), 2*(scan.out$combBoth - scan.out$combNull), 2*(scan.out$combSummed - scan.out$combNull), 2*(scan.out$combBoth - scan.out$combSummed))

names(scan.out)[12:20] <- c("binBothChisq", "binSummedChisq", "binBetasChisq", "bcBothChisq", "bcSummedChisq", "bcBetasChisq", "combBothChisq", "combSummedChisq", "combBetasChisq")

ChnumScan3.results <- scan.out
rm(scan.out)

#### End of second forward search.

apply(ChnumScan3.results[,c(3:4,6:7,9:10)], 2, function(x){ChnumScan3.results[which(x == max(x)),]})
# $binBoth
       # chr      pos  binBoth binSummed   binNull    bcBoth  bcSummed    bcNull  combBoth combSummed  combNull binBothChisq binSummedChisq binBetasChisq
# SNP983   1 19.07275 -1825.77 -1826.016 -1830.935 -4181.544 -4181.553 -4182.028 -6007.314  -6007.569 -6012.963     10.32996       9.837597      0.492361
       # bcBothChisq bcSummedChisq bcBetasChisq combBothChisq combSummedChisq combBetasChisq
# SNP983   0.9689733     0.9517884   0.01718488      11.29893        10.78939      0.5095459

# $binSummed
       # chr      pos  binBoth binSummed   binNull    bcBoth  bcSummed    bcNull  combBoth combSummed  combNull binBothChisq binSummedChisq binBetasChisq
# SNP983   1 19.07275 -1825.77 -1826.016 -1830.935 -4181.544 -4181.553 -4182.028 -6007.314  -6007.569 -6012.963     10.32996       9.837597      0.492361
       # bcBothChisq bcSummedChisq bcBetasChisq combBothChisq combSummedChisq combBetasChisq
# SNP983   0.9689733     0.9517884   0.01718488      11.29893        10.78939      0.5095459

# $bcBoth
       # chr      pos   binBoth binSummed   binNull    bcBoth  bcSummed    bcNull  combBoth combSummed  combNull binBothChisq binSummedChisq binBetasChisq
# SNP537   5 18.12664 -1830.208 -1830.248 -1830.935 -4177.016 -4177.709 -4182.028 -6007.223  -6007.957 -6012.963     1.454376       1.374027    0.08034884
       # bcBothChisq bcSummedChisq bcBetasChisq combBothChisq combSummedChisq combBetasChisq
# SNP537    10.02562      8.638154     1.387468         11.48        10.01218       1.467817

# $bcSummed
       # chr      pos   binBoth binSummed   binNull    bcBoth  bcSummed    bcNull  combBoth combSummed  combNull binBothChisq binSummedChisq binBetasChisq
# SNP537   5 18.12664 -1830.208 -1830.248 -1830.935 -4177.016 -4177.709 -4182.028 -6007.223  -6007.957 -6012.963     1.454376       1.374027    0.08034884
       # bcBothChisq bcSummedChisq bcBetasChisq combBothChisq combSummedChisq combBetasChisq
# SNP537    10.02562      8.638154     1.387468         11.48        10.01218       1.467817

# $combBoth
       # chr      pos   binBoth binSummed   binNull    bcBoth  bcSummed    bcNull  combBoth combSummed  combNull binBothChisq binSummedChisq binBetasChisq
# SNP283   2 6.312749 -1825.921  -1830.14 -1830.935 -4178.213 -4181.744 -4182.028 -6004.134  -6011.884 -6012.963     10.02759        1.59019      8.437401
       # bcBothChisq bcSummedChisq bcBetasChisq combBothChisq combSummedChisq combBetasChisq
# SNP283    7.630818     0.5692215     7.061597      17.65841        2.159412         15.499

# $combSummed
       # chr      pos  binBoth binSummed   binNull    bcBoth  bcSummed    bcNull  combBoth combSummed  combNull binBothChisq binSummedChisq binBetasChisq
# SNP983   1 19.07275 -1825.77 -1826.016 -1830.935 -4181.544 -4181.553 -4182.028 -6007.314  -6007.569 -6012.963     10.32996       9.837597      0.492361
       # bcBothChisq bcSummedChisq bcBetasChisq combBothChisq combSummedChisq combBetasChisq
# SNP983   0.9689733     0.9517884   0.01718488      11.29893        10.78939      0.5095459

# permutations for autosomes and X
perm.combo <- c(MatGeno,PatGeno)
combo.dups <- findDupMarkers(perm.combo, adjacent.only = F)
MGnd <- drop.markers(MatGeno, unlist(combo.dups))
PGnd <- drop.markers(PatGeno, unlist(combo.dups))

# How many columns of results?
n.results <- 6
# How many permutations?
n.perms <- 400

#Prepare data for permutation by parent
cross.rows <- row.names(unique(data.frame(MatGeno$pheno$Male_ID, MatGeno$pheno$Female_ID)))
#Make mini cross objects that include each cross only once. 
mini.MGnd <- subset(MGnd, ind = as.numeric(cross.rows))
#make a mini.Gpat cross object:
mini.PGnd <- subset(MGnd, ind = as.numeric(cross.rows))

perm.scan.holder <- scanone(mini.MGnd, pheno = "nChaetae")
perm.scan.holder$lod <- 0
perm.scan.holder[,3:(2+n.results)] <- 0
perm.results <- matrix(nrow = n.perms, ncol = 2*9) # matrix(nrow = n.perms, ncol = n.results)

date()
for(m in 1:n.perms){
	
	male.perm <- data.frame(levels(MatGeno$pheno$Male_ID), levels(MatGeno$pheno$Male_ID))
	# Permutations are stratified by the F1 mother of the F2 fathers, i.e., 1A, 3C, 6F, 8H.
	male.perm[1:10,2] <- sample(male.perm[1:10,1])	
	male.perm[11:24,2] <- sample(male.perm[11:24,1])	
	male.perm[25:32,2] <- sample(male.perm[25:32,1])	
	male.perm[33:53,2] <- sample(male.perm[33:53,1])	

	female.perm <- data.frame(levels(MatGeno$pheno$Female_ID), levels(MatGeno$pheno$Female_ID)) 
	# Permutations are stratified by the F1 mother of the F2 mothers, i.e., 1A, 3C, 6F, 8H.
	female.perm[1:23,2] <- sample(female.perm[1:23,1])	
	female.perm[24:60,2] <- sample(female.perm[24:60,1])		
	female.perm[61:81,2] <- sample(female.perm[61:81,1])	
	female.perm[82:115,2] <- sample(female.perm[82:115,1])	
	
#	perm.G3s <- G3s.nodups
	perm.Gmat <- MGnd
	perm.Gpat <- PGnd
	
	for(i in 1:dim(MatGeno$pheno)[1]){
		permdad <- male.perm[which(male.perm[,1] == MatGeno$pheno$Male_ID[i]),2]
		permmom <- female.perm[which(female.perm[,1] == MatGeno$pheno$Female_ID[i]),2]
		for(j in 1:11){
			perm.Gpat$geno[[j]]$data[i,] <- mini.PGnd$geno[[j]]$data[which(mini.MGnd$pheno$Male_ID == permdad)[1],]
			perm.Gmat$geno[[j]]$data[i,] <- mini.MGnd$geno[[j]]$data[which(mini.MGnd$pheno$Female_ID == permmom)[1],]
		}
	}
	
permscan <- MZscan3(perm.Gmat, perm.Gpat, perm.scan.holder)
permscan[,9:11] <- NA
permscan[,9:11] <- c(permscan$binBoth+ permscan$bcBoth, permscan$binSummed+ permscan$bcSummed, permscan$binNull+ permscan$bcNull)
names(permscan)[9:11] <- c("combBoth", "combSummed", "combNull")

perm.results[m,1:9] <- apply(as.matrix(permscan[permscan$chr != "X",3:(2+9)]), 2, max)
perm.results[m, (9+1):(2*9)] <- apply(as.matrix(permscan[permscan$chr == "X",3:(2+9)]), 2, max)
perm.out <- data.frame(perm.results)
names(perm.out)  <- colnames(permscan)[3:(2+9)]
print(m); print(date());
}

rm(perm.results)
perm.out[,19:30] <- NA

perm.out[,19:30] <- cbind( 2*(perm.out[,1] - perm.out[,3]), 2*(perm.out[,2] - perm.out[,3]), 2*(perm.out[,4] - perm.out[,6]), 2*(perm.out[,5] - perm.out[,6]), 2*(perm.out[,7] - perm.out[,9]), 2*(perm.out[,8] - perm.out[,9]), 2*(perm.out[,10] - perm.out[,12]), 2*(perm.out[,11] - perm.out[,12]), 2*(perm.out[,13] - perm.out[,15]), 2*(perm.out[,14] - perm.out[,15]), 2*(perm.out[,16] - perm.out[,18]), 2*(perm.out[,17] - perm.out[,18]))

names(perm.out)[19:30] <- c("binBothChisq", "binSummedChisq", "bcBothChisq", "bcSummedChisq", "combBothChisq", "combSummedChisq", paste0("X",c("binBothChisq", "binSummedChisq", "bcBothChisq", "bcSummedChisq", "combBothChisq", "combSummedChisq")))


apply(perm.out[,19:30], 2, function(x){quantile(x,0.95)})
    binBothChisq   binSummedChisq      bcBothChisq    bcSummedChisq    combBothChisq  combSummedChisq    XbinBothChisq  XbinSummedChisq     XbcBothChisq 
       12.333200         9.961165        14.985880        12.079124        21.694342        16.644677         8.307000         6.611866        11.851016 
  XbcSummedChisq   XcombBothChisq XcombSummedChisq 
        6.783045        15.962713        10.675099 

perm.out -> ChnumMZScan3Perms
save(ChnumMZScan3Perms, file = "ChnumScan3.400perms.R")
rm(perm.out)

# No additional significant QTL identified.
```
