## Supplementary File 1. Data and Code for "Baby makes three: maternal, paternal, and zygotic genetic effects shape larval phenotypic evolution": CirriScan.R

```
# Cirri trait scans for G3 phenotypes

library(qtl)
library(lme4)

load(file = "G3AnalysisObjects.R")

# Create an empty scanone object to hold the results
scan.holder <- scanone(MatGeno, pheno = "nChaetae")
# This will give a "calc.genoprob" warning; this can be ignored because we are just making an empty scanone object to fill in later.
scan.holder$lod <- 0
scan.holder[,4:5] <- 0

MZscan1 <- function(geno.mat, geno.pat, scan.object) {
	cirr.bin <- geno.mat$pheno$cirri
	dadID <- geno.mat$pheno$Dad_G2_Name
	momID <-  geno.mat$pheno$Mom_G2_Name
	null.LL <- logLik(glmer(cirr.bin ~ 1 + (1|dadID) + (1|momID), family = "binomial"))[1]

		for(j in 1: 10){	# autosomes
		chr <- names(nmar(geno.mat))[j]
		cirri.both <- NULL
		cirri.summed <- NULL

			for(i in 1:(nmar(geno.mat)[chr])){ 							
				qtlMom <- geno.mat$geno[[chr]]$data[,i] -1 # Count of lecithotroph alleles: 0, 1, or 2.
				qtlDad <- geno.pat$geno[[chr]]$data[,i] -1 # Count of lecithotroph alleles: 0, 1, or 2.
				qtlAll <- qtlMom + qtlDad				
		# Fit mixed models		
				cirri.both[i] <- logLik(glmer(cirr.bin ~ qtlMom + qtlDad + (1|dadID) + (1|momID), family = "binomial"))[1]
				cirri.summed[i] <- logLik(glmer(cirr.bin ~ qtlAll + (1|dadID) + (1|momID), family = "binomial"))[1]
		}				
	scan.object[which(scan.object$chr == chr),3] <- cirri.both
	scan.object[which(scan.object$chr == chr),4] <- cirri.summed
	scan.object[which(scan.object$chr == chr),5] <- null.LL
	}

			for(j in 11){	# X only
		chr <- names(nmar(geno.mat))[j]
		cirri.both <- NULL
		cirri.summed <- NULL

			for(i in 1:(nmar(geno.mat)[chr])){ 	
						
#				print(paste(chr, i))	
						
				qtlMom <- geno.mat$geno[[chr]]$data[,i] -1 # Count of lecithotroph alleles: 0, or 1.
				qtlDad <- geno.pat$geno[[chr]]$data[,i] -1 # Count of lecithotroph alleles: 0, or 1.
				qtlAll <- (1/4)*qtlDad + (3/8)*qtlMom # X-specific G3 additive effect parameterization
		# Fit mixed models		
				cirri.both[i] <- logLik(glmer(cirr.bin ~ qtlMom + qtlDad + (1|dadID) + (1|momID), family = "binomial"))[1]
				cirri.summed[i] <- logLik(glmer(cirr.bin ~ qtlAll + (1|dadID) + (1|momID), family = "binomial"))[1]
		}				
	scan.object[which(scan.object$chr == chr),3] <- cirri.both
	scan.object[which(scan.object$chr == chr),4] <- cirri.summed
	scan.object[which(scan.object$chr == chr),5] <- null.LL

	}
	names(scan.object)[3:5] <- c("Both","Summed","Null")
		
	return(scan.object)
}

date()
scan.out <- MZscan1(MatGeno, PatGeno, scan.holder)
date()

scan.out[,6:8] <- NA
scan.out[,6:8] <- cbind(2*(scan.out$Both - scan.out$Null), 2*(scan.out$Summed - scan.out$Null), 2*(scan.out$Both - scan.out$Summed))
names(scan.out)[6:8] <- c("BothChisq", "SummedChisq", "BetasChisq")

CirriScan1.results <- scan.out
rm(scan.out)

#### End of first forward search.
apply(CirriScan1.results[,c(3:4)], 2, function(x){CirriScan1.results[which(x == max(x)),]})
#$Both
#       chr      pos      Both    Summed     Null BothChisq SummedChisq BetasChisq
#SNP913   5 12.07148 -2033.099 -2033.567 -2046.69  27.18267    26.24764  0.9350248
#$Summed
#        chr     pos    Both    Summed     Null BothChisq SummedChisq BetasChisq
#SNP1072   5 7.66355 -2033.2 -2033.234 -2046.69  26.98175    26.91213 0.06961645


# permutations for autosomes and X

perm.combo <- c(MatGeno,PatGeno)
combo.dups <- findDupMarkers(perm.combo, adjacent.only = F)
MGnd <- drop.markers(MatGeno, unlist(combo.dups))
PGnd <- drop.markers(PatGeno, unlist(combo.dups))

# How many columns of results?
n.results <- 3
# How many permutations?
n.perms <- 100

#Prepare data for permutation by parent
cross.rows <- row.names(unique(data.frame(MatGeno$pheno$Male_ID, MatGeno$pheno$Female_ID)))
#Make mini cross objects that include each cross only once. 
mini.MGnd <- subset(MGnd, ind = as.numeric(cross.rows))
#make a mini.Gpat cross object:
mini.PGnd <- subset(MGnd, ind = as.numeric(cross.rows))

perm.scan.holder <- scanone(mini.MGnd, pheno = "nChaetae")
perm.scan.holder$lod <- 0
perm.scan.holder[,3:(2+n.results)] <- 0
perm.results <- matrix(nrow = n.perms, ncol = 2*n.results) # matrix(nrow = n.perms, ncol = n.results)

date()
for(m in 1:n.perms){
	
	male.perm <- data.frame(levels(MatGeno$pheno$Male_ID), levels(MatGeno$pheno$Male_ID))
	# Permutations are stratified by the F1 mother of the F2 fathers, i.e., 1A, 3C, 6F, 8H.
	male.perm[1:10,2] <- sample(male.perm[1:10,1])	
	male.perm[11:24,2] <- sample(male.perm[11:24,1])	
	male.perm[25:32,2] <- sample(male.perm[25:32,1])	
	male.perm[33:53,2] <- sample(male.perm[33:53,1])	

	female.perm <- data.frame(levels(MatGeno$pheno$Female_ID), levels(MatGeno$pheno$Female_ID)) 
	# Permutations are stratified by the F1 mother of the F2 mothers, i.e., 1A, 3C, 6F, 8H.
	female.perm[1:23,2] <- sample(female.perm[1:23,1])	
	female.perm[24:60,2] <- sample(female.perm[24:60,1])		
	female.perm[61:81,2] <- sample(female.perm[61:81,1])	
	female.perm[82:115,2] <- sample(female.perm[82:115,1])	
	
	perm.Gmat <- MGnd
	perm.Gpat <- PGnd
	
	for(i in 1:dim(MatGeno$pheno)[1]){
		permdad <- male.perm[which(male.perm[,1] == MatGeno$pheno$Male_ID[i]),2]
		permmom <- female.perm[which(female.perm[,1] == MatGeno$pheno$Female_ID[i]),2]
		for(j in 1:11){
			perm.Gpat$geno[[j]]$data[i,] <- mini.PGnd$geno[[j]]$data[which(mini.MGnd$pheno$Male_ID == permdad)[1],]
			perm.Gmat$geno[[j]]$data[i,] <- mini.MGnd$geno[[j]]$data[which(mini.MGnd$pheno$Female_ID == permmom)[1],]
		}
	}
	
permscan <- MZscan1(perm.Gmat, perm.Gpat, perm.scan.holder)

perm.results[m,1:n.results] <- apply(as.matrix(permscan[permscan$chr != "X",3:(2+n.results)]), 2, max)
perm.results[m, (n.results+1):(2*n.results)] <- apply(as.matrix(permscan[permscan$chr == "X",3:(2+n.results)]), 2, max)
perm.out <- data.frame(perm.results)
names(perm.out)  <- colnames(permscan)[3:(2+n.results)]
print(m); print(date());
}

rm(perm.results)
perm.out[,7:12] <- NA

perm.out[,7:12] <- cbind(2*(perm.out$Both - perm.out$Null), 2*(perm.out$Summed - perm.out$Null), 2*(perm.out$Both - perm.out$Summed), 2*(perm.out[,4] - perm.out$Null), 2*(perm.out[,5] - perm.out$Null), 2*(perm.out[,4] - perm.out[,5]))
names(perm.out)[7:12] <- c("BothChisq", "SummedChisq", "BetasChisq", "XBothChisq", "XSummedChisq", "XBetasChisq")

apply(perm.out[,7:12], 2, function(x){quantile(x,0.95)})
#   BothChisq  SummedChisq   BetasChisq   XBothChisq XSummedChisq  XBetasChisq 
#   13.410302     9.944934     7.487175     9.840751     6.831397     6.013042 

perm.out -> CirriMZScan1Perms
save(CirriMZScan1Perms, file = "CirriScan1.100perms.R")
rm(perm.out)

# SNP1072 is an additive-effect QTL

#################

cirri.qtl.5.1072 <- MatGeno$geno[[5]]$data[,"SNP1072"] + PatGeno$geno[[5]]$data[,"SNP1072"]
last.column <- dim(MatGeno$pheno)[2]
MatGeno$pheno[,last.column+1] <- cirri.qtl.5.1072
names(MatGeno$pheno)[last.column + 1] <- "cirri.qtl.5.1072"

# After including the QTL covariate, the models have convergence problems or singular fits for the random effects, because the estimated paternal random effect is zero. For easier fitting, we remove the paternal random effect from these models. Note that the likelihoods are identical with the effects included, but the models take longer to run, which is inconvenient for the permutations. 

MZscan2 <- function(geno.mat, geno.pat, scan.object) {
	cirr.bin <- geno.mat$pheno$cirri
	dadID <- geno.mat$pheno$Dad_G2_Name
	momID <-  geno.mat$pheno$Mom_G2_Name
	null.LL <- logLik(glmer(cirr.bin ~ geno.mat$pheno$cirri.qtl.5.1072 + (1|momID), family = "binomial"))[1]

		for(j in 1: 10){	# autosomes
		chr <- names(nmar(geno.mat))[j]
		cirri.both <- NULL
		cirri.summed <- NULL

			for(i in 1:(nmar(geno.mat)[chr])){ 							
				qtlMom <- geno.mat$geno[[chr]]$data[,i] -1 # Count of lecithotroph alleles: 0, 1, or 2.
				qtlDad <- geno.pat$geno[[chr]]$data[,i] -1 # Count of lecithotroph alleles: 0, 1, or 2.
				qtlAll <- qtlMom + qtlDad				
		# Fit mixed models		
				cirri.both[i] <- logLik(glmer(cirr.bin ~ qtlMom + qtlDad + geno.mat$pheno$cirri.qtl.5.1072 + (1|momID), family = "binomial"))[1]
				cirri.summed[i] <- logLik(glmer(cirr.bin ~ qtlAll + geno.mat$pheno$cirri.qtl.5.1072 + (1|momID), family = "binomial"))[1]
		}				
	scan.object[which(scan.object$chr == chr),3] <- cirri.both
	scan.object[which(scan.object$chr == chr),4] <- cirri.summed
	scan.object[which(scan.object$chr == chr),5] <- null.LL
	}

			for(j in 11){	# X only
		chr <- names(nmar(geno.mat))[j]
		cirri.both <- NULL
		cirri.summed <- NULL

			for(i in 1:(nmar(geno.mat)[chr])){ 	
						
#				print(paste(chr, i))	
						
				qtlMom <- geno.mat$geno[[chr]]$data[,i] -1 # Count of lecithotroph alleles: 0, or 1.
				qtlDad <- geno.pat$geno[[chr]]$data[,i] -1 # Count of lecithotroph alleles: 0, or 1.
				qtlAll <- (1/4)*qtlDad + (3/8)*qtlMom # X-specific G3 additive effect parameterization
		# Fit mixed models		
				cirri.both[i] <- logLik(glmer(cirr.bin ~ qtlMom + qtlDad + geno.mat$pheno$cirri.qtl.5.1072 + (1|momID), family = "binomial"))[1]
				cirri.summed[i] <- logLik(glmer(cirr.bin ~ qtlAll + geno.mat$pheno$cirri.qtl.5.1072 + (1|momID), family = "binomial"))[1]
		}				
	scan.object[which(scan.object$chr == chr),3] <- cirri.both
	scan.object[which(scan.object$chr == chr),4] <- cirri.summed
	scan.object[which(scan.object$chr == chr),5] <- null.LL

	}
	names(scan.object)[3:5] <- c("Both","Summed","Null")
		
	return(scan.object)
}

date()
scan.out <- MZscan2(MatGeno, PatGeno, scan.holder)
date()

scan.out[,6:8] <- NA
scan.out[,6:8] <- cbind(2*(scan.out$Both - scan.out$Null), 2*(scan.out$Summed - scan.out$Null), 2*(scan.out$Both - scan.out$Summed))
names(scan.out)[6:8] <- c("BothChisq", "SummedChisq", "BetasChisq")

CirriScan2.results <- scan.out
rm(scan.out)

#### End of second forward search.

apply(CirriScan2.results[,c(3:4)], 2, function(x){CirriScan2.results[which(x == max(x)),]})
#$Both
#       chr      pos      Both    Summed      Null BothChisq SummedChisq BetasChisq
#SNP894   2 27.83767 -2029.039 -2029.441 -2033.234  8.390619    7.587144  0.8034743
#SNP365   2 28.19860 -2029.039 -2029.441 -2033.234  8.390619    7.587144  0.8034743
#$Summed
#       chr      pos      Both    Summed      Null BothChisq SummedChisq BetasChisq
#SNP894   2 27.83767 -2029.039 -2029.441 -2033.234  8.390619    7.587144  0.8034743
#SNP365   2 28.19860 -2029.039 -2029.441 -2033.234  8.390619    7.587144  0.8034743

# permutations for autosomes and X
perm.combo <- c(MatGeno,PatGeno)
combo.dups <- findDupMarkers(perm.combo, adjacent.only = F)
MGnd <- drop.markers(MatGeno, unlist(combo.dups))
PGnd <- drop.markers(PatGeno, unlist(combo.dups))

# How many columns of results?
n.results <- 3
# How many permutations?
n.perms <- 100

#Prepare data for permutation by parent
cross.rows <- row.names(unique(data.frame(MatGeno$pheno$Male_ID, MatGeno$pheno$Female_ID)))
#Make mini cross objects that include each cross only once. 
mini.MGnd <- subset(MGnd, ind = as.numeric(cross.rows))
#make a mini.Gpat cross object:
mini.PGnd <- subset(MGnd, ind = as.numeric(cross.rows))

perm.scan.holder <- scanone(mini.MGnd, pheno = "nChaetae")
perm.scan.holder$lod <- 0
perm.scan.holder[,3:(2+n.results)] <- 0
perm.results <- matrix(nrow = n.perms, ncol = 2*n.results) # matrix(nrow = n.perms, ncol = n.results)

date()
for(m in 1:n.perms){
	
	male.perm <- data.frame(levels(MatGeno$pheno$Male_ID), levels(MatGeno$pheno$Male_ID))
	# Permutations are stratified by the F1 mother of the F2 fathers, i.e., 1A, 3C, 6F, 8H.
	male.perm[1:10,2] <- sample(male.perm[1:10,1])	
	male.perm[11:24,2] <- sample(male.perm[11:24,1])	
	male.perm[25:32,2] <- sample(male.perm[25:32,1])	
	male.perm[33:53,2] <- sample(male.perm[33:53,1])	

	female.perm <- data.frame(levels(MatGeno$pheno$Female_ID), levels(MatGeno$pheno$Female_ID)) 
	# Permutations are stratified by the F1 mother of the F2 mothers, i.e., 1A, 3C, 6F, 8H.
	female.perm[1:23,2] <- sample(female.perm[1:23,1])	
	female.perm[24:60,2] <- sample(female.perm[24:60,1])		
	female.perm[61:81,2] <- sample(female.perm[61:81,1])	
	female.perm[82:115,2] <- sample(female.perm[82:115,1])	
	
	perm.Gmat <- MGnd
	perm.Gpat <- PGnd
	
	for(i in 1:dim(MatGeno$pheno)[1]){
		permdad <- male.perm[which(male.perm[,1] == MatGeno$pheno$Male_ID[i]),2]
		permmom <- female.perm[which(female.perm[,1] == MatGeno$pheno$Female_ID[i]),2]
		for(j in 1:11){
			perm.Gpat$geno[[j]]$data[i,] <- mini.PGnd$geno[[j]]$data[which(mini.MGnd$pheno$Male_ID == permdad)[1],]
			perm.Gmat$geno[[j]]$data[i,] <- mini.MGnd$geno[[j]]$data[which(mini.MGnd$pheno$Female_ID == permmom)[1],]
		}
	}
	
permscan <- MZscan2(perm.Gmat, perm.Gpat, perm.scan.holder)

perm.results[m,1:n.results] <- apply(as.matrix(permscan[permscan$chr != "X",3:(2+n.results)]), 2, max)
perm.results[m, (n.results+1):(2*n.results)] <- apply(as.matrix(permscan[permscan$chr == "X",3:(2+n.results)]), 2, max)
perm.out <- data.frame(perm.results)
names(perm.out)  <- colnames(permscan)[3:(2+n.results)]
print(m); print(date());
}

rm(perm.results)
perm.out[,7:12] <- NA

perm.out[,7:12] <- cbind(2*(perm.out$Both - perm.out$Null), 2*(perm.out$Summed - perm.out$Null), 2*(perm.out$Both - perm.out$Summed), 2*(perm.out[,4] - perm.out$Null), 2*(perm.out[,5] - perm.out$Null), 2*(perm.out[,4] - perm.out[,5]))
names(perm.out)[7:12] <- c("BothChisq", "SummedChisq", "BetasChisq", "XBothChisq", "XSummedChisq", "XBetasChisq")

apply(perm.out[,7:12], 2, function(x){quantile(x,0.95)})
#   BothChisq  SummedChisq   BetasChisq   XBothChisq XSummedChisq  XBetasChisq 
#   12.212682     8.780852     5.290903     9.139501     7.115481     6.077243 

perm.out -> CirriMZScan2Perms
save(CirriMZScan2Perms, file = "CirriScan2.100perms.R")
rm(perm.out)

# No additional significant QTL from the second scan.
```
