## Supplementary File 1. Data and Code for "Baby makes three: maternal, paternal, and zygotic genetic effects shape larval phenotypic evolution": DataFigures.R

```
# These code blocks reproduce the data figures in the manuscript (Figures 2-6)

#############################
# Figure 2: chaetae number as a function of larval size
load(file = "G3AnalysisObjects.R")
plot(I(MatGeno$pheno$nChaetae + rnorm(length(MatGeno$pheno$nChaetae), 0, 0.1)) ~MatGeno$pheno$area, pch = 19, cex = 0.3, ylab = "number of larval chaetae", xlab = "larval size (square microns)")
abline(h = seq(-0.5, 24.5, 1), col = "gray")

#############################
# Figure 3: Distribution of chaetae number and chaetae length
load(file = "G3AnalysisObjects.R")
par(mfrow = c(1,2))
hist(MatGeno$pheno$nChaetae, br= seq(-0.5,24.5,1), col = "red", main = "", ylab = "Frequency", xlab = "number of chaetae")
hist(MatGeno$pheno$ChLen, br = seq(0,300,10), col = "red", main = "", ylab = "Frequency", xlab = "length of chaetae")

#############################
# Figure 4: Joint G2-G3 QTL scans for chaetae number, chaetae length, and anal cirri
# File JointScanResults.R contains the linkage mapping generated by the scripts JointChNumScan.R, JointCirriScan.R, and JointChLenScan.R
library(qtl)
load(file = "JointScanResults.R")
# names(JointScanResults)
# [1] "chr"     "pos"     "Cirri.1" "Cirri.2" "Chlen.1" "Chlen.2" "Chlen.3" "Chnum.1" "Chnum.2" "Chnum.3"

# Cirri
JointScanResults[JointScanResults$chr != 5,"Cirri.1"] <- -10
plot(JointScanResults, lod = 1, col = "purple", ylim = c(0,135), ylab = "Test Statistic")
plot(JointScanResults, lod = 2, col = "purple", ylim = c(0,135), lwd = 0.5, add = T)

# Chaetae length
JointScanResults[JointScanResults$chr != 3, "Chlen.1"] <- -10
JointScanResults[JointScanResults$chr != 9, "Chlen.2"] <- -10
plot(JointScanResults, lod = 3, col = "blue", ylim = c(0,135), add = T)
plot(JointScanResults, lod = 4, col = "blue", ylim = c(0,135), add = T)
plot(JointScanResults, lod = 5, col = "blue", ylim = c(0,135), lwd = 0.5, add = T)

# Chaetae number
JointScanResults[JointScanResults$chr != 8, "Chnum.1"] <- -10
JointScanResults[JointScanResults$chr != 3, "Chnum.2"] <- -10
plot(JointScanResults, lod = 6, col = "red", ylim = c(0,135), add = T)
plot(JointScanResults, lod = 7, col = "red", ylim = c(0,135), add = T)
plot(JointScanResults, lod = 8, col = "red", ylim = c(0,135), lwd = 0.5, add = T)

abline(h = c(14.33, 16.13, 20.26), lty = 3, lwd = 0.75, col =c("purple", "blue", "red"))

#############################
# Figure 5: larval size QTL scans
# File AreaScanResults.R contains the linkage mapping generated by the script AreaScanAndTestsForDominanceEtc.R
load(file = "AreaScanResults.R")
#names(AreaScanResults)
#[1] "chr"   "pos"   "Scan1" "Scan2" "Scan3" "Scan4" "Scan5"

AreaScanResults[AreaScanResults$chr != 7, "Scan1"] <- -10
AreaScanResults[AreaScanResults$chr != 6, "Scan2"] <- -10
AreaScanResults[AreaScanResults$chr != "X", "Scan3"] <- -10
AreaScanResults[AreaScanResults$chr != 9, "Scan4"] <- -10
plot(AreaScanResults, lod = 1, col = "black", ylim = c(0,40), ylab = "Test Statistic")
plot(AreaScanResults, lod = 2, col = "black", ylim = c(0,40), add = T)
plot(AreaScanResults, lod = 3, col = "black", ylim = c(0,40), add = T)
plot(AreaScanResults, lod = 4, col = "black", ylim = c(0,40), add = T)
plot(AreaScanResults, lod = 5, col = "gray", ylim = c(0,40), add = T)

abline(h = 13.78, lty = 3, lwd = 0.75, col = "gray")

#############################
# Figure 6: effects of larval-size QTLs
library(qtl)
library(lme4)
load(file = "G3AnalysisObjects.R")

 	dadID <- MatGeno$pheno$Dad_G2_Name
	momID <-  MatGeno$pheno$Mom_G2_Name
	area.pheno <- MatGeno$pheno$area
	
	# Area QTL are parental-effect loci
	qtl1.area <- MatGeno$geno[[7]]$data[,"SNP68"] - 1
	qtl2.area <- MatGeno$geno[[6]]$data[,"SNP93"] - 1
	qtl3.areaP <- PatGeno$geno[["X"]]$data[,"SNP1249"] - 1.  
	qtl3.areaM <- MatGeno$geno[["X"]]$data[,"SNP1249"] - 1.  
	qtl4.areaP <- PatGeno$geno[[9]]$data[,"SNP1244"] - 1
	qtl4.areaM <- MatGeno$geno[[9]]$data[,"SNP1244"] - 1

	# Dominance for qtl2 
	qtl2.area.h <- qtl2.area
	qtl2.area.h[qtl2.area == 2] <- 0

	get.lecitho.proportion <- function(genotypes, chromosomes = c(1:10)){ 
	allelecountbychr <- matrix(nrow = dim(genotypes$pheno)[1], ncol = length(chromosomes))	
			for(j in 1: length(chromosomes)){
			allelecountbychr[,j] <- apply(genotypes$geno[[j]]$data -1, 1, sum)
			}
	TotalAlleleCount <- apply(allelecountbychr, 1, sum)
	TotalAlleleCount/(2*sum(nmar(genotypes)[c(chromosomes)]))
	}
	
	area.dom2 <- lmer(area.pheno  ~ 1 + qtl1.area + qtl2.area + qtl3.areaP + qtl3.areaM + qtl4.areaP + qtl4.areaM + qtl2.area.h + (1|momID), REML = F)
	
	LpropM.area <- get.lecitho.proportion(MatGeno, c(1:5,8,10)) # excludes 6, 7, 9, and X

# Fit the models
# Because we're not using likelihood ratio tests for this, we can run the models with REML 
area.polyMx <- lmer(area.pheno  ~ 1 + qtl1.area + qtl2.area + qtl3.areaP + qtl3.areaM + qtl4.areaP + qtl4.areaM + qtl2.area.h + LpropM.area + (1|momID))
# This is the model selected by successive tests as described in the manuscript

# Here are the leave-one-out models
area.NoQTL1 <- lmer(area.pheno  ~ 1 + qtl2.area + qtl3.areaP + qtl3.areaM + qtl4.areaP + qtl4.areaM + qtl2.area.h + LpropM.area + (1|momID))
area.NoQTL2 <- lmer(area.pheno  ~ 1 + qtl1.area + qtl3.areaP + qtl3.areaM + qtl4.areaP + qtl4.areaM + LpropM.area + (1|momID))
area.NoQTL3P <- lmer(area.pheno  ~ 1 + qtl1.area + qtl2.area + qtl3.areaM + qtl4.areaP + qtl4.areaM + qtl2.area.h + LpropM.area + (1|momID))
area.NoQTL3M <- lmer(area.pheno  ~ 1 + qtl1.area + qtl2.area + qtl3.areaP + qtl4.areaP + qtl4.areaM + qtl2.area.h + LpropM.area + (1|momID))
area.NoQTL4P <- lmer(area.pheno  ~ 1 + qtl1.area + qtl2.area + qtl3.areaP + qtl3.areaM + qtl4.areaM + qtl2.area.h + LpropM.area + (1|momID))
area.NoQTL4M <- lmer(area.pheno  ~ 1 + qtl1.area + qtl2.area + qtl3.areaP + qtl3.areaM + qtl4.areaP + qtl2.area.h + LpropM.area + (1|momID))
area.NoPoly <- lmer(area.pheno  ~ 1 + qtl1.area + qtl2.area + qtl3.areaP + qtl3.areaM + qtl4.areaP + qtl4.areaM + qtl2.area.h  + (1|momID))

# Get the genotypes for each family
qtl1.geno <- NULL
qtl2.geno <- NULL
qtl3P.geno <- NULL
qtl3M.geno <- NULL
qtl4P.geno <- NULL
qtl4M.geno <- NULL
poly.geno <- NULL

for(i in 1:115){
	qtl1.geno[i] <- qtl1.area[which(MatGeno$pheno$Mom_G2_Name == row.names(ranef(area.polyMx)$momID)[i])[1]]
	qtl2.geno[i] <- qtl2.area[which(MatGeno$pheno$Mom_G2_Name == row.names(ranef(area.polyMx)$momID)[i])[1]]
	qtl3P.geno[i] <- qtl3.areaP[which(MatGeno$pheno$Mom_G2_Name == row.names(ranef(area.polyMx)$momID)[i])[1]]
	qtl3M.geno[i] <- qtl3.areaM[which(MatGeno$pheno$Mom_G2_Name == row.names(ranef(area.polyMx)$momID)[i])[1]]
	qtl4P.geno[i] <- qtl4.areaP[which(MatGeno$pheno$Mom_G2_Name == row.names(ranef(area.polyMx)$momID)[i])[1]]
	qtl4M.geno[i] <- qtl4.areaM[which(MatGeno$pheno$Mom_G2_Name == row.names(ranef(area.polyMx)$momID)[i])[1]]
	poly.geno[i] <- LpropM.area[which(MatGeno$pheno$Mom_G2_Name == row.names(ranef(area.polyMx)$momID)[i])[1]]
}

# Plots of BLUPs that control for all the other effects, with boxplots
par(mfrow = c(1,6))
boxplot(ranef(area.NoQTL1)$momID[,1] ~ qtl1.geno, ylim = c(-11000,11000), outline = F)
stripchart(ranef(area.NoQTL1)$momID[,1] ~ qtl1.geno, method = "jitter", jitter = 0.1, vertical = T, pch = 19, cex= 0.4, ylim = c(-11000,11000), add = T)
abline(lm(ranef(area.NoQTL1)$momID[,1] ~ I(qtl1.geno+1)))

boxplot(ranef(area.NoQTL2)$momID[,1] ~ qtl2.geno, ylim = c(-11000,11000), outline = F)
stripchart(ranef(area.NoQTL2)$momID[,1] ~ qtl2.geno, method = "jitter", jitter = 0.1, vertical = T, pch = 19, cex= 0.4, ylim = c(-11000,11000), add = T)
abline(lm(ranef(area.NoQTL2)$momID[,1] ~ I(qtl2.geno+1)))

boxplot(ranef(area.NoQTL3P)$momID[,1] ~ qtl3P.geno , ylim = c(-11000,11000), outline = F)
stripchart(ranef(area.NoQTL3P)$momID[,1] ~ qtl3P.geno, method = "jitter", jitter = 0.1, vertical = T, pch = 19, cex= 0.4, ylim = c(-11000,11000), add =T)
abline(lm(ranef(area.NoQTL3P)$momID[,1] ~ I(qtl3P.geno+1)))

boxplot(ranef(area.NoQTL3M)$momID[,1] ~ qtl3M.geno , ylim = c(-11000,11000), outline = F)
stripchart(ranef(area.NoQTL3M)$momID[,1] ~ qtl3M.geno, method = "jitter", jitter = 0.1, vertical = T, pch = 19, cex= 0.4, ylim = c(-11000,11000), add = T)
abline(lm(ranef(area.NoQTL3M)$momID[,1] ~ I(qtl3M.geno+1)))

boxplot(ranef(area.NoQTL4P)$momID[,1] ~ qtl4P.geno , ylim = c(-11000,11000), outline = F)
stripchart(ranef(area.NoQTL4P)$momID[,1] ~ qtl4P.geno, method = "jitter", jitter = 0.1, vertical = T, pch = 19, cex= 0.4, ylim = c(-11000,11000), add = T)
abline(lm(ranef(area.NoQTL4P)$momID[,1] ~ I(qtl4P.geno+1)))

boxplot(ranef(area.NoQTL4M)$momID[,1] ~ qtl4M.geno , ylim = c(-11000,11000), outline = F)
stripchart(ranef(area.NoQTL4M)$momID[,1] ~ qtl4M.geno, method = "jitter", jitter = 0.1, vertical = T, pch = 19, cex= 0.4, ylim = c(-11000,11000), add =T)
abline(lm(ranef(area.NoQTL4M)$momID[,1] ~ I(qtl4M.geno+1)))

par(mfrow =c(1,1))
plot(ranef(area.NoPoly)$momID[,1] ~ poly.geno, ylim = c(-11000,11000), pch = 19, cex = 0.4, xlim = c(0,1))
abline(lm(ranef(area.NoPoly)$momID[,1] ~ poly.geno))
```
