## Supplementary File 1. Data and Code for "Baby makes three: maternal, paternal, and zygotic genetic effects shape larval phenotypic evolution": HeritabilityEstimates.R

```
# Heritability estimates from parent-offspring regression

# Procedure overview:
# Generate a dataset that lacks NAs, then perform mixed-model regression with mid-parent phenotype as fixed effect
# Test for significance by comparing to a null model with only intercept as fixed effect.
# If significant, estimate heritability and its standard error
# Test for parental effects by comparing the mid-parent model to one with separate maternal and paternal phenotypes
# Test for residual heritability after accounting for QTLs by comparing a model with QTLs to one without.

########################
# Chaetae length, among animals with chaetae
chl <- subset(MatGeno, ind = !is.na(MatGeno$pheno$ChLen))
chl <- subset(chl, ind = !is.na(chl$pheno$Dad_ChLen))
chl <- subset(chl, ind = !is.na(chl$pheno$Mom_ChLen))

null.chl <- lmer(chl$pheno$ChLen ~ 1 + (1|chl$pheno$Mom_G2_Name) + (1|chl$pheno$Dad_G2_Name) + (1|chl$pheno$Mom_Fam) + (1|chl$pheno$Dad_Fam))

full.chl <- lmer(chl$pheno$ChLen ~ I((chl$pheno$Dad_ChLen + chl$pheno$Mom_ChLen)/2) + (1|chl$pheno$Mom_G2_Name) + (1|chl$pheno$Dad_G2_Name) + (1|chl$pheno$Mom_Fam) + (1|chl$pheno$Dad_Fam))

# Is there significant heritability?
#         npar   AIC   BIC logLik deviance  Chisq Df Pr(>Chisq)    
#null.chl    6 21905 21940 -10947    21893                         
#full.chl    7 21890 21930 -10938    21876 17.576  1  2.761e-05 ***

# Yes.

# Estimate the heritability from midparent regression:
#Linear mixed model fit by REML ['lmerMod']
#Formula: chl$pheno$ChLen ~ I((chl$pheno$Dad_ChLen + chl$pheno$Mom_ChLen)/2) + (1 | chl$pheno$Mom_G2_Name) + (1 | chl$pheno$Dad_G2_Name) +      (1 | chl$pheno$Mom_Fam) + (1 | chl$pheno$Dad_Fam)
#REML criterion at convergence: 21874.47
#Random effects:
# Groups                Name        Std.Dev. 
# chl$pheno$Mom_G2_Name (Intercept) 16.449255
# chl$pheno$Dad_G2_Name (Intercept) 12.946202
# chl$pheno$Mom_Fam     (Intercept)  3.764129
# chl$pheno$Dad_Fam     (Intercept)  0.002457
# Residual                          36.115165
#Number of obs: 2169, groups:  chl$pheno$Mom_G2_Name, 79; chl$pheno$Dad_G2_Name, 42; chl$pheno$Mom_Fam, 4; chl$pheno$Dad_Fam, 4
#Fixed Effects:
#                                     (Intercept)  I((chl$pheno$Dad_ChLen + chl$pheno$Mom_ChLen)/2)  
#                                         79.8206                                            0.4305  

# h2 = 0.4305
# standard error:
sqrt(vcov(full)[2,2])
#[1] 0.09680211

# estimates from each parent are very similar (these are both estimates of heritability/2)
biparentchl <- lmer(chl$pheno$ChLen ~ chl$pheno$Dad_ChLen + chl$pheno$Mom_ChLen + (1|chl$pheno$Mom_G2_Name) + (1|chl$pheno$Dad_G2_Name) + (1|chl$pheno$Mom_Fam) + (1|chl$pheno$Dad_Fam)) 
#Fixed Effects:
#        (Intercept)  chl$pheno$Dad_ChLen  chl$pheno$Mom_ChLen  
#            78.6601               0.2384               0.2018  
# standard error:
sqrt(vcov(biparentchl))
#                    (Intercept) chl$pheno$Dad_ChLen chl$pheno$Mom_ChLen
#(Intercept)            13.50226                 NaN                 NaN
#chl$pheno$Dad_ChLen         NaN          0.07484492          0.02408498
#chl$pheno$Mom_ChLen         NaN          0.02408498          0.05811363

# Not better than midparent
#anova(biparentchl,full)
#            npar   AIC   BIC logLik deviance  Chisq Df Pr(>Chisq)
#full           7 21890 21930 -10938    21876                     
#biparentchl    8 21892 21937 -10938    21876 0.1948  1     0.6589


# Accounting for the major-effect QTL

Pchl <- subset(PatGeno, ind = !is.na(PatGeno$pheno$ChLen))
Pchl <- subset(Pchl, ind = !is.na(Pchl$pheno$Dad_ChLen))
Pchl <- subset(Pchl, ind = !is.na(Pchl$pheno$Mom_ChLen))

	get.expected.heterozygosity <- function(geno.mat, geno.pat, chrom, snpname){
	snp.M <- geno.mat$geno[[chrom]]$data[,snpname] - 2
	snp.P <- geno.pat$geno[[chrom]]$data[,snpname] - 2
	snp <- snp.M + snp.P
	snp.het <- snp
	snp.het[snp %in% c(-2,2)] <- 0
	snp.het[snp.M == 0] <- 0.5
	snp.het[snp.P == 0] <- 0.5
	snp.het[snp.M == -1 & snp.P == 1] <- 1
	snp.het[snp.M == 1 & snp.P == -1] <- 1
	return(snp.het)
}

chlQ1 <- chl$geno[[9]]$data[,"SNP272"] + Pchl$geno[[9]]$data[,"SNP272"]
chlQ2 <- chl$geno[[3]]$data[,"SNP497"] + Pchl$geno[[3]]$data[,"SNP497"]
chlQ2h <- get.expected.heterozygosity(chl, Pchl, 3, "SNP497")

null.chl.qtls <- lmer(chl$pheno$ChLen ~ 1 + chlQ1 + chlQ2 + chlQ2h + (1|chl$pheno$Mom_G2_Name) + (1|chl$pheno$Dad_G2_Name) + (1|chl$pheno$Mom_Fam) + (1|chl$pheno$Dad_Fam))

full.chl.qtls <- lmer(chl$pheno$ChLen ~ I((chl$pheno$Dad_ChLen + chl$pheno$Mom_ChLen)/2) + chlQ1 + chlQ2 + chlQ2h + (1|chl$pheno$Mom_G2_Name) + (1|chl$pheno$Dad_G2_Name) + (1|chl$pheno$Mom_Fam) + (1|chl$pheno$Dad_Fam))

# Is there evidence that the residual variation is heritable?
anova(full.chl.qtls, null.chl.qtls)
#              npar   AIC   BIC logLik deviance  Chisq Df Pr(>Chisq)
#null.chl.qtls    9 21846 21897 -10914    21828                     
#full.chl.qtls   10 21847 21904 -10914    21827 0.1091  1     0.7412

# No.

#Linear mixed model fit by REML ['lmerMod']
#Formula: chl$pheno$ChLen ~ I((chl$pheno$Dad_ChLen + chl$pheno$Mom_ChLen)/2) +  
#    chlQ1 + chlQ2 + chlQ2h + (1 | chl$pheno$Mom_G2_Name) + (1|chl$pheno$Dad_G2_Name) + (1 | chl$pheno$Mom_Fam) + (1 | chl$pheno$Dad_Fam)

#REML criterion at convergence: 21813.3
#Scaled residuals: 
#    Min      1Q  Median      3Q     Max 
#-3.4919 -0.6791  0.0165  0.6744  3.5582 

#Random effects:
# Groups                Name        Variance Std.Dev.
# chl$pheno$Mom_G2_Name (Intercept)   88.229  9.393  
# chl$pheno$Dad_G2_Name (Intercept)  117.924 10.859  
# chl$pheno$Mom_Fam     (Intercept)   21.366  4.622  
# chl$pheno$Dad_Fam     (Intercept)    1.714  1.309  
# Residual                          1306.366 36.144  
#Number of obs: 2169, groups:  chl$pheno$Mom_G2_Name, 79; chl$pheno$Dad_G2_Name, 42; chl$pheno$Mom_Fam, 4; chl$pheno$Dad_Fam, 4

#Fixed effects:
#                                                  Estimate Std. Error t value
#(Intercept)                                      262.42058   25.04688  10.477
#I((chl$pheno$Dad_ChLen + chl$pheno$Mom_ChLen)/2)   0.03038    0.08875   0.342
#chlQ1                                            -14.90949    2.85627  -5.220
#chlQ2                                            -14.09178    2.35157  -5.992
#chlQ2h                                           -16.83753    9.01901  -1.867

sqrt(vcov(full.chl.qtls))[2,2]
#[1] 0.08875204

# After accounting for QTL effects, residual heritability is 0.03 +/- 0.09, p = 0.74

########################
# Larval area

ar <- subset(MatGeno, ind = !is.na(MatGeno$pheno$area))
ar <- subset(ar, ind = !is.na(ar$pheno$Mom_area))
ar <- subset(ar, ind = !is.na(ar$pheno$Dad_area))

null.ar <- lmer(ar$pheno$area/10^5 ~ 1 + (1|ar$pheno$Mom_G2_Name) + (1|ar$pheno$Dad_G2_Name) + (1|ar$pheno$Mom_Fam) + (1|ar$pheno$Dad_Fam))

full.ar <- lmer(ar$pheno$area/10^5 ~ I((ar$pheno$Dad_area/10^5 + ar$pheno$Mom_area/10^5)/2) + (1|ar$pheno$Mom_G2_Name) + (1|ar$pheno$Dad_G2_Name) + (1|ar$pheno$Mom_Fam) + (1|ar$pheno$Dad_Fam))

# Is there significant heritability?
anova(full.ar, null.ar)
#        npar    AIC    BIC logLik deviance  Chisq Df Pr(>Chisq)
#null.ar    6 -11733 -11695 5872.3   -11745                     
#full.ar    7 -11732 -11688 5873.1   -11746 1.6996  1     0.1923

# No

########################
# Number of chaetae
nch <- subset(MatGeno, ind = !is.na(MatGeno$pheno$nChaetae))
nch <- subset(nch, ind = !is.na(nch$pheno$Mom_nChaetae))
nch <- subset(nch, ind = !is.na(nch$pheno$Dad_nChaetae))

# Observed distribution

null.nch <- lmer(nch$pheno$nChaetae ~ 1 + (1|nch$pheno$Mom_G2_Name) + (1|nch$pheno$Dad_G2_Name) + (1|nch$pheno$Mom_Fam) + (1|nch$pheno$Dad_Fam))

full.nch <- lmer(nch$pheno$nChaetae ~ 1 + I((nch$pheno$Mom_nChaetae + nch$pheno$Dad_nChaetae)/2) + (1|nch$pheno$Mom_G2_Name) + (1|nch$pheno$Dad_G2_Name) + (1|nch$pheno$Mom_Fam) + (1|nch$pheno$Dad_Fam))

anova(full.nch, null.nch)
#         npar   AIC   BIC logLik deviance  Chisq Df Pr(>Chisq)    
#null.nch    6 20620 20658 -10304    20608                         
#full.nch    7 20603 20647 -10294    20589 19.136  1  1.218e-05 ***


#Linear mixed model fit by REML ['lmerMod']
#Formula: nch$pheno$nChaetae ~ 1 + I((nch$pheno$Mom_nChaetae + nch$pheno$Dad_nChaetae)/2) + (1 | nch$pheno$Mom_G2_Name) + (1 | nch$pheno$Dad_G2_Name) +      (1 | nch$pheno$Mom_Fam) + (1 | nch$pheno$Dad_Fam)
#REML criterion at convergence: 20592.79
#Random effects:
# Groups                Name        Std.Dev. 
# nch$pheno$Mom_G2_Name (Intercept) 1.8003766
# nch$pheno$Dad_G2_Name (Intercept) 0.9062903
# nch$pheno$Mom_Fam     (Intercept) 0.0002710
# nch$pheno$Dad_Fam     (Intercept) 0.0001533
# Residual                          3.2740910
#Number of obs: 3899, groups:  nch$pheno$Mom_G2_Name, 114; nch$pheno$Dad_G2_Name, 53; nch$pheno$Mom_Fam, 4; nch$pheno$Dad_Fam, 4
#Fixed Effects:
#                                           (Intercept)  I((nch$pheno$Mom_nChaetae + nch$pheno$Dad_nChaetae)/2)  
#                                                1.8222                                                  0.4315  

sqrt(vcov(full.nch)[2,2])
#[1] 0.09360339

# heritability = 0.43 +/- 0.09

biparent.nch <- lmer(nch$pheno$nChaetae ~ 1 + nch$pheno$Mom_nChaetae + nch$pheno$Dad_nChaetae + (1|nch$pheno$Mom_G2_Name) + (1|nch$pheno$Dad_G2_Name) + (1|nch$pheno$Mom_Fam) + (1|nch$pheno$Dad_Fam))

anova(biparent.nch, full.nch)
#             npar   AIC   BIC logLik deviance  Chisq Df Pr(>Chisq)
#full.nch        7 20603 20647 -10294    20589                     
#biparent.nch    8 20604 20654 -10294    20588 0.9496  1     0.3298
# Not better with separate parental effects


# Accounting for known QTLs

Pnch <- subset(PatGeno, ind = !is.na(PatGeno$pheno$nChaetae))
Pnch <- subset(Pnch, ind = !is.na(Pnch$pheno$Mom_nChaetae))
Pnch <- subset(Pnch, ind = !is.na(Pnch$pheno$Dad_nChaetae))

		chnQ1 <- nch$geno[[8]]$data[,"SNP231"] + Pnch$geno[[8]]$data[,"SNP231"] -2
		chnQ2 <- nch$geno[[3]]$data[,"SNP497"] + Pnch$geno[[3]]$data[,"SNP497"] -2
		chnQ1h <- get.expected.heterozygosity(nch, Pnch, 8, "SNP231")
		chnQ2h <- get.expected.heterozygosity(nch, Pnch, 3, "SNP497")

null.nch.qtl <- lmer(nch$pheno$nChaetae ~ chnQ1 + chnQ2 + chnQ1h + chnQ2h + (1|nch$pheno$Mom_G2_Name) + (1|nch$pheno$Dad_G2_Name) + (1|nch$pheno$Mom_Fam) + (1|nch$pheno$Dad_Fam))


full.nch.qtl <- lmer(nch$pheno$nChaetae ~ I((nch$pheno$Mom_nChaetae + nch$pheno$Dad_nChaetae)/2) + chnQ1 + chnQ2 + chnQ1h + chnQ2h + (1|nch$pheno$Mom_G2_Name) + (1|nch$pheno$Dad_G2_Name) + (1|nch$pheno$Mom_Fam) + (1|nch$pheno$Dad_Fam))

anova(full.nch.qtl, null.nch.qtl)
#             npar   AIC   BIC logLik deviance  Chisq Df Pr(>Chisq)
#null.nch.qtl   10 20555 20618 -10268    20535                     
#full.nch.qtl   11 20557 20626 -10268    20535 0.0244  1     0.8758

#Linear mixed model fit by REML ['lmerMod']
#Formula: nch$pheno$nChaetae ~ I((nch$pheno$Mom_nChaetae + nch$pheno$Dad_nChaetae)/2) + chnQ1 + chnQ2 + chnQ1h + chnQ2h + (1 | nch$pheno$Mom_G2_Name) + (1 | nch$pheno$Dad_G2_Name) + (1 | nch$pheno$Mom_Fam) + (1 |nch$pheno$Dad_Fam)
#REML criterion at convergence: 20540.48
#Random effects:
# Groups                Name        Std.Dev.
# nch$pheno$Mom_G2_Name (Intercept) 1.4839  
# nch$pheno$Dad_G2_Name (Intercept) 0.3742  
# nch$pheno$Mom_Fam     (Intercept) 0.0000  
# nch$pheno$Dad_Fam     (Intercept) 0.3177  
# Residual                          3.2740  
#Number of obs: 3899, groups:  nch$pheno$Mom_G2_Name, 114; nch$pheno$Dad_G2_Name, 53; nch$pheno$Mom_Fam, 4; nch$pheno$Dad_Fam, 4
#Fixed Effects:
#                                           (Intercept)  I((nch$pheno$Mom_nChaetae + nch$pheno$Dad_nChaetae)/2)  
#                                                7.4706                                                  0.0153  
#                                                 chnQ1                                                   chnQ2  
#                                               -1.5301                                                 -1.0986  
#                                                chnQ1h                                                  chnQ2h  
#                                                1.6693                                                  0.4461  

sqrt(vcov(full.nch.qtl))[2,2]
#[1] 0.09117138

# After accounting for QTL effects, residual heritability is 0.015 +/- 0.09, p = 0.74
```
