## Supplementary File 1. Data and Code for "Baby makes three: maternal, paternal, and zygotic genetic effects shape larval phenotypic evolution": JointChLenScan.R

```
# Combine likelihoods from G2s and G3s

library(qtl)
library(lme4)

load(file = "G3AnalysisObjects.R")
# [1] "dups"       "MatGeno"    "PatGeno"    "Sb.deduped"


scan.holder <- scanone(Sb.deduped, pheno = "nchaetae")
scan.holder$lod <- 0
scan.holder[,4] <- 0

G2.scan <- function(crossfile, scan.object){
		G2sex <- crossfile$pheno$Sex
		AutoNull <- logLik(lm(crossfile $pheno$maxchaetlength ~ 1))
		XNull <- logLik(lm(crossfile $pheno$maxchaetlength ~ G2sex))
		for(j in 1: 10){		#autosomes only
		chr <- names(nmar(crossfile))[j]

				LL.G2chlen <- NULL
					
		for(i in 1:(nmar(crossfile)[chr])){ 	
										
				qtlG2 <- crossfile $geno[[chr]]$data[,i] -1
				LL.G2chlen[i] <- logLik(lm(crossfile $pheno$maxchaetlength ~ qtlG2 ))		
		}					
	
	scan.object[which(scan.object$chr == chr),3] <- LL.G2chlen
	scan.object[which(scan.object$chr == chr),4] <- AutoNull
				
	}
			for(j in 11){		# X only
			chr <- names(nmar(crossfile))[j]
			LL.G2chlen <- NULL
						
			for(i in 1:(nmar(crossfile)[chr])){ 	
										
				qtlG2 <- crossfile $geno[[chr]]$data[,i] -1
				LL.G2chlen[i] <- logLik(lm(crossfile $pheno$maxchaetlength ~ qtlG2*G2sex))		

		}					
	
	scan.object[which(scan.object$chr == chr),3] <- LL.G2chlen
	scan.object[which(scan.object$chr == chr),4] <- XNull
	}

	names(scan.object)[3:4] <- c("LL.G2chlen", "LL.G2Null")
	return(scan.object)
}

ObsG2ChlenScan <- G2.scan(Sb.deduped, scan.holder)

G3.scan <- function(geno.mat, geno.pat, scan.object) {
	dadID <- geno.mat$pheno$Dad_G2_Name
	momID <-  geno.mat$pheno$Mom_G2_Name
	chlen.pheno <- geno.mat$pheno$ChLen
	chlen.G3.null <- logLik(lmer(chlen.pheno ~ 1 + (1|dadID) + (1|momID), REML = F))[1]

		for(j in 1: 10){	# This function is only for the autosomes
		chr <- names(nmar(geno.mat))[j]
		chlen.summed <- NULL

			for(i in 1:(nmar(geno.mat)[chr])){ 							
				qtlAll <- geno.mat$geno[[chr]]$data[,i] + geno.pat$geno[[chr]]$data[,i] - 2 				
				chlen.summed[i] <- logLik(lmer(chlen.pheno ~ qtlAll + (1|dadID) + (1|momID), REML = F))[1]
		}				
	scan.object[which(scan.object$chr == chr),3] <- chlen.summed
	scan.object[which(scan.object$chr == chr),4] <- chlen.G3.null
	}
			for(j in 11){	# X only
		chr <- names(nmar(geno.mat))[j]
		chlen.summed <- NULL

			for(i in 1:(nmar(geno.mat)[chr])){ 													
				qtlMom <- geno.mat$geno[[chr]]$data[,i] -1 # Count of lecithotroph alleles: 0, or 1.
				qtlDad <- geno.pat$geno[[chr]]$data[,i] -1 # Count of lecithotroph alleles: 0, or 1.
				qtlAll <- (1/4)*qtlDad + (3/8)*qtlMom # X-specific G3 additive effect parameterization
				chlen.summed[i] <- logLik(lmer(chlen.pheno ~ qtlAll + (1|dadID) + (1|momID), REML = F))[1]
		}				
	scan.object[which(scan.object$chr == chr),3] <- chlen.summed
	scan.object[which(scan.object$chr == chr),4] <- chlen.G3.null

	}
	names(scan.object)[3:4] <- c("LL.G3chlen", "LL.G3Null")
		
	return(scan.object)
}

ObsG3ChlenScan <- G3.scan(MatGeno,PatGeno, scan.holder)


# Joint Scan
ObsJointChlenScan <- ObsG2ChlenScan + ObsG3ChlenScan
ObsJointChlenScan[,5] <- NA
ObsJointChlenScan[,5] <-  2*(ObsJointChlenScan[,3] - ObsJointChlenScan[,4])
names(ObsJointChlenScan)[c(3,4,5)] <- c("JointChlen", "JointChlenNull", "ChlenChisq")

# Top QTL peak:
ObsJointChlenScan[which(ObsJointChlenScan$ChlenChisq == max(ObsJointChlenScan$ChlenChisq)),]
#       chr      pos JointChlen JointChlenNull ChlenChisq
#SNP497   3 5.441238  -15564.46      -15606.47   84.03385

###############
# A loop to perform the permutations
n.perms = 500
perm.results <- matrix(nrow = n.perms, ncol = 4)
date();
for(m in 1:n.perms){
	
		# Permute the G2 males
	male.perm <- data.frame(levels(factor(Sb.deduped$pheno$Ind[Sb.deduped$pheno$Sex == "M"])), levels(factor(Sb.deduped$pheno$Ind[Sb.deduped$pheno$Sex == "M"])))
	
	# Permutations are stratified by the F1 mother of the F2 fathers, i.e., 1A, 3C, 6F, 8H.
	male.perm[1:12,2] <- sample(male.perm[1:12,1])			
	male.perm[13:27,2] <- sample(male.perm[13:27,1])	
	male.perm[28:36,2] <- sample(male.perm[28:36,1])	
	male.perm[37:58,2] <- sample(male.perm[37:58,1])	

	perm.Sb <- Sb.deduped
	
	for(i in 1:dim(male.perm)[1]){
		for(ch in 1:11){
			perm.Sb$geno[[ch]]$data[i,] <- Sb.deduped$geno[[ch]]$data[which(Sb.deduped$pheno$Ind == male.perm[i,2])   ,]
		}
	}
		
	# Permute the G2 females
	female.perm <-  data.frame(levels(factor(Sb.deduped$pheno$Ind[Sb.deduped$pheno$Sex == "F"])), levels(factor(Sb.deduped$pheno$Ind[Sb.deduped$pheno$Sex == "F"])))

	# Permutations are stratified by the F1 mother of the F2 mothers, i.e., 1A, 3C, 6F, 8H.
	female.perm[1:39,2] <- sample(female.perm[1:39,1])	
	female.perm[40:98,2] <- sample(female.perm[40:98,1])		
	female.perm[99:126,2] <- sample(female.perm[99:126,1])	
	female.perm[127:183,2] <- sample(female.perm[127:183,1])	

	for(i in 1:dim(female.perm)[1]){
		for(ch in 1:11){
			perm.Sb$geno[[ch]]$data[i,] <- Sb.deduped$geno[[ch]]$data[which(Sb.deduped$pheno$Ind == female.perm[i,2])   ,]
		}
	}

	#subset down to the right number for the G3 data
	male.permG3 <- male.perm[which(male.perm[,1] %in% MatGeno$pheno$Dad_G2_Name),]
	female.permG3 <- female.perm[which(female.perm[,1] %in% MatGeno$pheno$Mom_G2_Name),]
	
	perm.Gmat <- MatGeno
	perm.Gpat <- PatGeno
	
	for(i in 1:dim(MatGeno$pheno)[1]){
		permdad <- male.permG3[which(male.permG3[,1] == MatGeno$pheno$Dad_G2_Name[i]),2]
		permmom <- female.permG3[which(female.permG3[,1] == MatGeno$pheno$Mom_G2_Name[i]),2]
		for(j in 1:10){
			perm.Gpat$geno[[j]]$data[i,] <- Sb.deduped$geno[[j]]$data[which(Sb.deduped$pheno$Ind == permdad)[1],]
			perm.Gmat$geno[[j]]$data[i,] <- Sb.deduped$geno[[j]]$data[which(Sb.deduped$pheno$Ind == permmom)[1],]
		}
	}
		
	# perform the joint scan
G2results <- G2.scan(perm.Sb, scan.holder)
G3results <- G3.scan(perm.Gmat, perm.Gpat, scan.holder)
Joint.results <- G2results+G3results
names(Joint.results)[3:4] <- c("Joint.Chlen", "Joint.Chlen.Null") 

	# add the maximum likelihoods to perm.results
perm.results[m,1:2] <- apply(as.matrix(Joint.results[Joint.results $chr != "X",3:4]), 2, max)
perm.results[m,3:4] <- apply(as.matrix(Joint.results[Joint.results $chr == "X",3:4]), 2, max)
perm.out <- data.frame(perm.results)
names(perm.out)  <- c("Joint.Chlen", "Joint.Chlen.Null", "Joint.Chlen.X", "Joint.Chlen.X.Null")
print(m); print(date());
}

rm(perm.results)
perm.out[,5:6] <- NA

perm.out[,5:6] <- cbind(2*(perm.out$Joint.Chlen - perm.out$Joint.Chlen.Null), 2*(perm.out$Joint.Chlen.X - perm.out$Joint.Chlen.X.Null))
names(perm.out)[5:6] <- c("JointAutChisq", "JointXChisq")

apply(perm.out[,c(5,6)], 2, function(x){quantile(x,0.95)})
#JointAutChisq   JointXChisq 
#     16.83911       9.99508 

perm.out -> ChLenJointScan1Perms
save(ChLenJointScan1Perms, file = "ChLenJointScan1.500Perms.R")
rm(perm.out)

##############################################################################
# Second Forward Search
		
	G2chlenQ3.497 <- Sb.deduped$geno[[3]]$data[,"SNP497"] -1
	G3chlenQ3.497 <- MatGeno$geno[[3]]$data[,"SNP497"] + PatGeno$geno[[3]]$data[,"SNP497"] -2

G2.scan <- function(crossfile, scan.object){
		G2sex <- crossfile$pheno$Sex
		AutoNull <- logLik(lm(crossfile $pheno$maxchaetlength ~ G2chlenQ3.497))
		XNull <- logLik(lm(crossfile $pheno$maxchaetlength ~ G2sex + G2chlenQ3.497))
		for(j in 1: 10){		#autosomes only
		chr <- names(nmar(crossfile))[j]

				LL.G2chlen <- NULL
					
		for(i in 1:(nmar(crossfile)[chr])){ 	
										
				qtlG2 <- crossfile $geno[[chr]]$data[,i] -1
				LL.G2chlen[i] <- logLik(lm(crossfile $pheno$maxchaetlength ~ qtlG2 + G2chlenQ3.497))		
		}					
	
	scan.object[which(scan.object$chr == chr),3] <- LL.G2chlen
	scan.object[which(scan.object$chr == chr),4] <- AutoNull
				
	}
			for(j in 11){		# X only
			chr <- names(nmar(crossfile))[j]
			LL.G2chlen <- NULL
						
			for(i in 1:(nmar(crossfile)[chr])){ 	
										
				qtlG2 <- crossfile $geno[[chr]]$data[,i] -1
				LL.G2chlen[i] <- logLik(lm(crossfile $pheno$maxchaetlength ~ qtlG2*G2sex + G2chlenQ3.497))	
		}					
	
	scan.object[which(scan.object$chr == chr),3] <- LL.G2chlen
	scan.object[which(scan.object$chr == chr),4] <- XNull
	}

	names(scan.object)[c(3:4)] <- c("LL.G2chlen", "LL.G2Null")
	return(scan.object)
}

ObsG2ChlenScan <- G2.scan(Sb.deduped, scan.holder)

G3.scan <- function(geno.mat, geno.pat, scan.object) {
	dadID <- geno.mat$pheno$Dad_G2_Name
	momID <-  geno.mat$pheno$Mom_G2_Name
	chlen.pheno <- geno.mat$pheno$ChLen
	chlen.G3.null <- logLik(lmer(chlen.pheno ~ G3chlenQ3.497 + (1|dadID) + (1|momID), REML = F))[1]

		for(j in 1: 10){	# This function is only for the autosomes
		chr <- names(nmar(geno.mat))[j]
		chlen.summed <- NULL

			for(i in 1:(nmar(geno.mat)[chr])){ 							
				qtlAll <- geno.mat$geno[[chr]]$data[,i] + geno.pat$geno[[chr]]$data[,i] - 2 				
				chlen.summed[i] <- logLik(lmer(chlen.pheno ~ qtlAll + G3chlenQ3.497 + (1|dadID) + (1|momID), REML = F))[1]
		}				
	scan.object[which(scan.object$chr == chr),3] <- chlen.summed
	scan.object[which(scan.object$chr == chr),4] <- chlen.G3.null
	}
			for(j in 11){	# X only
		chr <- names(nmar(geno.mat))[j]
		chlen.summed <- NULL

			for(i in 1:(nmar(geno.mat)[chr])){ 													
				qtlMom <- geno.mat$geno[[chr]]$data[,i] -1 # Count of lecithotroph alleles: 0, or 1.
				qtlDad <- geno.pat$geno[[chr]]$data[,i] -1 # Count of lecithotroph alleles: 0, or 1.
				qtlAll <- (1/4)*qtlDad + (3/8)*qtlMom # X-specific G3 additive effect parameterization
				chlen.summed[i] <- logLik(lmer(chlen.pheno ~ qtlAll + G3chlenQ3.497 + (1|dadID) + (1|momID), REML = F))[1]
		}				
	scan.object[which(scan.object$chr == chr),3] <- chlen.summed
	scan.object[which(scan.object$chr == chr),4] <- chlen.G3.null

	}
	names(scan.object)[3:4] <- c("LL.G3chlen", "LL.G3Null")
		
	return(scan.object)
}

ObsG3ChlenScan <- G3.scan(MatGeno,PatGeno, scan.holder)


# Joint Scan
ObsJointChlenScan <- ObsG2ChlenScan + ObsG3ChlenScan
ObsJointChlenScan[,5] <- NA
ObsJointChlenScan[,5] <-  2*(ObsJointChlenScan[,3] - ObsJointChlenScan[,4])
names(ObsJointChlenScan)[c(3,4,5)] <- c("JointChlen", "JointChlenNull", "ChlenChisq")

# Top QTL peak:
ObsJointChlenScan[which(ObsJointChlenScan$ChlenChisq == max(ObsJointChlenScan$ChlenChisq)),]
#       chr      pos JointChlen JointChlenNull ChlenChisq
#SNP272   9 1.457207  -15514.53      -15564.46   99.86374

n.perms = 500
perm.results <- matrix(nrow = n.perms, ncol = 4)
date();
for(m in 1:n.perms){
	
		# Permute the G2 males
	male.perm <- data.frame(levels(factor(Sb.deduped$pheno$Ind[Sb.deduped$pheno$Sex == "M"])), levels(factor(Sb.deduped$pheno$Ind[Sb.deduped$pheno$Sex == "M"])))
	
	# Permutations are stratified by the F1 mother of the F2 fathers, i.e., 1A, 3C, 6F, 8H.
	male.perm[1:12,2] <- sample(male.perm[1:12,1])			
	male.perm[13:27,2] <- sample(male.perm[13:27,1])	
	male.perm[28:36,2] <- sample(male.perm[28:36,1])	
	male.perm[37:58,2] <- sample(male.perm[37:58,1])	

	perm.Sb <- Sb.deduped
	
	for(i in 1:dim(male.perm)[1]){
		for(ch in 1:11){
			perm.Sb$geno[[ch]]$data[i,] <- Sb.deduped$geno[[ch]]$data[which(Sb.deduped$pheno$Ind == male.perm[i,2])   ,]
		}
	}
		
	# Permute the G2 females
	female.perm <-  data.frame(levels(factor(Sb.deduped$pheno$Ind[Sb.deduped$pheno$Sex == "F"])), levels(factor(Sb.deduped$pheno$Ind[Sb.deduped$pheno$Sex == "F"])))

	# Permutations are stratified by the F1 mother of the F2 mothers, i.e., 1A, 3C, 6F, 8H.
	female.perm[1:39,2] <- sample(female.perm[1:39,1])	
	female.perm[40:98,2] <- sample(female.perm[40:98,1])		
	female.perm[99:126,2] <- sample(female.perm[99:126,1])	
	female.perm[127:183,2] <- sample(female.perm[127:183,1])	

	for(i in 1:dim(female.perm)[1]){
		for(ch in 1:11){
			perm.Sb$geno[[ch]]$data[i,] <- Sb.deduped$geno[[ch]]$data[which(Sb.deduped$pheno$Ind == female.perm[i,2])   ,]
		}
	}

	#subset down to the right number for the G3 data
	male.permG3 <- male.perm[which(male.perm[,1] %in% MatGeno$pheno$Dad_G2_Name),]
	female.permG3 <- female.perm[which(female.perm[,1] %in% MatGeno$pheno$Mom_G2_Name),]
	
	perm.Gmat <- MatGeno
	perm.Gpat <- PatGeno
	
	for(i in 1:dim(MatGeno$pheno)[1]){
		permdad <- male.permG3[which(male.permG3[,1] == MatGeno$pheno$Dad_G2_Name[i]),2]
		permmom <- female.permG3[which(female.permG3[,1] == MatGeno$pheno$Mom_G2_Name[i]),2]
		for(j in 1:10){
			perm.Gpat$geno[[j]]$data[i,] <- Sb.deduped$geno[[j]]$data[which(Sb.deduped$pheno$Ind == permdad)[1],]
			perm.Gmat$geno[[j]]$data[i,] <- Sb.deduped$geno[[j]]$data[which(Sb.deduped$pheno$Ind == permmom)[1],]
		}
	}
		
	# perform the joint scan
G2results <- G2.scan(perm.Sb, scan.holder)
G3results <- G3.scan(perm.Gmat, perm.Gpat, scan.holder)
Joint.results <- G2results+G3results
names(Joint.results)[3:4] <- c("Joint.Chlen", "Joint.Chlen.Null") 

	# add the maximum likelihoods to perm.results
perm.results[m,1:2] <- apply(as.matrix(Joint.results[Joint.results $chr != "X",3:4]), 2, max)
perm.results[m,3:4] <- apply(as.matrix(Joint.results[Joint.results $chr == "X",3:4]), 2, max)
perm.out <- data.frame(perm.results)
names(perm.out)  <- c("Joint.Chlen", "Joint.Chlen.Null", "Joint.Chlen.X", "Joint.Chlen.X.Null")
print(m); print(date());
}

rm(perm.results)
perm.out[,5:6] <- NA

perm.out[,5:6] <- cbind(2*(perm.out$Joint.Chlen - perm.out$Joint.Chlen.Null), 2*(perm.out$Joint.Chlen.X - perm.out$Joint.Chlen.X.Null))
names(perm.out)[5:6] <- c("JointAutChisq", "JointXChisq")

apply(perm.out[,c(5,6)], 2, function(x){quantile(x,0.95)})
#JointAutChisq   JointXChisq 
#  18.06530      10.53385 


perm.out -> ChLenJointScan2Perms
save(ChLenJointScan2Perms, file = "ChLenJointScan2.500Perms.R")
rm(perm.out)


##############################################################################
# Third Forward Search
		
	G2chlenQ3.497 <- Sb.deduped$geno[[3]]$data[,"SNP497"] -1
	G3chlenQ3.497 <- MatGeno$geno[[3]]$data[,"SNP497"] + PatGeno$geno[[3]]$data[,"SNP497"] -2

	G2chlenQ9.272 <- Sb.deduped$geno[[9]]$data[,"SNP272"] -1
	G3chlenQ9.272 <- MatGeno$geno[[9]]$data[,"SNP272"] + PatGeno$geno[[9]]$data[,"SNP272"] -2

G2.scan <- function(crossfile, scan.object){
		G2sex <- crossfile$pheno$Sex
		AutoNull <- logLik(lm(crossfile $pheno$maxchaetlength ~ G2chlenQ3.497 + G2chlenQ9.272))
		XNull <- logLik(lm(crossfile $pheno$maxchaetlength ~ G2sex + G2chlenQ3.497 + G2chlenQ9.272))
		for(j in 1: 10){		#autosomes only
		chr <- names(nmar(crossfile))[j]

				LL.G2chlen <- NULL
					
		for(i in 1:(nmar(crossfile)[chr])){ 	
										
				qtlG2 <- crossfile $geno[[chr]]$data[,i] -1
				LL.G2chlen[i] <- logLik(lm(crossfile $pheno$maxchaetlength ~ qtlG2 + G2chlenQ3.497 + G2chlenQ9.272))		
		}					
	
	scan.object[which(scan.object$chr == chr),3] <- LL.G2chlen
	scan.object[which(scan.object$chr == chr),4] <- AutoNull
				
	}
			for(j in 11){		# X only
			chr <- names(nmar(crossfile))[j]
			LL.G2chlen <- NULL
						
			for(i in 1:(nmar(crossfile)[chr])){ 	
										
				qtlG2 <- crossfile $geno[[chr]]$data[,i] -1
				LL.G2chlen[i] <- logLik(lm(crossfile $pheno$maxchaetlength ~ qtlG2*G2sex + G2chlenQ3.497 + G2chlenQ9.272))	
		}					
	
	scan.object[which(scan.object$chr == chr),3] <- LL.G2chlen
	scan.object[which(scan.object$chr == chr),4] <- XNull
	}

	names(scan.object)[c(3:4)] <- c("LL.G2chlen", "LL.G2Null")
	return(scan.object)
}

ObsG2ChlenScan <- G2.scan(Sb.deduped, scan.holder)

G3.scan <- function(geno.mat, geno.pat, scan.object) {
	dadID <- geno.mat$pheno$Dad_G2_Name
	momID <-  geno.mat$pheno$Mom_G2_Name
	chlen.pheno <- geno.mat$pheno$ChLen
	chlen.G3.null <- logLik(lmer(chlen.pheno ~ G3chlenQ3.497 + G3chlenQ9.272 + (1|dadID) + (1|momID), REML = F))[1]

		for(j in 1: 10){	# This function is only for the autosomes
		chr <- names(nmar(geno.mat))[j]
		chlen.summed <- NULL

			for(i in 1:(nmar(geno.mat)[chr])){ 							
				qtlAll <- geno.mat$geno[[chr]]$data[,i] + geno.pat$geno[[chr]]$data[,i] - 2 				
				chlen.summed[i] <- logLik(lmer(chlen.pheno ~ qtlAll + G3chlenQ3.497 + G3chlenQ9.272 + (1|dadID) + (1|momID), REML = F))[1]
		}				
	scan.object[which(scan.object$chr == chr),3] <- chlen.summed
	scan.object[which(scan.object$chr == chr),4] <- chlen.G3.null
	}
			for(j in 11){	# X only
		chr <- names(nmar(geno.mat))[j]
		chlen.summed <- NULL

			for(i in 1:(nmar(geno.mat)[chr])){ 													
				qtlMom <- geno.mat$geno[[chr]]$data[,i] -1 # Count of lecithotroph alleles: 0, or 1.
				qtlDad <- geno.pat$geno[[chr]]$data[,i] -1 # Count of lecithotroph alleles: 0, or 1.
				qtlAll <- (1/4)*qtlDad + (3/8)*qtlMom # X-specific G3 additive effect parameterization
				chlen.summed[i] <- logLik(lmer(chlen.pheno ~ qtlAll + G3chlenQ3.497 + G3chlenQ9.272 + (1|dadID) + (1|momID), REML = F))[1]
		}				
	scan.object[which(scan.object$chr == chr),3] <- chlen.summed
	scan.object[which(scan.object$chr == chr),4] <- chlen.G3.null

	}
	names(scan.object)[3:4] <- c("LL.G3chlen", "LL.G3Null")
		
	return(scan.object)
}

ObsG3ChlenScan <- G3.scan(MatGeno,PatGeno, scan.holder)


# Joint Scan
ObsJointChlenScan <- ObsG2ChlenScan + ObsG3ChlenScan
ObsJointChlenScan[,5] <- NA
ObsJointChlenScan[,5] <-  2*(ObsJointChlenScan[,3] - ObsJointChlenScan[,4])
names(ObsJointChlenScan)[c(3,4,5)] <- c("JointChlen", "JointChlenNull", "ChlenChisq")


# Top QTL peak:
ObsJointChlenScan[which(ObsJointChlenScan$ChlenChisq == max(ObsJointChlenScan$ChlenChisq)),]
#        chr      pos JointChlen JointChlenNull ChlenChisq
#SNP1262   5 9.347682  -15507.96      -15514.53   13.12713


n.perms = 500
perm.results <- matrix(nrow = n.perms, ncol = 4)
date();
for(m in 1:n.perms){
	
		# Permute the G2 males
	male.perm <- data.frame(levels(factor(Sb.deduped$pheno$Ind[Sb.deduped$pheno$Sex == "M"])), levels(factor(Sb.deduped$pheno$Ind[Sb.deduped$pheno$Sex == "M"])))
	
	# Permutations are stratified by the F1 mother of the F2 fathers, i.e., 1A, 3C, 6F, 8H.
	male.perm[1:12,2] <- sample(male.perm[1:12,1])			
	male.perm[13:27,2] <- sample(male.perm[13:27,1])	
	male.perm[28:36,2] <- sample(male.perm[28:36,1])	
	male.perm[37:58,2] <- sample(male.perm[37:58,1])	

	perm.Sb <- Sb.deduped
	
	for(i in 1:dim(male.perm)[1]){
		for(ch in 1:11){
			perm.Sb$geno[[ch]]$data[i,] <- Sb.deduped$geno[[ch]]$data[which(Sb.deduped$pheno$Ind == male.perm[i,2])   ,]
		}
	}
		
	# Permute the G2 females
	female.perm <-  data.frame(levels(factor(Sb.deduped$pheno$Ind[Sb.deduped$pheno$Sex == "F"])), levels(factor(Sb.deduped$pheno$Ind[Sb.deduped$pheno$Sex == "F"])))

	# Permutations are stratified by the F1 mother of the F2 mothers, i.e., 1A, 3C, 6F, 8H.
	female.perm[1:39,2] <- sample(female.perm[1:39,1])	
	female.perm[40:98,2] <- sample(female.perm[40:98,1])		
	female.perm[99:126,2] <- sample(female.perm[99:126,1])	
	female.perm[127:183,2] <- sample(female.perm[127:183,1])	

	for(i in 1:dim(female.perm)[1]){
		for(ch in 1:11){
			perm.Sb$geno[[ch]]$data[i,] <- Sb.deduped$geno[[ch]]$data[which(Sb.deduped$pheno$Ind == female.perm[i,2])   ,]
		}
	}

	#subset down to the right number for the G3 data
	male.permG3 <- male.perm[which(male.perm[,1] %in% MatGeno$pheno$Dad_G2_Name),]
	female.permG3 <- female.perm[which(female.perm[,1] %in% MatGeno$pheno$Mom_G2_Name),]
	
	perm.Gmat <- MatGeno
	perm.Gpat <- PatGeno
	
	for(i in 1:dim(MatGeno$pheno)[1]){
		permdad <- male.permG3[which(male.permG3[,1] == MatGeno$pheno$Dad_G2_Name[i]),2]
		permmom <- female.permG3[which(female.permG3[,1] == MatGeno$pheno$Mom_G2_Name[i]),2]
		for(j in 1:10){
			perm.Gpat$geno[[j]]$data[i,] <- Sb.deduped$geno[[j]]$data[which(Sb.deduped$pheno$Ind == permdad)[1],]
			perm.Gmat$geno[[j]]$data[i,] <- Sb.deduped$geno[[j]]$data[which(Sb.deduped$pheno$Ind == permmom)[1],]
		}
	}
		
	# perform the joint scan
G2results <- G2.scan(perm.Sb, scan.holder)
G3results <- G3.scan(perm.Gmat, perm.Gpat, scan.holder)
Joint.results <- G2results+G3results
names(Joint.results)[3:4] <- c("Joint.Chlen", "Joint.Chlen.Null") 

	# add the maximum likelihoods to perm.results
perm.results[m,1:2] <- apply(as.matrix(Joint.results[Joint.results $chr != "X",3:4]), 2, max)
perm.results[m,3:4] <- apply(as.matrix(Joint.results[Joint.results $chr == "X",3:4]), 2, max)
perm.out <- data.frame(perm.results)
names(perm.out)  <- c("Joint.Chlen", "Joint.Chlen.Null", "Joint.Chlen.X", "Joint.Chlen.X.Null")
print(m); print(date());
}

rm(perm.results)
perm.out[,5:6] <- NA

perm.out[,5:6] <- cbind(2*(perm.out$Joint.Chlen - perm.out$Joint.Chlen.Null), 2*(perm.out$Joint.Chlen.X - perm.out$Joint.Chlen.X.Null))
names(perm.out)[5:6] <- c("JointAutChisq", "JointXChisq")

apply(perm.out[,c(5,6)], 2, function(x){quantile(x,0.95)})
#JointAutChisq   JointXChisq 
#     16.12797      10.35372 

perm.out -> ChLenJointScan3Perms
save(ChLenJointScan3Perms, file = "ChLenJointScan3.500Perms.R")
rm(perm.out)

# No locus exceeds the p=0.05 threshold. 
```
