## Supplementary File 1. Data and Code for "Baby makes three: maternal, paternal, and zygotic genetic effects shape larval phenotypic evolution": JointChNumScan.R

```
# Combine likelihoods from G2s and G3s

library(qtl)
library(lme4)

load(file = "G3AnalysisObjects.R")
# [1] "dups"       "MatGeno"    "PatGeno"    "Sb.deduped"


scan.holder <- scanone(Sb.deduped, pheno = "nchaetae")
scan.holder$lod <- 0
scan.holder[,4:6] <- 0

# Redefine the phenotypes as a binary trait and a Box-Cox-adjusted continuous trait
# First for the G2s
G2chnum.bin.pheno <- Sb.deduped$pheno$nchaetae
G2chnum.bin.pheno[G2chnum.bin.pheno > 0] <- 1
G2bc.chnum <- ((Sb.deduped$pheno$nchaetae ^0.18)-1)/0.18 # uses the maximum likelihood Box-Cox coefficient for these data
G2bc.chnum[Sb.deduped$pheno$nchaetae == 0] <- NA
Sb.deduped$pheno[,9] <- G2chnum.bin.pheno
Sb.deduped$pheno[,10] <- G2bc.chnum
names(Sb.deduped$pheno)[c(9,10)] <- c("G2.chnum.bin.pheno", "G2bc.chnum")

# Then for the G3s
chnum.bin.pheno <- MatGeno$pheno$nChaetae
chnum.bin.pheno[chnum.bin.pheno > 0] <- 1
bc.chnum0 <- ((MatGeno$pheno$nChaetae^0.29)-1)/0.29 # uses the maximum likelihood Box-Cox coefficient for these data
bc.chnum0[MatGeno$pheno$nChaetae == 0] <- NA
last.column <- dim(MatGeno$pheno)[2]
MatGeno$pheno[,(last.column + 1)] <- chnum.bin.pheno
MatGeno$pheno[,(last.column + 2)] <- bc.chnum0
names(MatGeno$pheno)[c((last.column +1) : (last.column +2))] <- c("chnum.bin.pheno","bc.chnum0")

###
# First round forward search

G2.scan <- function(crossfile, scan.object){
		G2sex <- crossfile$pheno$Sex
		binpheno <- crossfile$pheno$G2.chnum.bin.pheno
		bcpheno <- crossfile$pheno$G2bc.chnum
		binAutoNull <- logLik(glm(binpheno ~ 1, family = "binomial"))
		binXNull <- logLik(glm(binpheno ~ G2sex, family = "binomial"))
		bcAutoNull <- logLik(lm(bcpheno ~ 1))	
		bcXNull <- logLik(lm(bcpheno ~ G2sex))	
				
		for(j in 1: 10){		#autosomes only
		chr <- names(nmar(crossfile))[j]

				LL.G2chnumbin <- NULL
				LL.G2bcchnum <- NULL	
					
		for(i in 1:(nmar(crossfile)[chr])){ 	
										
				qtlG2 <- crossfile$geno[[chr]]$data[,i] -1
				LL.G2chnumbin[i] <- logLik(glm(binpheno ~ qtlG2, family = "binomial" ))		
				LL.G2bcchnum[i] <- logLik(lm(bcpheno ~ qtlG2))	
					}					
	
	scan.object[which(scan.object$chr == chr),3] <- LL.G2chnumbin
	scan.object[which(scan.object$chr == chr),4] <- binAutoNull
	scan.object[which(scan.object$chr == chr),5] <- LL.G2bcchnum
	scan.object[which(scan.object$chr == chr),6] <- bcAutoNull
	
				
	}
			for(j in 11){		# X only
			chr <- names(nmar(crossfile))[j]
				LL.G2chnumbin <- NULL
				LL.G2bcchnum <- NULL	
						
			for(i in 1:(nmar(crossfile)[chr])){ 	
										
				qtlG2 <- crossfile$geno[[chr]]$data[,i] -1
				LL.G2chnumbin[i] <- logLik(glm(binpheno ~ qtlG2*G2sex, family = "binomial" ))		
				LL.G2bcchnum[i] <- logLik(lm(bcpheno ~ qtlG2*G2sex ))	

		}					
	
	scan.object[which(scan.object$chr == chr),3] <- LL.G2chnumbin
	scan.object[which(scan.object$chr == chr),4] <- binXNull
	scan.object[which(scan.object$chr == chr),5] <- LL.G2bcchnum
	scan.object[which(scan.object$chr == chr),6] <- bcXNull
	}

	names(scan.object)[c(3:6)] <- c("LL.G2ChNumBin", "LL.G2ChNumBinNull", "LL.G2ChNum", "LL.G2ChNumNull")
	return(scan.object)
}

ObsG2ChNumScan <- G2.scan(Sb.deduped, scan.holder)

G3.scan <- function(geno.mat, geno.pat, scan.object) {
	dadID <- geno.mat$pheno$Dad_G2_Name
	momID <-  geno.mat$pheno$Mom_G2_Name
		
	binpheno <- geno.mat$pheno$chnum.bin.pheno
	bcpheno <- geno.mat$pheno$bc.chnum0
	bin.null <- logLik(glmer(binpheno ~ 1 + (1|dadID) + (1|momID), family = "binomial"))[1]
	bc.null <- 	logLik(lmer(bcpheno ~ 1 + (1|dadID) + (1|momID), REML = F))[1]
	
		for(j in 1: 10){	# This function is only for the autosomes
		chr <- names(nmar(geno.mat))[j]
		bin.summed <- NULL
		bc.summed <- NULL
			for(i in 1:(nmar(geno.mat)[chr])){ 							
				qtlAll <- geno.mat$geno[[chr]]$data[,i] + geno.pat$geno[[chr]]$data[,i] - 2 				

				bin.summed[i] <- logLik(glmer(binpheno ~ qtlAll + (1|dadID) + (1|momID), family = "binomial"))[1]
				bc.summed[i] <- logLik(lmer(bcpheno ~ qtlAll + (1|dadID) + (1|momID), REML = F))[1]


		}				
	scan.object[which(scan.object$chr == chr),3] <- bin.summed
	scan.object[which(scan.object$chr == chr),4] <- bin.null
	scan.object[which(scan.object$chr == chr),5] <- bc.summed
	scan.object[which(scan.object$chr == chr),6] <- bc.null
	}
			for(j in 11){	# X only
		chr <- names(nmar(geno.mat))[j]
		bin.summed <- NULL
		bc.summed <- NULL

			for(i in 1:(nmar(geno.mat)[chr])){ 													
				qtlMom <- geno.mat$geno[[chr]]$data[,i] -1 # Count of lecithotroph alleles: 0, or 1.
				qtlDad <- geno.pat$geno[[chr]]$data[,i] -1 # Count of lecithotroph alleles: 0, or 1.
				qtlAll <- (1/4)*qtlDad + (3/8)*qtlMom # X-specific G3 additive effect parameterization
				bin.summed[i] <- logLik(glmer(binpheno ~ qtlAll + (1|dadID) + (1|momID), family = "binomial"))[1]
				bc.summed[i] <- logLik(lmer(bcpheno ~ qtlAll + (1|dadID) + (1|momID), REML = F))[1]

		}				
	scan.object[which(scan.object$chr == chr),3] <- bin.summed
	scan.object[which(scan.object$chr == chr),4] <- bin.null
	scan.object[which(scan.object$chr == chr),5] <- bc.summed
	scan.object[which(scan.object$chr == chr),6] <- bc.null
	}
	names(scan.object)[c(3:6)] <- c("LL.G3Bin", "LL.G3BinNull", "LL.G3ChNum", "LL.G3ChNumNull")
		
	return(scan.object)
}

ObsG3ChNumScan <- G3.scan(MatGeno,PatGeno, scan.holder)


# Joint Scan
ObsJointChNumScan <- ObsG2ChNumScan + ObsG3ChNumScan
names(ObsJointChNumScan)[c(3:6)] <- c("JointBin", "JointBinNull", "JointBC", "JointBCNull")
ObsJointChNumScan[,7:11] <- NA
ObsJointChNumScan[,c(7,8)] <-  c(ObsJointChNumScan$JointBin + ObsJointChNumScan$JointBC, ObsJointChNumScan$JointBinNull + ObsJointChNumScan$JointBCNull)
ObsJointChNumScan[,c(9:11)] <- c(2*(ObsJointChNumScan[,3] - ObsJointChNumScan[,4]), 2*(ObsJointChNumScan[,5] - ObsJointChNumScan[,6]), 2*(ObsJointChNumScan[,7] - ObsJointChNumScan[,8]))
names(ObsJointChNumScan)[c(7:11)] <- c("JointCombo", "JointComboNull", "BinChisq", "BCChisq", "ComboChisq")


# Top QTL peak:
ObsJointChNumScan[which(ObsJointChNumScan $ComboChisq == max(ObsJointChNumScan $ComboChisq)),]
#       chr      pos  JointBin JointBinNull   JointBC JointBCNull JointCombo JointComboNull
#SNP231   8 1.663753 -1937.477    -1966.547 -4425.415   -4454.326  -6362.892      -6420.873
#       BinChisq  BCChisq ComboChisq
#SNP231 58.13829 57.82302   115.9613


###############
# A loop to perform the permutations
n.perms = 100
perm.results <- matrix(nrow = n.perms, ncol = 12)
date();
for(m in 1:n.perms){
	
		# Permute the G2 males
	male.perm <- data.frame(levels(factor(Sb.deduped$pheno$Ind[Sb.deduped$pheno$Sex == "M"])), levels(factor(Sb.deduped$pheno$Ind[Sb.deduped$pheno$Sex == "M"])))
	
	# Permutations are stratified by the F1 mother of the F2 fathers, i.e., 1A, 3C, 6F, 8H.
	male.perm[1:12,2] <- sample(male.perm[1:12,1])			
	male.perm[13:27,2] <- sample(male.perm[13:27,1])	
	male.perm[28:36,2] <- sample(male.perm[28:36,1])	
	male.perm[37:58,2] <- sample(male.perm[37:58,1])	

	perm.Sb <- Sb.deduped
	
	for(i in 1:dim(male.perm)[1]){
		for(ch in 1:11){
			perm.Sb$geno[[ch]]$data[i,] <- Sb.deduped$geno[[ch]]$data[which(Sb.deduped$pheno$Ind == male.perm[i,2])   ,]
		}
	}
		
	# Permute the G2 females
	female.perm <-  data.frame(levels(factor(Sb.deduped$pheno$Ind[Sb.deduped$pheno$Sex == "F"])), levels(factor(Sb.deduped$pheno$Ind[Sb.deduped$pheno$Sex == "F"])))

	# Permutations are stratified by the F1 mother of the F2 mothers, i.e., 1A, 3C, 6F, 8H.
	female.perm[1:39,2] <- sample(female.perm[1:39,1])	
	female.perm[40:98,2] <- sample(female.perm[40:98,1])		
	female.perm[99:126,2] <- sample(female.perm[99:126,1])	
	female.perm[127:183,2] <- sample(female.perm[127:183,1])	

	for(i in 1:dim(female.perm)[1]){
		for(ch in 1:11){
			perm.Sb$geno[[ch]]$data[i,] <- Sb.deduped$geno[[ch]]$data[which(Sb.deduped$pheno$Ind == female.perm[i,2])   ,]
		}
	}

	#subset down to the right number for the G3 data
	male.permG3 <- male.perm[which(male.perm[,1] %in% MatGeno$pheno$Dad_G2_Name),]
	female.permG3 <- female.perm[which(female.perm[,1] %in% MatGeno$pheno$Mom_G2_Name),]
	
	perm.Gmat <- MatGeno
	perm.Gpat <- PatGeno
	
	for(i in 1:dim(MatGeno$pheno)[1]){
		permdad <- male.permG3[which(male.permG3[,1] == MatGeno$pheno$Dad_G2_Name[i]),2]
		permmom <- female.permG3[which(female.permG3[,1] == MatGeno$pheno$Mom_G2_Name[i]),2]
		for(j in 1:10){
			perm.Gpat$geno[[j]]$data[i,] <- Sb.deduped$geno[[j]]$data[which(Sb.deduped$pheno$Ind == permdad)[1],]
			perm.Gmat$geno[[j]]$data[i,] <- Sb.deduped$geno[[j]]$data[which(Sb.deduped$pheno$Ind == permmom)[1],]
		}
	}
		
	# perform the joint scan
	G2results <- G2.scan(perm.Sb, scan.holder)
	G3results <- G3.scan(perm.Gmat, perm.Gpat, scan.holder)
	Joint.results <- G2results+G3results
	names(Joint.results)[3:6] <- c("JointBin", "JointBinNull", "JointBC", "JointBCNull")
	Joint.results[,7:8] <- NA
	Joint.results[,c(7,8)] <-  c(Joint.results$JointBin + Joint.results$JointBC, Joint.results$JointBinNull + Joint.results$JointBCNull)
	
		# add the maximum likelihoods to perm.results
	perm.results[m,1:6] <- apply(as.matrix(Joint.results[Joint.results $chr != "X",3:8]), 2, max)
	perm.results[m,7:12] <- apply(as.matrix(Joint.results[Joint.results $chr == "X",3:8]), 2, max)
	perm.out <- data.frame(perm.results)
	names(perm.out)  <- c("JointBin", "JointBinNull", "JointBC", "JointBCNull", "JointCombo","JointComboNull", "JointBinX", "JointBinNullX", "JointBCX", "JointBCNullX", "JointComboX","JointComboNullX")
	print(m); print(date());
}

rm(perm.results)
perm.out[,13:18] <- NA

perm.out[,13:18] <- cbind(2*(perm.out$JointBin - perm.out$JointBinNull), 2*(perm.out$JointBinX - perm.out$JointBinNullX), 2*(perm.out$JointBC - perm.out$JointBCNull), 2*(perm.out$JointBCX - perm.out$JointBCNullX), 2*(perm.out$JointCombo - perm.out$JointComboNull), 2*(perm.out$JointComboX - perm.out$JointComboNullX))
names(perm.out)[13:18] <- c("BinAutChisq", "BinXChisq", "BCAutChisq", "BCXChisq", "ComboAutChisq","ComboXChisq")

apply(perm.out[,c(13:18)], 2, function(x){quantile(x,0.95)})
#  BinAutChisq     BinXChisq    BCAutChisq      BCXChisq ComboAutChisq   ComboXChisq 
#    14.197261     12.201311     14.459103      9.516525     20.623615     16.967095 


perm.out -> ChNumJointScan1Perms
save(ChNumJointScan1Perms, file = "ChNumJointScan1.Perms.R")
rm(perm.out)

##############################################################################
# Second Forward Search
# The G3 scans for the logistic part of the model do not include random effect of G2 father to avoid singular fit		
		
	G2chlenQ8.231 <- Sb.deduped$geno[[8]]$data[,"SNP231"] -1
	G3chlenQ8.231 <- MatGeno$geno[[8]]$data[,"SNP231"] + PatGeno$geno[[8]]$data[,"SNP231"] -2

G2.scan <- function(crossfile, scan.object){
		G2sex <- crossfile$pheno$Sex
		binpheno <- crossfile$pheno$G2.chnum.bin.pheno
		bcpheno <- crossfile$pheno$G2bc.chnum
		binAutoNull <- logLik(glm(binpheno ~ G2chlenQ8.231, family = "binomial"))
		binXNull <- logLik(glm(binpheno ~ G2sex + G2chlenQ8.231, family = "binomial"))
		bcAutoNull <- logLik(lm(bcpheno ~ G2chlenQ8.231))	
		bcXNull <- logLik(lm(bcpheno ~ G2sex + G2chlenQ8.231))	
				
		for(j in 1: 10){		#autosomes only
		chr <- names(nmar(crossfile))[j]

				LL.G2chnumbin <- NULL
				LL.G2bcchnum <- NULL	
					
		for(i in 1:(nmar(crossfile)[chr])){ 	
										
				qtlG2 <- crossfile$geno[[chr]]$data[,i] -1
				LL.G2chnumbin[i] <- logLik(glm(binpheno ~ qtlG2 + G2chlenQ8.231, family = "binomial" ))		
				LL.G2bcchnum[i] <- logLik(lm(bcpheno ~ qtlG2 + G2chlenQ8.231))	
					}					
	
	scan.object[which(scan.object$chr == chr),3] <- LL.G2chnumbin
	scan.object[which(scan.object$chr == chr),4] <- binAutoNull
	scan.object[which(scan.object$chr == chr),5] <- LL.G2bcchnum
	scan.object[which(scan.object$chr == chr),6] <- bcAutoNull
	
				
	}
			for(j in 11){		# X only
			chr <- names(nmar(crossfile))[j]
				LL.G2chnumbin <- NULL
				LL.G2bcchnum <- NULL	
						
			for(i in 1:(nmar(crossfile)[chr])){ 	
										
				qtlG2 <- crossfile$geno[[chr]]$data[,i] -1
				LL.G2chnumbin[i] <- logLik(glm(binpheno ~ qtlG2*G2sex + G2chlenQ8.231, family = "binomial" ))		
				LL.G2bcchnum[i] <- logLik(lm(bcpheno ~ qtlG2*G2sex + G2chlenQ8.231 ))	

		}					
	
	scan.object[which(scan.object$chr == chr),3] <- LL.G2chnumbin
	scan.object[which(scan.object$chr == chr),4] <- binXNull
	scan.object[which(scan.object$chr == chr),5] <- LL.G2bcchnum
	scan.object[which(scan.object$chr == chr),6] <- bcXNull
	}

	names(scan.object)[c(3:6)] <- c("LL.G2ChNumBin", "LL.G2ChNumBinNull", "LL.G2ChNum", "LL.G2ChNumNull")
	return(scan.object)
}

ObsG2ChNumScan <- G2.scan(Sb.deduped, scan.holder)

G3.scan <- function(geno.mat, geno.pat, scan.object) {
	dadID <- geno.mat$pheno$Dad_G2_Name
	momID <-  geno.mat$pheno$Mom_G2_Name
		
	binpheno <- geno.mat$pheno$chnum.bin.pheno
	bcpheno <- geno.mat$pheno$bc.chnum0
	bin.null <- logLik(glmer(binpheno ~ G3chlenQ8.231 + (1|momID), family = "binomial"))[1]
	bc.null <- 	logLik(lmer(bcpheno ~ G3chlenQ8.231 + (1|dadID) + (1|momID), REML = F))[1]
	
		for(j in 1: 10){	# This function is only for the autosomes
		chr <- names(nmar(geno.mat))[j]
		bin.summed <- NULL
		bc.summed <- NULL
			for(i in 1:(nmar(geno.mat)[chr])){ 							
				qtlAll <- geno.mat$geno[[chr]]$data[,i] + geno.pat$geno[[chr]]$data[,i] - 2 				

				bin.summed[i] <- logLik(glmer(binpheno ~ qtlAll + G3chlenQ8.231 + (1|momID), family = "binomial"))[1]
				bc.summed[i] <- logLik(lmer(bcpheno ~ qtlAll + G3chlenQ8.231 + (1|dadID) + (1|momID), REML = F))[1]


		}				
	scan.object[which(scan.object$chr == chr),3] <- bin.summed
	scan.object[which(scan.object$chr == chr),4] <- bin.null
	scan.object[which(scan.object$chr == chr),5] <- bc.summed
	scan.object[which(scan.object$chr == chr),6] <- bc.null
	}
			for(j in 11){	# X only
		chr <- names(nmar(geno.mat))[j]
		bin.summed <- NULL
		bc.summed <- NULL

			for(i in 1:(nmar(geno.mat)[chr])){ 													
				qtlMom <- geno.mat$geno[[chr]]$data[,i] -1 # Count of lecithotroph alleles: 0, or 1.
				qtlDad <- geno.pat$geno[[chr]]$data[,i] -1 # Count of lecithotroph alleles: 0, or 1.
				qtlAll <- (1/4)*qtlDad + (3/8)*qtlMom # X-specific G3 additive effect parameterization
				bin.summed[i] <- logLik(glmer(binpheno ~ qtlAll + G3chlenQ8.231 + (1|momID), family = "binomial"))[1]
				bc.summed[i] <- logLik(lmer(bcpheno ~ qtlAll + G3chlenQ8.231 + (1|dadID) + (1|momID), REML = F))[1]

		}				
	scan.object[which(scan.object$chr == chr),3] <- bin.summed
	scan.object[which(scan.object$chr == chr),4] <- bin.null
	scan.object[which(scan.object$chr == chr),5] <- bc.summed
	scan.object[which(scan.object$chr == chr),6] <- bc.null
	}
	names(scan.object)[c(3:6)] <- c("LL.G3Bin", "LL.G3BinNull", "LL.G3ChNum", "LL.G3ChNumNull")
		
	return(scan.object)
}

ObsG3ChNumScan <- G3.scan(MatGeno,PatGeno, scan.holder)


# Joint Scan
ObsJointChNumScan <- ObsG2ChNumScan + ObsG3ChNumScan
names(ObsJointChNumScan)[c(3:6)] <- c("JointBin", "JointBinNull", "JointBC", "JointBCNull")
ObsJointChNumScan[,7:11] <- NA
ObsJointChNumScan[,c(7,8)] <-  c(ObsJointChNumScan$JointBin + ObsJointChNumScan$JointBC, ObsJointChNumScan$JointBinNull + ObsJointChNumScan$JointBCNull)
ObsJointChNumScan[,c(9:11)] <- c(2*(ObsJointChNumScan[,3] - ObsJointChNumScan[,4]), 2*(ObsJointChNumScan[,5] - ObsJointChNumScan[,6]), 2*(ObsJointChNumScan[,7] - ObsJointChNumScan[,8]))
names(ObsJointChNumScan)[c(7:11)] <- c("JointCombo", "JointComboNull", "BinChisq", "BCChisq", "ComboChisq")

# Top QTL peak:
ObsJointChNumScan[which(ObsJointChNumScan $ComboChisq == max(ObsJointChNumScan $ComboChisq)),]
#       chr      pos  JointBin JointBinNull  JointBC JointBCNull JointCombo JointComboNull
#SNP497   3 5.441238 -1910.682    -1937.481 -4385.14   -4425.415  -6295.822      -6362.896
#       BinChisq  BCChisq ComboChisq
#SNP497 53.59675 80.54993   134.1467


############
# A loop to perform the permutations
n.perms = 100
perm.results <- matrix(nrow = n.perms, ncol = 12)
date();
for(m in 1:n.perms){
	
		# Permute the G2 males
	male.perm <- data.frame(levels(factor(Sb.deduped$pheno$Ind[Sb.deduped$pheno$Sex == "M"])), levels(factor(Sb.deduped$pheno$Ind[Sb.deduped$pheno$Sex == "M"])))
	
	# Permutations are stratified by the F1 mother of the F2 fathers, i.e., 1A, 3C, 6F, 8H.
	male.perm[1:12,2] <- sample(male.perm[1:12,1])			
	male.perm[13:27,2] <- sample(male.perm[13:27,1])	
	male.perm[28:36,2] <- sample(male.perm[28:36,1])	
	male.perm[37:58,2] <- sample(male.perm[37:58,1])	

	perm.Sb <- Sb.deduped
	
	for(i in 1:dim(male.perm)[1]){
		for(ch in 1:11){
			perm.Sb$geno[[ch]]$data[i,] <- Sb.deduped$geno[[ch]]$data[which(Sb.deduped$pheno$Ind == male.perm[i,2])   ,]
		}
	}
		
	# Permute the G2 females
	female.perm <-  data.frame(levels(factor(Sb.deduped$pheno$Ind[Sb.deduped$pheno$Sex == "F"])), levels(factor(Sb.deduped$pheno$Ind[Sb.deduped$pheno$Sex == "F"])))

	# Permutations are stratified by the F1 mother of the F2 mothers, i.e., 1A, 3C, 6F, 8H.
	female.perm[1:39,2] <- sample(female.perm[1:39,1])	
	female.perm[40:98,2] <- sample(female.perm[40:98,1])		
	female.perm[99:126,2] <- sample(female.perm[99:126,1])	
	female.perm[127:183,2] <- sample(female.perm[127:183,1])	

	for(i in 1:dim(female.perm)[1]){
		for(ch in 1:11){
			perm.Sb$geno[[ch]]$data[i,] <- Sb.deduped$geno[[ch]]$data[which(Sb.deduped$pheno$Ind == female.perm[i,2])   ,]
		}
	}

	#subset down to the right number for the G3 data
	male.permG3 <- male.perm[which(male.perm[,1] %in% MatGeno$pheno$Dad_G2_Name),]
	female.permG3 <- female.perm[which(female.perm[,1] %in% MatGeno$pheno$Mom_G2_Name),]
	
	perm.Gmat <- MatGeno
	perm.Gpat <- PatGeno
	
	for(i in 1:dim(MatGeno$pheno)[1]){
		permdad <- male.permG3[which(male.permG3[,1] == MatGeno$pheno$Dad_G2_Name[i]),2]
		permmom <- female.permG3[which(female.permG3[,1] == MatGeno$pheno$Mom_G2_Name[i]),2]
		for(j in 1:10){
			perm.Gpat$geno[[j]]$data[i,] <- Sb.deduped$geno[[j]]$data[which(Sb.deduped$pheno$Ind == permdad)[1],]
			perm.Gmat$geno[[j]]$data[i,] <- Sb.deduped$geno[[j]]$data[which(Sb.deduped$pheno$Ind == permmom)[1],]
		}
	}
		
	# perform the joint scan
	G2results <- G2.scan(perm.Sb, scan.holder)
	G3results <- G3.scan(perm.Gmat, perm.Gpat, scan.holder)
	Joint.results <- G2results+G3results
	names(Joint.results)[3:6] <- c("JointBin", "JointBinNull", "JointBC", "JointBCNull")
	Joint.results[,7:8] <- NA
	Joint.results[,c(7,8)] <-  c(Joint.results$JointBin + Joint.results$JointBC, Joint.results$JointBinNull + Joint.results$JointBCNull)
	
		# add the maximum likelihoods to perm.results
	perm.results[m,1:6] <- apply(as.matrix(Joint.results[Joint.results $chr != "X",3:8]), 2, max)
	perm.results[m,7:12] <- apply(as.matrix(Joint.results[Joint.results $chr == "X",3:8]), 2, max)
	perm.out <- data.frame(perm.results)
	names(perm.out)  <- c("JointBin", "JointBinNull", "JointBC", "JointBCNull", "JointCombo","JointComboNull", "JointBinX", "JointBinNullX", "JointBCX", "JointBCNullX", "JointComboX","JointComboNullX")
	print(m); print(date());
}

rm(perm.results)
perm.out[,13:18] <- NA

perm.out[,13:18] <- cbind(2*(perm.out$JointBin - perm.out$JointBinNull), 2*(perm.out$JointBinX - perm.out$JointBinNullX), 2*(perm.out$JointBC - perm.out$JointBCNull), 2*(perm.out$JointBCX - perm.out$JointBCNullX), 2*(perm.out$JointCombo - perm.out$JointComboNull), 2*(perm.out$JointComboX - perm.out$JointComboNullX))
names(perm.out)[13:18] <- c("BinAutChisq", "BinXChisq", "BCAutChisq", "BCXChisq", "ComboAutChisq","ComboXChisq")

apply(perm.out[,c(13:18)], 2, function(x){quantile(x,0.95)})
#  BinAutChisq     BinXChisq    BCAutChisq      BCXChisq ComboAutChisq   ComboXChisq 
#     12.54124      11.35325      14.50577      11.05000      19.17413      17.17872 


perm.out -> ChNumJointScan2Perms
save(ChNumJointScan2Perms, file = "ChNumJointScan2.Perms.R")
rm(perm.out)


##############################################################################
# Third Forward Search
# The G3 scans do not include random effect of G2 father to avoid singular fit		
		
	G2chlenQ8.231 <- Sb.deduped$geno[[8]]$data[,"SNP231"] -1
	G3chlenQ8.231 <- MatGeno$geno[[8]]$data[,"SNP231"] + PatGeno$geno[[8]]$data[,"SNP231"] -2

	G2chlenQ3.497 <- Sb.deduped$geno[[3]]$data[,"SNP497"] -1
	G3chlenQ3.497 <- MatGeno$geno[[3]]$data[,"SNP497"] + PatGeno$geno[[3]]$data[,"SNP497"] -2

G2.scan <- function(crossfile, scan.object){
		G2sex <- crossfile$pheno$Sex
		binpheno <- crossfile$pheno$G2.chnum.bin.pheno
		bcpheno <- crossfile$pheno$G2bc.chnum
		binAutoNull <- logLik(glm(binpheno ~ G2chlenQ8.231 + G2chlenQ3.497, family = "binomial"))
		binXNull <- logLik(glm(binpheno ~ G2sex + G2chlenQ8.231 + G2chlenQ3.497, family = "binomial"))
		bcAutoNull <- logLik(lm(bcpheno ~ G2chlenQ8.231 + G2chlenQ3.497))	
		bcXNull <- logLik(lm(bcpheno ~ G2sex + G2chlenQ8.231 + G2chlenQ3.497))	
				
		for(j in 1: 10){		#autosomes only
		chr <- names(nmar(crossfile))[j]

				LL.G2chnumbin <- NULL
				LL.G2bcchnum <- NULL	
					
		for(i in 1:(nmar(crossfile)[chr])){ 	
										
				qtlG2 <- crossfile$geno[[chr]]$data[,i] -1
				LL.G2chnumbin[i] <- logLik(glm(binpheno ~ qtlG2 + G2chlenQ8.231 + G2chlenQ3.497, family = "binomial" ))		
				LL.G2bcchnum[i] <- logLik(lm(bcpheno ~ qtlG2 + G2chlenQ8.231 + G2chlenQ3.497))	
					}					
	
	scan.object[which(scan.object$chr == chr),3] <- LL.G2chnumbin
	scan.object[which(scan.object$chr == chr),4] <- binAutoNull
	scan.object[which(scan.object$chr == chr),5] <- LL.G2bcchnum
	scan.object[which(scan.object$chr == chr),6] <- bcAutoNull
	
				
	}
			for(j in 11){		# X only
			chr <- names(nmar(crossfile))[j]
				LL.G2chnumbin <- NULL
				LL.G2bcchnum <- NULL	
						
			for(i in 1:(nmar(crossfile)[chr])){ 	
										
				qtlG2 <- crossfile$geno[[chr]]$data[,i] -1
				LL.G2chnumbin[i] <- logLik(glm(binpheno ~ qtlG2*G2sex + G2chlenQ8.231 + G2chlenQ3.497, family = "binomial" ))		
				LL.G2bcchnum[i] <- logLik(lm(bcpheno ~ qtlG2*G2sex + G2chlenQ8.231 + G2chlenQ3.497))	

		}					
	
	scan.object[which(scan.object$chr == chr),3] <- LL.G2chnumbin
	scan.object[which(scan.object$chr == chr),4] <- binXNull
	scan.object[which(scan.object$chr == chr),5] <- LL.G2bcchnum
	scan.object[which(scan.object$chr == chr),6] <- bcXNull
	}

	names(scan.object)[c(3:6)] <- c("LL.G2ChNumBin", "LL.G2ChNumBinNull", "LL.G2ChNum", "LL.G2ChNumNull")
	return(scan.object)
}

ObsG2ChNumScan <- G2.scan(Sb.deduped, scan.holder)

G3.scan <- function(geno.mat, geno.pat, scan.object) {
	dadID <- geno.mat$pheno$Dad_G2_Name
	momID <-  geno.mat$pheno$Mom_G2_Name
		
	binpheno <- geno.mat$pheno$chnum.bin.pheno
	bcpheno <- geno.mat$pheno$bc.chnum0
	bin.null <- logLik(glmer(binpheno ~ G3chlenQ8.231 + G3chlenQ3.497 + (1|momID), family = "binomial"))[1]
	bc.null <- 	logLik(lmer(bcpheno ~ G3chlenQ8.231 + G3chlenQ3.497 + (1|dadID) + (1|momID), REML = F))[1]
	
		for(j in 1: 10){	# This function is only for the autosomes
		chr <- names(nmar(geno.mat))[j]
		bin.summed <- NULL
		bc.summed <- NULL
			for(i in 1:(nmar(geno.mat)[chr])){ 							
				qtlAll <- geno.mat$geno[[chr]]$data[,i] + geno.pat$geno[[chr]]$data[,i] - 2 				

				bin.summed[i] <- logLik(glmer(binpheno ~ qtlAll + G3chlenQ8.231 + G3chlenQ3.497 + (1|momID), family = "binomial"))[1]
				bc.summed[i] <- logLik(lmer(bcpheno ~ qtlAll + G3chlenQ8.231 + G3chlenQ3.497 + (1|dadID) + (1|momID), REML = F))[1]


		}				
	scan.object[which(scan.object$chr == chr),3] <- bin.summed
	scan.object[which(scan.object$chr == chr),4] <- bin.null
	scan.object[which(scan.object$chr == chr),5] <- bc.summed
	scan.object[which(scan.object$chr == chr),6] <- bc.null
	}
			for(j in 11){	# X only
		chr <- names(nmar(geno.mat))[j]
		bin.summed <- NULL
		bc.summed <- NULL

			for(i in 1:(nmar(geno.mat)[chr])){ 													
				qtlMom <- geno.mat$geno[[chr]]$data[,i] -1 # Count of lecithotroph alleles: 0, or 1.
				qtlDad <- geno.pat$geno[[chr]]$data[,i] -1 # Count of lecithotroph alleles: 0, or 1.
				qtlAll <- (1/4)*qtlDad + (3/8)*qtlMom # X-specific G3 additive effect parameterization
				bin.summed[i] <- logLik(glmer(binpheno ~ qtlAll + G3chlenQ8.231 + G3chlenQ3.497 + (1|momID), family = "binomial"))[1]
				bc.summed[i] <- logLik(lmer(bcpheno ~ qtlAll + G3chlenQ8.231 + G3chlenQ3.497 + (1|dadID) + (1|momID), REML = F))[1]

		}				
	scan.object[which(scan.object$chr == chr),3] <- bin.summed
	scan.object[which(scan.object$chr == chr),4] <- bin.null
	scan.object[which(scan.object$chr == chr),5] <- bc.summed
	scan.object[which(scan.object$chr == chr),6] <- bc.null
	}
	names(scan.object)[c(3:6)] <- c("LL.G3Bin", "LL.G3BinNull", "LL.G3ChNum", "LL.G3ChNumNull")
		
	return(scan.object)
}

ObsG3ChNumScan <- G3.scan(MatGeno,PatGeno, scan.holder)


# Joint Scan
ObsJointChNumScan <- ObsG2ChNumScan + ObsG3ChNumScan
names(ObsJointChNumScan)[c(3:6)] <- c("JointBin", "JointBinNull", "JointBC", "JointBCNull")
ObsJointChNumScan[,7:11] <- NA
ObsJointChNumScan[,c(7,8)] <-  c(ObsJointChNumScan$JointBin + ObsJointChNumScan$JointBC, ObsJointChNumScan$JointBinNull + ObsJointChNumScan$JointBCNull)
ObsJointChNumScan[,c(9:11)] <- c(2*(ObsJointChNumScan[,3] - ObsJointChNumScan[,4]), 2*(ObsJointChNumScan[,5] - ObsJointChNumScan[,6]), 2*(ObsJointChNumScan[,7] - ObsJointChNumScan[,8]))
names(ObsJointChNumScan)[c(7:11)] <- c("JointCombo", "JointComboNull", "BinChisq", "BCChisq", "ComboChisq")


# Top QTL peak:
ObsJointChNumScan[which(ObsJointChNumScan $ComboChisq == max(ObsJointChNumScan $ComboChisq)),]
#       chr      pos  JointBin JointBinNull   JointBC JointBCNull JointCombo JointComboNull
#SNP537   5 18.12664 -1908.271    -1910.682 -4378.512    -4385.14  -6286.783      -6295.822
#       BinChisq  BCChisq ComboChisq
#SNP537 4.823268 13.25529   18.07856

############
# A loop to perform the permutations
n.perms = 500
perm.results <- matrix(nrow = n.perms, ncol = 12)
date();
for(m in 1:n.perms){
	
		# Permute the G2 males
	male.perm <- data.frame(levels(factor(Sb.deduped$pheno$Ind[Sb.deduped$pheno$Sex == "M"])), levels(factor(Sb.deduped$pheno$Ind[Sb.deduped$pheno$Sex == "M"])))
	
	# Permutations are stratified by the F1 mother of the F2 fathers, i.e., 1A, 3C, 6F, 8H.
	male.perm[1:12,2] <- sample(male.perm[1:12,1])			
	male.perm[13:27,2] <- sample(male.perm[13:27,1])	
	male.perm[28:36,2] <- sample(male.perm[28:36,1])	
	male.perm[37:58,2] <- sample(male.perm[37:58,1])	

	perm.Sb <- Sb.deduped
	
	for(i in 1:dim(male.perm)[1]){
		for(ch in 1:11){
			perm.Sb$geno[[ch]]$data[i,] <- Sb.deduped$geno[[ch]]$data[which(Sb.deduped$pheno$Ind == male.perm[i,2])   ,]
		}
	}
		
	# Permute the G2 females
	female.perm <-  data.frame(levels(factor(Sb.deduped$pheno$Ind[Sb.deduped$pheno$Sex == "F"])), levels(factor(Sb.deduped$pheno$Ind[Sb.deduped$pheno$Sex == "F"])))

	# Permutations are stratified by the F1 mother of the F2 mothers, i.e., 1A, 3C, 6F, 8H.
	female.perm[1:39,2] <- sample(female.perm[1:39,1])	
	female.perm[40:98,2] <- sample(female.perm[40:98,1])		
	female.perm[99:126,2] <- sample(female.perm[99:126,1])	
	female.perm[127:183,2] <- sample(female.perm[127:183,1])	

	for(i in 1:dim(female.perm)[1]){
		for(ch in 1:11){
			perm.Sb$geno[[ch]]$data[i,] <- Sb.deduped$geno[[ch]]$data[which(Sb.deduped$pheno$Ind == female.perm[i,2])   ,]
		}
	}

	#subset down to the right number for the G3 data
	male.permG3 <- male.perm[which(male.perm[,1] %in% MatGeno$pheno$Dad_G2_Name),]
	female.permG3 <- female.perm[which(female.perm[,1] %in% MatGeno$pheno$Mom_G2_Name),]
	
	perm.Gmat <- MatGeno
	perm.Gpat <- PatGeno
	
	for(i in 1:dim(MatGeno$pheno)[1]){
		permdad <- male.permG3[which(male.permG3[,1] == MatGeno$pheno$Dad_G2_Name[i]),2]
		permmom <- female.permG3[which(female.permG3[,1] == MatGeno$pheno$Mom_G2_Name[i]),2]
		for(j in 1:10){
			perm.Gpat$geno[[j]]$data[i,] <- Sb.deduped$geno[[j]]$data[which(Sb.deduped$pheno$Ind == permdad)[1],]
			perm.Gmat$geno[[j]]$data[i,] <- Sb.deduped$geno[[j]]$data[which(Sb.deduped$pheno$Ind == permmom)[1],]
		}
	}
		
	# perform the joint scan
	G2results <- G2.scan(perm.Sb, scan.holder)
	G3results <- G3.scan(perm.Gmat, perm.Gpat, scan.holder)
	Joint.results <- G2results+G3results
	names(Joint.results)[3:6] <- c("JointBin", "JointBinNull", "JointBC", "JointBCNull")
	Joint.results[,7:8] <- NA
	Joint.results[,c(7,8)] <-  c(Joint.results$JointBin + Joint.results$JointBC, Joint.results$JointBinNull + Joint.results$JointBCNull)
	
		# add the maximum likelihoods to perm.results
	perm.results[m,1:6] <- apply(as.matrix(Joint.results[Joint.results $chr != "X",3:8]), 2, max)
	perm.results[m,7:12] <- apply(as.matrix(Joint.results[Joint.results $chr == "X",3:8]), 2, max)
	perm.out <- data.frame(perm.results)
	names(perm.out)  <- c("JointBin", "JointBinNull", "JointBC", "JointBCNull", "JointCombo","JointComboNull", "JointBinX", "JointBinNullX", "JointBCX", "JointBCNullX", "JointComboX","JointComboNullX")
	print(m); print(date());
}

rm(perm.results)
perm.out[,13:18] <- NA

perm.out[,13:18] <- cbind(2*(perm.out$JointBin - perm.out$JointBinNull), 2*(perm.out$JointBinX - perm.out$JointBinNullX), 2*(perm.out$JointBC - perm.out$JointBCNull), 2*(perm.out$JointBCX - perm.out$JointBCNullX), 2*(perm.out$JointCombo - perm.out$JointComboNull), 2*(perm.out$JointComboX - perm.out$JointComboNullX))
names(perm.out)[13:18] <- c("BinAutChisq", "BinXChisq", "BCAutChisq", "BCXChisq", "ComboAutChisq","ComboXChisq")

apply(perm.out[,c(13:18)], 2, function(x){quantile(x,0.95)})
#  BinAutChisq     BinXChisq    BCAutChisq      BCXChisq ComboAutChisq   ComboXChisq 
#    12.015250     12.334409     14.304296      9.801793     20.256244     15.955419 


perm.out -> ChNumJointScan3Perms
save(ChNumJointScan3Perms, file = "ChNumJointScan3.Perms.R")
rm(perm.out)
```
