## Supplementary File 1. Data and Code for "Baby makes three: maternal, paternal, and zygotic genetic effects shape larval phenotypic evolution": JointCirriScan.R

```
# Combine likelihoods from G2s and G3s

library(qtl)
library(lme4)

load(file = "G3AnalysisObjects.R")
# [1] "dups"       "MatGeno"    "PatGeno"    "Sb.deduped"


scan.holder <- scanone(Sb.deduped, pheno = "nchaetae")
scan.holder$lod <- 0
scan.holder[,4] <- 0

G2.scan <- function(crossfile, scan.object){
		G2sex <- crossfile$pheno$Sex
		cirr.bin <- crossfile$pheno$cirri.binary
		AutoNull <- logLik(glm(cirr.bin ~ 1,  family = "binomial" ))[1]
		XNull <- logLik(glm(cirr.bin ~ G2sex,  family = "binomial" ))[1]
		for(j in 1: 10){		#autosomes only
			chr <- names(nmar(crossfile))[j]
			LL.G2cirri <- NULL
					
		for(i in 1:(nmar(crossfile)[chr])){ 	
										
				qtlG2 <- crossfile $geno[[chr]]$data[,i] -1
				LL.G2cirri[i] <- logLik(glm(cirr.bin ~ qtlG2, family = "binomial" ))	[1]	
		}					
	scan.object[which(scan.object$chr == chr),3] <- LL.G2cirri
	scan.object[which(scan.object$chr == chr),4] <- AutoNull
	}
			for(j in 11){		# X only
			chr <- names(nmar(crossfile))[j]
			LL.G2cirri <- NULL
						
			for(i in 1:(nmar(crossfile)[chr])){ 	
										
				qtlG2 <- crossfile $geno[[chr]]$data[,i] -1
				LL.G2cirri[i] <- logLik(glm(cirr.bin ~ qtlG2*G2sex, family = "binomial"))[1]		
		}					
	scan.object[which(scan.object$chr == chr),3] <- LL.G2cirri
	scan.object[which(scan.object$chr == chr),4] <- XNull
	}

	names(scan.object)[3:4] <- c("LL.G2cirri", "LL.G2Null")
	return(scan.object)
}

ObsG2CirriScan <- G2.scan(Sb.deduped, scan.holder)

G3.scan <- function(geno.mat, geno.pat, scan.object) {
	dadID <- geno.mat$pheno$Dad_G2_Name
	momID <-  geno.mat$pheno$Mom_G2_Name
	cirr.bin <- geno.mat$pheno$cirri
	cirri.G3.null <- logLik(glmer(cirr.bin ~ 1 + (1|dadID) + (1|momID), family= "binomial"))[1]

		for(j in 1: 10){	# This function is only for the autosomes
		chr <- names(nmar(geno.mat))[j]
		cirri.summed <- NULL

			for(i in 1:(nmar(geno.mat)[chr])){ 							
				qtlAll <- geno.mat$geno[[chr]]$data[,i] + geno.pat$geno[[chr]]$data[,i] - 2 				
				cirri.summed[i] <- logLik(glmer(cirr.bin ~ qtlAll + (1|dadID) + (1|momID), family = "binomial"))[1]
		}				
	scan.object[which(scan.object$chr == chr),3] <- cirri.summed
	scan.object[which(scan.object$chr == chr),4] <- cirri.G3.null
	}
			for(j in 11){	# X only
		chr <- names(nmar(geno.mat))[j]
		cirri.summed <- NULL

			for(i in 1:(nmar(geno.mat)[chr])){ 													
				qtlMom <- geno.mat$geno[[chr]]$data[,i] -1 # Count of lecithotroph alleles: 0, or 1.
				qtlDad <- geno.pat$geno[[chr]]$data[,i] -1 # Count of lecithotroph alleles: 0, or 1.
				qtlAll <- (1/4)*qtlDad + (3/8)*qtlMom # X-specific G3 additive effect parameterization
				cirri.summed[i] <- logLik(glmer(cirr.bin ~ qtlAll + (1|dadID) + (1|momID), family = "binomial"))[1]
		}				
	scan.object[which(scan.object$chr == chr),3] <- cirri.summed
	scan.object[which(scan.object$chr == chr),4] <- cirri.G3.null

	}
	names(scan.object)[3:4] <- c("LL.G3cirri", "LL.G3Null")
		
	return(scan.object)
}

ObsG3CirriScan <- G3.scan(MatGeno,PatGeno, scan.holder)


# Joint Scan
ObsJointCirriScan <- ObsG2CirriScan + ObsG3CirriScan
ObsJointCirriScan[,5] <- NA
ObsJointCirriScan[,5] <-  2*(ObsJointCirriScan[,3] - ObsJointCirriScan[,4])
names(ObsJointCirriScan)[c(3,4,5)] <- c("JointCirri", "JointCirriNull", "CirriChisq")

# Top QTL peak:
ObsJointCirriScan[which(ObsJointCirriScan$CirriChisq == max(ObsJointCirriScan$CirriChisq)),]
#       chr      pos JointCirri JointCirriNull CirriChisq
#SNP574   5 3.679437  -2149.537      -2180.541   62.00907

###############
# A loop to perform the permutations
n.perms = 200
perm.results <- matrix(nrow = n.perms, ncol = 4)
date();
for(m in 1:n.perms){
	
		# Permute the G2 males
	male.perm <- data.frame(levels(factor(Sb.deduped$pheno$Ind[Sb.deduped$pheno$Sex == "M"])), levels(factor(Sb.deduped$pheno$Ind[Sb.deduped$pheno$Sex == "M"])))
	
	# Permutations are stratified by the F1 mother of the F2 fathers, i.e., 1A, 3C, 6F, 8H.
	male.perm[1:12,2] <- sample(male.perm[1:12,1])			
	male.perm[13:27,2] <- sample(male.perm[13:27,1])	
	male.perm[28:36,2] <- sample(male.perm[28:36,1])	
	male.perm[37:58,2] <- sample(male.perm[37:58,1])	

	perm.Sb <- Sb.deduped
	
	for(i in 1:dim(male.perm)[1]){
		for(ch in 1:11){
			perm.Sb$geno[[ch]]$data[i,] <- Sb.deduped$geno[[ch]]$data[which(Sb.deduped$pheno$Ind == male.perm[i,2])   ,]
		}
	}
		
	# Permute the G2 females
	female.perm <-  data.frame(levels(factor(Sb.deduped$pheno$Ind[Sb.deduped$pheno$Sex == "F"])), levels(factor(Sb.deduped$pheno$Ind[Sb.deduped$pheno$Sex == "F"])))

	# Permutations are stratified by the F1 mother of the F2 mothers, i.e., 1A, 3C, 6F, 8H.
	female.perm[1:39,2] <- sample(female.perm[1:39,1])	
	female.perm[40:98,2] <- sample(female.perm[40:98,1])		
	female.perm[99:126,2] <- sample(female.perm[99:126,1])	
	female.perm[127:183,2] <- sample(female.perm[127:183,1])	

	for(i in 1:dim(female.perm)[1]){
		for(ch in 1:11){
			perm.Sb$geno[[ch]]$data[i,] <- Sb.deduped$geno[[ch]]$data[which(Sb.deduped$pheno$Ind == female.perm[i,2])   ,]
		}
	}

	#subset down to the right number for the G3 data
	male.permG3 <- male.perm[which(male.perm[,1] %in% MatGeno$pheno$Dad_G2_Name),]
	female.permG3 <- female.perm[which(female.perm[,1] %in% MatGeno$pheno$Mom_G2_Name),]
	
	perm.Gmat <- MatGeno
	perm.Gpat <- PatGeno
	
	for(i in 1:dim(MatGeno$pheno)[1]){
		permdad <- male.permG3[which(male.permG3[,1] == MatGeno$pheno$Dad_G2_Name[i]),2]
		permmom <- female.permG3[which(female.permG3[,1] == MatGeno$pheno$Mom_G2_Name[i]),2]
		for(j in 1:10){
			perm.Gpat$geno[[j]]$data[i,] <- Sb.deduped$geno[[j]]$data[which(Sb.deduped$pheno$Ind == permdad)[1],]
			perm.Gmat$geno[[j]]$data[i,] <- Sb.deduped$geno[[j]]$data[which(Sb.deduped$pheno$Ind == permmom)[1],]
		}
	}
		
	# perform the joint scan
G2results <- G2.scan(perm.Sb, scan.holder)
G3results <- G3.scan(perm.Gmat, perm.Gpat, scan.holder)
Joint.results <- G2results+G3results
names(Joint.results)[3:4] <- c("Joint.Cirri", "Joint.Cirri.Null") 

	# add the maximum likelihoods to perm.results
perm.results[m,1:2] <- apply(as.matrix(Joint.results[Joint.results $chr != "X",3:4]), 2, max)
perm.results[m,3:4] <- apply(as.matrix(Joint.results[Joint.results $chr == "X",3:4]), 2, max)
perm.out <- data.frame(perm.results)
names(perm.out)  <- c("Joint.Cirri", "Joint.Cirri.Null", "Joint.Cirri.X", "Joint.Cirri.X.Null")
print(m); print(date());
}

rm(perm.results)
perm.out[,5:6] <- NA

perm.out[,5:6] <- cbind(2*(perm.out$Joint.Cirri - perm.out$Joint.Cirri.Null), 2*(perm.out$Joint.Cirri.X - perm.out$Joint.Cirri.X.Null))
names(perm.out)[5:6] <- c("JointAutChisq", "JointXChisq")

apply(perm.out[,c(5,6)], 2, function(x){quantile(x,0.95)})
#JointAutChisq   JointXChisq 
#     14.42411      12.29986 

perm.out -> CirriJointScan1Perms
save(CirriJointScan1Perms, file = "CirriJointScan1.200Perms.R")
rm(perm.out)

##############################################################################
# Second Forward Search
# The G3 scans do not include random effect of G2 father to avoid singular fit		
		
	G2chlenQ5.574 <- Sb.deduped$geno[[5]]$data[,"SNP574"] -1
	G3chlenQ5.574 <- MatGeno$geno[[5]]$data[,"SNP574"] + PatGeno$geno[[5]]$data[,"SNP574"] -2

G2.scan <- function(crossfile, scan.object){
		G2sex <- crossfile$pheno$Sex
		cirr.bin <- crossfile$pheno$cirri.binary
		AutoNull <- logLik(glm(cirr.bin ~ G2chlenQ5.574,  family = "binomial" ))[1]
		XNull <- logLik(glm(cirr.bin ~ G2sex + G2chlenQ5.574,  family = "binomial" ))[1]
		for(j in 1: 10){		#autosomes only
			chr <- names(nmar(crossfile))[j]
			LL.G2cirri <- NULL
					
		for(i in 1:(nmar(crossfile)[chr])){ 	
										
				qtlG2 <- crossfile $geno[[chr]]$data[,i] -1
				LL.G2cirri[i] <- logLik(glm(cirr.bin ~ qtlG2 + G2chlenQ5.574, family = "binomial" ))	[1]	
		}					
	scan.object[which(scan.object$chr == chr),3] <- LL.G2cirri
	scan.object[which(scan.object$chr == chr),4] <- AutoNull
	}
			for(j in 11){		# X only
			chr <- names(nmar(crossfile))[j]
			LL.G2cirri <- NULL
						
			for(i in 1:(nmar(crossfile)[chr])){ 	
										
				qtlG2 <- crossfile $geno[[chr]]$data[,i] -1
				LL.G2cirri[i] <- logLik(glm(cirr.bin ~ qtlG2*G2sex + G2chlenQ5.574, family = "binomial"))[1]		
		}					
	scan.object[which(scan.object$chr == chr),3] <- LL.G2cirri
	scan.object[which(scan.object$chr == chr),4] <- XNull
	}

	names(scan.object)[3:4] <- c("LL.G2cirri", "LL.G2Null")
	return(scan.object)
}

ObsG2CirriScan <- G2.scan(Sb.deduped, scan.holder)

G3.scan <- function(geno.mat, geno.pat, scan.object) {
	dadID <- geno.mat$pheno$Dad_G2_Name
	momID <-  geno.mat$pheno$Mom_G2_Name
	cirr.bin <- geno.mat$pheno$cirri
	cirri.G3.null <- logLik(glmer(cirr.bin ~ G3chlenQ5.574 + (1|momID), family= "binomial"))[1]

		for(j in 1: 10){	# This function is only for the autosomes
		chr <- names(nmar(geno.mat))[j]
		cirri.summed <- NULL

			for(i in 1:(nmar(geno.mat)[chr])){ 							
				qtlAll <- geno.mat$geno[[chr]]$data[,i] + geno.pat$geno[[chr]]$data[,i] - 2 				
				cirri.summed[i] <- logLik(glmer(cirr.bin ~ qtlAll + G3chlenQ5.574 + (1|momID), family = "binomial"))[1]
		}				
	scan.object[which(scan.object$chr == chr),3] <- cirri.summed
	scan.object[which(scan.object$chr == chr),4] <- cirri.G3.null
	}
			for(j in 11){	# X only
		chr <- names(nmar(geno.mat))[j]
		cirri.summed <- NULL

			for(i in 1:(nmar(geno.mat)[chr])){ 													
				qtlMom <- geno.mat$geno[[chr]]$data[,i] -1 # Count of lecithotroph alleles: 0, or 1.
				qtlDad <- geno.pat$geno[[chr]]$data[,i] -1 # Count of lecithotroph alleles: 0, or 1.
				qtlAll <- (1/4)*qtlDad + (3/8)*qtlMom # X-specific G3 additive effect parameterization
				cirri.summed[i] <- logLik(glmer(cirr.bin ~ qtlAll + G3chlenQ5.574 + (1|momID), family = "binomial"))[1]
		}				
	scan.object[which(scan.object$chr == chr),3] <- cirri.summed
	scan.object[which(scan.object$chr == chr),4] <- cirri.G3.null

	}
	names(scan.object)[3:4] <- c("LL.G3cirri", "LL.G3Null")
		
	return(scan.object)
}

ObsG3CirriScan <- G3.scan(MatGeno,PatGeno, scan.holder)

# Joint Scan
ObsJointCirriScan <- ObsG2CirriScan + ObsG3CirriScan
ObsJointCirriScan[,5] <- NA
ObsJointCirriScan[,5] <-  2*(ObsJointCirriScan[,3] - ObsJointCirriScan[,4])
names(ObsJointCirriScan)[c(3,4,5)] <- c("JointCirri", "JointCirriNull", "CirriChisq")

# Top QTL peak:
ObsJointCirriScan[which(ObsJointCirriScan$CirriChisq == max(ObsJointCirriScan$CirriChisq)),]
#       chr      pos JointCirri JointCirriNull CirriChisq
#SNP894   2 27.83767  -2144.574      -2149.536   9.925296

###############
# A loop to perform the permutations

n.perms = 100
perm.results <- matrix(nrow = n.perms, ncol = 4)
date();
for(m in 1:n.perms){
	
		# Permute the G2 males
	male.perm <- data.frame(levels(factor(Sb.deduped$pheno$Ind[Sb.deduped$pheno$Sex == "M"])), levels(factor(Sb.deduped$pheno$Ind[Sb.deduped$pheno$Sex == "M"])))
	
	# Permutations are stratified by the F1 mother of the F2 fathers, i.e., 1A, 3C, 6F, 8H.
	male.perm[1:12,2] <- sample(male.perm[1:12,1])			
	male.perm[13:27,2] <- sample(male.perm[13:27,1])	
	male.perm[28:36,2] <- sample(male.perm[28:36,1])	
	male.perm[37:58,2] <- sample(male.perm[37:58,1])	

	perm.Sb <- Sb.deduped
	
	for(i in 1:dim(male.perm)[1]){
		for(ch in 1:11){
			perm.Sb$geno[[ch]]$data[i,] <- Sb.deduped$geno[[ch]]$data[which(Sb.deduped$pheno$Ind == male.perm[i,2])   ,]
		}
	}
		
	# Permute the G2 females
	female.perm <-  data.frame(levels(factor(Sb.deduped$pheno$Ind[Sb.deduped$pheno$Sex == "F"])), levels(factor(Sb.deduped$pheno$Ind[Sb.deduped$pheno$Sex == "F"])))

	# Permutations are stratified by the F1 mother of the F2 mothers, i.e., 1A, 3C, 6F, 8H.
	female.perm[1:39,2] <- sample(female.perm[1:39,1])	
	female.perm[40:98,2] <- sample(female.perm[40:98,1])		
	female.perm[99:126,2] <- sample(female.perm[99:126,1])	
	female.perm[127:183,2] <- sample(female.perm[127:183,1])	

	for(i in 1:dim(female.perm)[1]){
		for(ch in 1:11){
			perm.Sb$geno[[ch]]$data[i,] <- Sb.deduped$geno[[ch]]$data[which(Sb.deduped$pheno$Ind == female.perm[i,2])   ,]
		}
	}

	#subset down to the right number for the G3 data
	male.permG3 <- male.perm[which(male.perm[,1] %in% MatGeno$pheno$Dad_G2_Name),]
	female.permG3 <- female.perm[which(female.perm[,1] %in% MatGeno$pheno$Mom_G2_Name),]
	
	perm.Gmat <- MatGeno
	perm.Gpat <- PatGeno
	
	for(i in 1:dim(MatGeno$pheno)[1]){
		permdad <- male.permG3[which(male.permG3[,1] == MatGeno$pheno$Dad_G2_Name[i]),2]
		permmom <- female.permG3[which(female.permG3[,1] == MatGeno$pheno$Mom_G2_Name[i]),2]
		for(j in 1:10){
			perm.Gpat$geno[[j]]$data[i,] <- Sb.deduped$geno[[j]]$data[which(Sb.deduped$pheno$Ind == permdad)[1],]
			perm.Gmat$geno[[j]]$data[i,] <- Sb.deduped$geno[[j]]$data[which(Sb.deduped$pheno$Ind == permmom)[1],]
		}
	}
		
	# perform the joint scan
G2results <- G2.scan(perm.Sb, scan.holder)
G3results <- G3.scan(perm.Gmat, perm.Gpat, scan.holder)
Joint.results <- G2results+G3results
names(Joint.results)[3:4] <- c("Joint.Cirri", "Joint.Cirri.Null") 

	# add the maximum likelihoods to perm.results
perm.results[m,1:2] <- apply(as.matrix(Joint.results[Joint.results $chr != "X",3:4]), 2, max)
perm.results[m,3:4] <- apply(as.matrix(Joint.results[Joint.results $chr == "X",3:4]), 2, max)
perm.out <- data.frame(perm.results)
names(perm.out)  <- c("Joint.Cirri", "Joint.Cirri.Null", "Joint.Cirri.X", "Joint.Cirri.X.Null")
print(m); print(date());
}

rm(perm.results)
perm.out[,5:6] <- NA

perm.out[,5:6] <- cbind(2*(perm.out$Joint.Cirri - perm.out$Joint.Cirri.Null), 2*(perm.out$Joint.Cirri.X - perm.out$Joint.Cirri.X.Null))
names(perm.out)[5:6] <- c("JointAutChisq", "JointXChisq")

apply(perm.out[,c(5,6)], 2, function(x){quantile(x,0.95)})
#JointAutChisq   JointXChisq 
#     14.32619      15.03404 


perm.out -> CirriJointScan2Perms
save(CirriJointScan2Perms, file = "CirriJointScan2.100Perms.R")
rm(perm.out)
```
