## Supplementary File 1. Data and Code for "Baby makes three: maternal, paternal, and zygotic genetic effects shape larval phenotypic evolution": G2andG3DataHarmonization.R

```
# Harmonize the G2 and G3 datasets by picking out the minimal set of non-redundant markers.
# This script starts with the G3 phenotypes and G2 genotypes and ends with the input files for linkage mapping.

library(qtl)
library(lme4)


# load Rqtl cross objects from the supplement of Zakas et al. 2018
load("SbenedictiGeneticMaps.R")  # This is FileS1 from the 2018 paper.

# impute missing data on the X chromosome
sim.geno(Sb702, n.draws = 1, error.prob=0) -> Sb702
Sb702$geno[["X"]]$data <- Sb702$geno[["X"]]$draws[,,1]

# find redundant markers
dups <- findDupMarkers(Sb702, adjacent.only = T)
Sb.deduped <- drop.markers(Sb702, unlist(dups))

summary(Sb.deduped)
# 275 markers, with 16-32 per chromosome.

# One worm name is ambiguous and should get corrected here to match the G3 data:
Sb.deduped$pheno$Ind <- as.character(Sb.deduped$pheno$Ind)
Sb.deduped$pheno$Ind[which(Sb.deduped$pheno$Ind == "C73.1756")] <- "C73.17"
Sb.deduped$pheno$Ind <- factor(Sb.deduped$pheno$Ind)

rm(list = ls()[c(3,4,5)])

#save.image(file = "DedupedG2cross.275.R")
## Use this file for downstream analyses, to deal with the thing where findDupMarkers picks a random marker name for each marker group.
load("DedupedG2cross.275.R")


#### NEXT STEP IS to get the X chromosome data into the G3 genotypes

G3pheno <- read.cross(file = "G3_pheno.csv", format = "csv") # This is a placeholder with columns for each of the 275 markers in the Sb.deduped G2 dataset.

G3DadGeno <- G3pheno
G3MomGeno <- G3pheno

for(i in 1:dim(G3pheno$pheno)[1]){
		G3dad <- G3pheno$pheno$Dad_G2_Name[i]
		G3mom <- G3pheno$pheno$Mom_G2_Name[i]

		for(j in 1:11){
			G3DadGeno$geno[[j]]$data[i,] <- Sb.deduped$geno[[j]]$data[which(Sb.deduped$pheno$Ind == as.character(G3dad)),]
			G3MomGeno$geno[[j]]$data[i,] <- Sb.deduped$geno[[j]]$data[which(Sb.deduped$pheno$Ind == as.character(G3mom)),]
		}
	}
write.cross(G3MomGeno, file = "G3MomGeno")
write.cross(G3DadGeno, file = "G3DadGeno")

MatGeno <- read.cross(file = "G3MomGeno.csv", format = "csv", geno = c("AA","AB","BB"))
PatGeno <- read.cross(file = "G3DadGeno.csv", format = "csv", geno = c("AA","AB","BB"))
# These are the input files for analysis of the G3 data: each G3 larva with the genotype of its mother or its father, at 275 markers that are non-redundant across the whole G2 dataset. 

save.image(file = "G3AnalysisObjects.R")
# [1] "dups"       "MatGeno"    "PatGeno"    "Sb.deduped"
```
